# Identification of differentially expressed genes and signaling pathways in gestational diabetes mellitus by integrated bioinformatics analysis

**DOI:** 10.1101/2021.11.24.469869

**Authors:** Basavaraj Vastrad, Chanabasayya Vastrad

## Abstract

Gestational diabetes mellitus (GDM) is a metabolic disorder during pregnancy. Numerous biomarkers have been identified that are linked with the occurrence and development of GDM. The aim of this investigation was to identify differentially expressed genes (DEGs) in GDM using a bioinformatics approach to elucidate their molecular pathogenesis. GDM associated expression profiling by high throughput sequencing dataset (GSE154377) was obtained from Gene Expression Omnibus (GEO) database including 28 normal pregnancy samples and 33 GDM samples. DEGs were identified using DESeq2. The gene ontology (GO) and REACTOME pathway enrichments of DEGs were performed by g:Profiler. Protein-protein interaction (PPI) networks were assembled with Cytoscape software and separated into modules using the PEWCC1 algorithm. MiRNA-hub gene regulatory network and TF-hub gene regulatory network were performed with the miRNet database and NetworkAnalyst database. Receiver Operating Characteristic (ROC) analyses was conducted to validate the hub genes. A total of 953 DEGs were identified, of which 478 DEGs were up regulated and 475 DEGs were down regulated. Furthermore, GO and REACTOME pathway enrichment analysis demonstrated that these DEGs were mainly enriched in multicellular organismal process, cell activation, formation of the cornified envelope and hemostasis. TRIM54, ELAVL2, PTN, KIT, BMPR1B, APP, SRC, ITGA4, RPA1 and ACTB were identified as key genes in the PPI network, miRNA-hub gene regulatory network and TF-hub gene regulatory network. TRIM54, ELAVL2, PTN, KIT, BMPR1B, APP, SRC, ITGA4, RPA1 and ACTB in GDM were validated using ROC analysis. This investigation provides further insights into the molecular pathogenesis of GDM, which might facilitate the diagnosis and treatment of GDM.

## Introduction

Gestational diabetes mellitus (GDM) has emerged as most common medical complications of pregnancy, but current treatments are still suboptimal [1]. The number of cases GDM is rising worldwide and it has become a key health concern [2]. There are several important risk factors for GDM, such as advanced maternal age [3], family history of diabetes [4], obesity [5], cigarette smoking [6], genetics factors [7], hypertension and type 2 diabetes [8], and cardiovascular diseases [9]. However, GDM is a complex disorder and its biology remains poorly understood. Therefore, mining key biomarkers linked with GDM occurrence and advancement is essential for GDM early diagnosis, prognosis and treatment.

The GDM is an extremely complicated pathophysiological process and controlled by various genes and signaling pathways. Genes such as PVT1 [10], IL-6, IL-10, and TNF-α [11], CAPN10 [12], ADCYAP1 [13] and ABHD5 [14] are associated with GDM. Signaling pathways such as PI3K/AKT signaling pathway [15], AMPK signaling pathway [16], vitamin D signaling pathways [17], TLR4/NF-kappaB Signaling Pathway [18] and AGEs-RAGE signaling pathway [19] are linked with GDM. Recently, next-generation sequencing (NGS) technology is extensively applied in clinical investigation, with the marked gain relying on its capacity to together determine the expression data of massive genes at a time [20]. The public database Gene Expression Omnibus (GEO) (http://www.ncbi.nlm.nih.gov/geo/) [21] are powerful tools used to screen the differentially expressed genes (DEGs) generated from NGS data corresponding to the progression of GDM [22]. Therefore, there are still many related genes to be identified, which will help us better understand the molecular pathogenesis of GDM and facilitate the discovery of new diagnostic biomarkers or therapeutic target.

In the current investigation, we tested integrated bioinformatics analysis for the identification of DEGs as potential biomarkers of GDM based on the expression profiling by high throughput sequencing from the GEO database. These analysis might provide possible perspectives for the progression and development of GDM as well as identification potential candidate genes for GDM diagnose or treatment.

## Materials and methods

### Data resources

The NGS data used in this investigation were acquired from GEO database, which is a public repository containing the expression profiling by high throughput sequencing data submitted by research institutions around the world. We chose the expression profiling by high throughput sequencing data GSE154377 [23] from the GEO database. The NGS data of GSE154377 was consisted with 28 normal pregnancy samples and 33 GDM samples based on the platform of GPL20301 Illumina HiSeq 4000 (Homo sapiens).

### Identification of DEGs

DESeq2 [24] packages of R Bioconductor (http://www.bioconductor.org/) was applied to significance analysis of DEGs between normal pregnancy samples and GDM samples. The log-fold change (FC) in expression and adjusted P-values (adj. P) were resolved. The criterion of statistical significance was adj. P value <0.05, |Log2 (FC)|> 1.73 for up regulated genes and |Log2 (FC)|> -0.69 for down regulated genes. A heatmap of the DEGs was produced using gplots in R software. All significant DEGs were shown in a volcano plot generated using ggplot2 in R software.

### Gene Ontology (GO) and REACTOME pathway enrichment analysis of DEGs

Gene Ontology (GO) (http://www.geneontology.org) [25] is a suitable technique to classify gene expression and its properties, including biological process (BP), cellular component (CC) and molecular function (MF), which provide extensive functional annotation tools for researchers to integrate key genes with a specific function. REACTOME (https://reactome.org/) [26] is an online pathway database that collects information on genomic, biochemical, and enzymatic pathways. The DEGs were mapped to the REACTOME pathway database and the significantly related pathways were screened. We could use g:Profiler (http://biit.cs.ut.ee/gprofiler/) [27] is an online functional annotation tool to visualize the DEGs enrichment of BP, MF, CC and pathways (P□<□0.05).

### Construction of the PPI network and module analysis

IID interactome (Integrated Interactions Database) (http://iid.ophid.utoronto.ca/) [28] was used to construct PPI network. The identified DEGs were input into IID interactome to unravel a potential PPI network. Afterwards, a PPI network was constructed and visualized by Cytoscape software (version 3.8.2, http://www.cytoscape.org/) [29]. Subsequently, the topological properties including node degree [30], betweenness centrality [31], stress centrality [32] and closeness centrality [33] of the DEGs in the PPI network were calculated by Network Analyzer plug-in of Cytoscape to further analyze the candidate hub genes from the PPI network. The PEWCC1 plug-in [34] was used to search sub-networks of the PPI network and the default parameters (Degree cutoff ≥ 10, node score cutoff ≥ 0.4, K-core ≥ 4, and max depth=100.) were set in the functional interface of Cytoscape software.

### MiRNA-hub gene regulatory network construction

The miRNA-hub gene interactions were predicted with miRNet database (https://www.mirnet.ca/) [35], which involves 14 predicted algorithms (TarBase, miRTarBase, miRecords, miRanda (S mansoni only), miR2Disease, HMDD, PhenomiR, SM2miR, PharmacomiR, EpimiR, starBase, TransmiR, ADmiRE, and TAM 2.0). The Cytoscape 3.8.2 software [29] was used to visualize the regulatory network.

### TF-hub gene regulatory network construction

The TF-hub gene interactions were predicted with NetworkAnalyst database (https://www.networkanalyst.ca/) [36], which involves 1 predicted algorithm (ChEA). The Cytoscape 3.8.2 software [29] was used to visualize the regulatory network.

### Receiver operating characteristic curve (ROC) analysis

pROC package in R statistical software [37] was used to operate ROC curve analysis and to resolve the specificity, sensitivity, positive and negative predictive values for all the feasible thresholds of the ROC curve. The value of the hub genes were predicted based on the ROC curve analysis.

## Results

### Identification of DEGs

DEGs were identified by DESeq2 in the R package. In total, 953 DEGs (478 up and 475 down regulated genes) were selected based on the criteria of adj. P value <0.05, |Log2 (FC)|> 1.73 for up regulated genes and |Log2 (FC)|> -0.69 for down regulated genes. Heatmap of DEGs is presented in Fig. 1, and volcano plot is presented in Fig. 2. The up and down regulated DEGs is listed in Table 1.

**Fig. 1.**
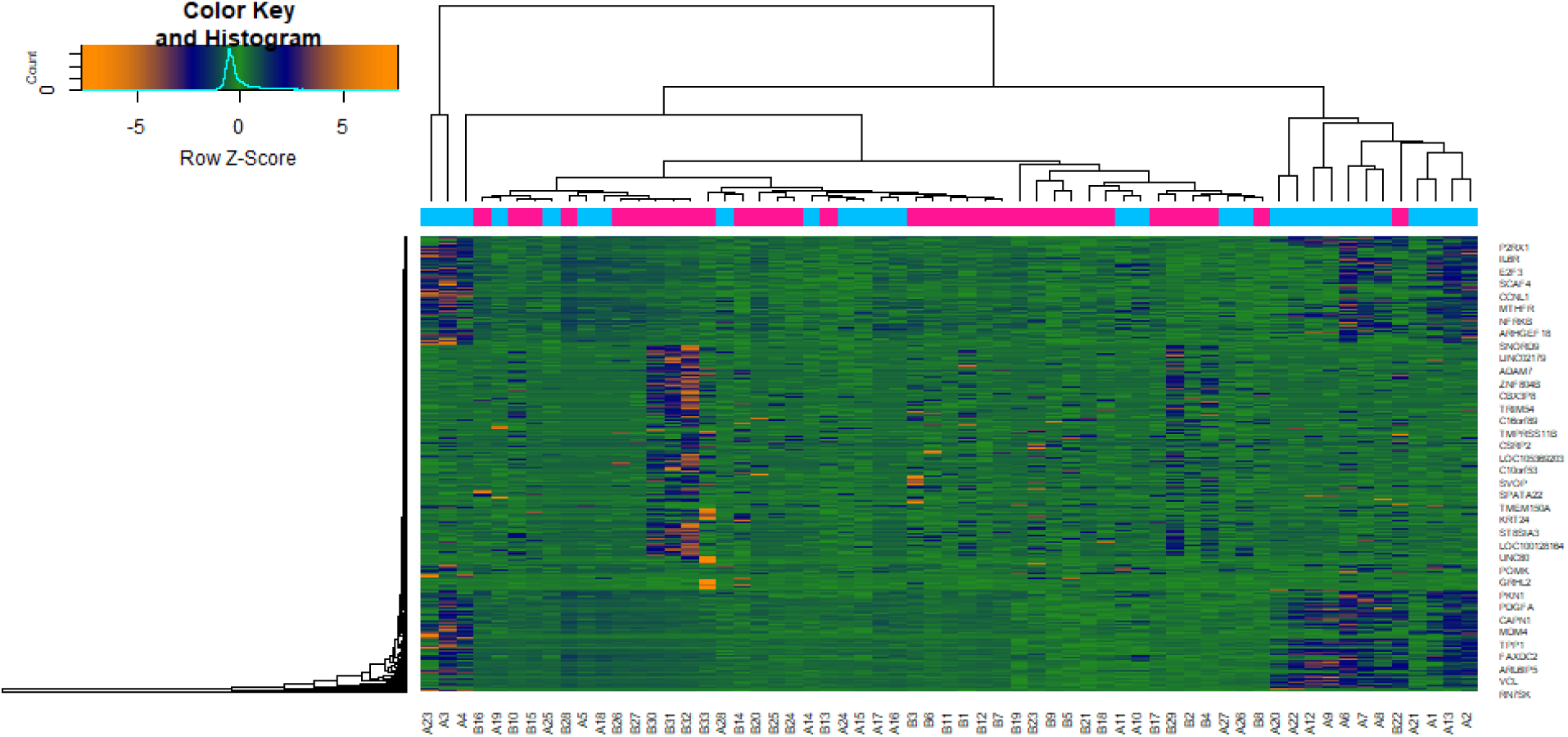
Heat map of differentially expressed genes. Legend on the top left indicate log fold change of genes. (A1 – A28= Normal Pregnancy Samples; B1 – B33= Gestational diabetes Samples)

**Fig. 2.**
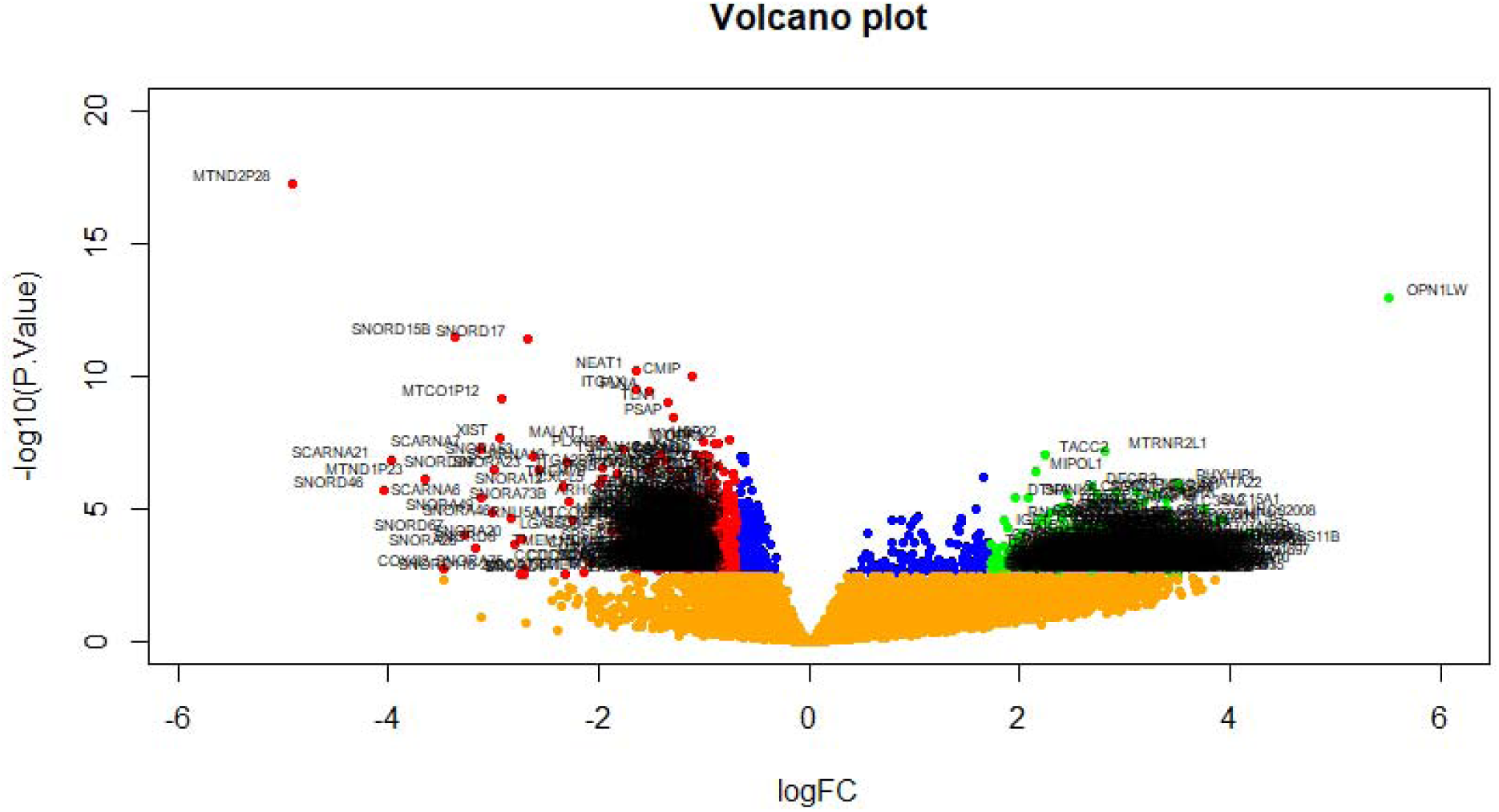
Volcano plot of differentially expressed genes. Genes with a significant change of more than two-fold were selected. Green dot represented up regulated significant genes and red dot represented down regulated significant genes.

**Table 1.**
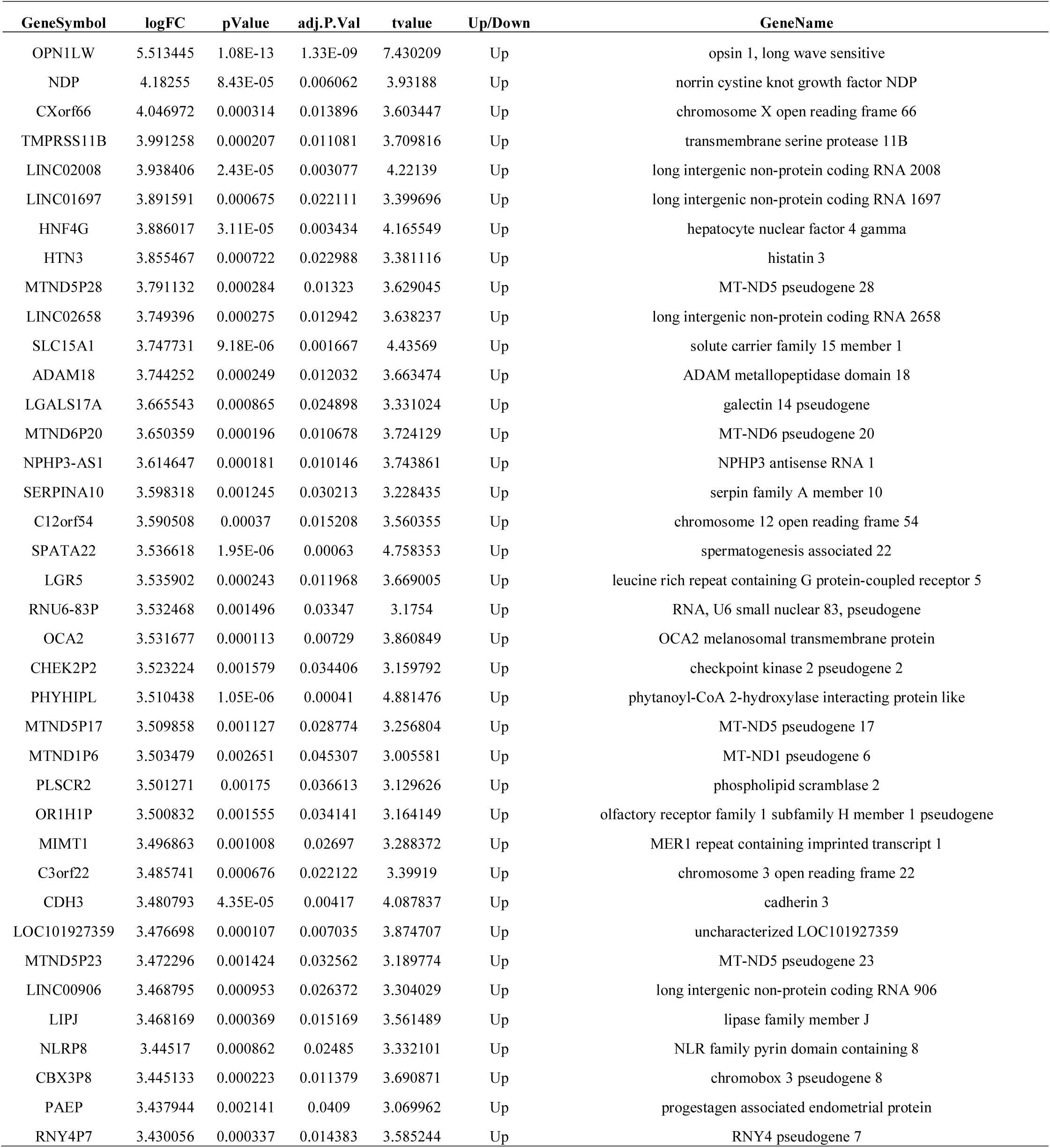
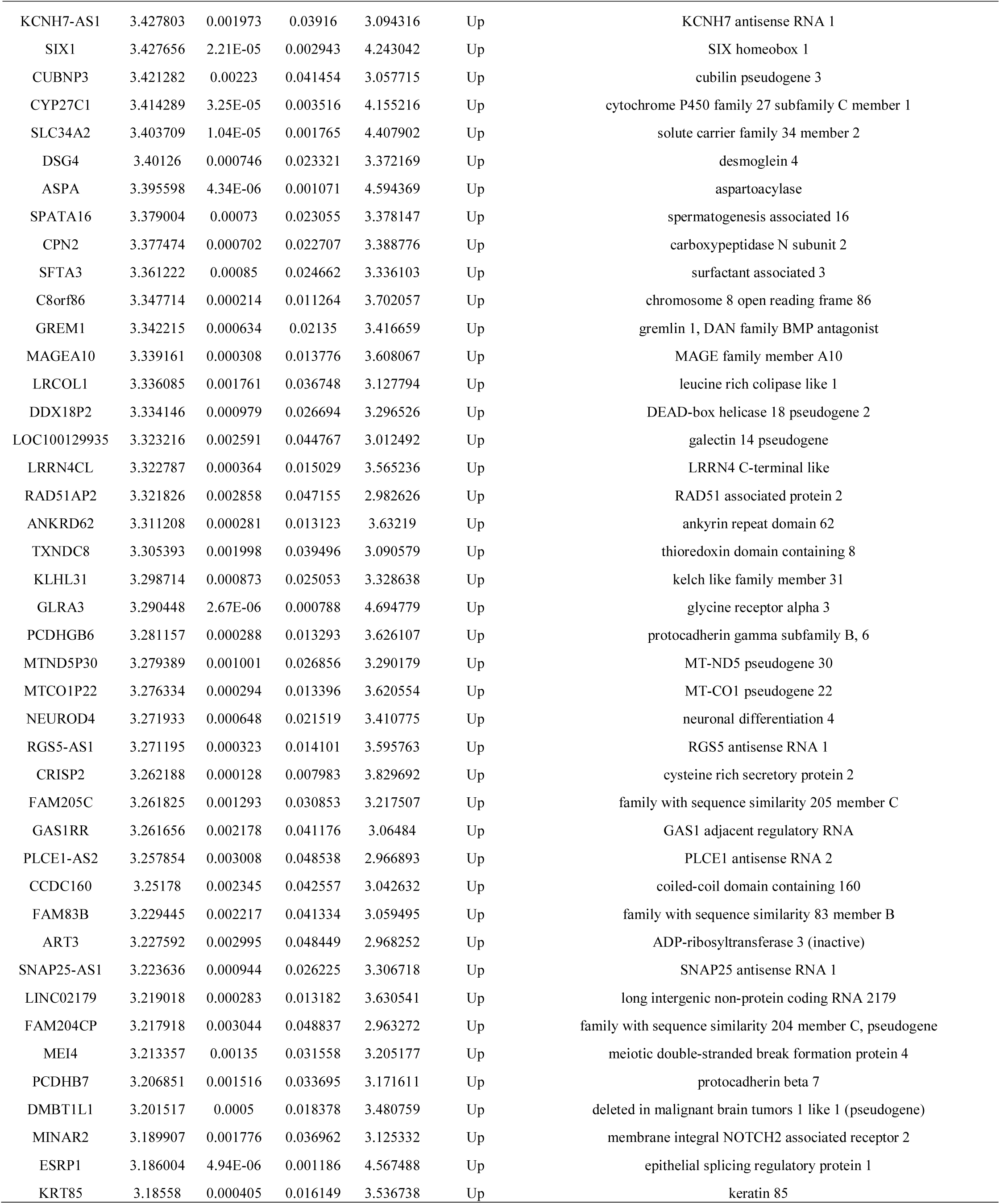
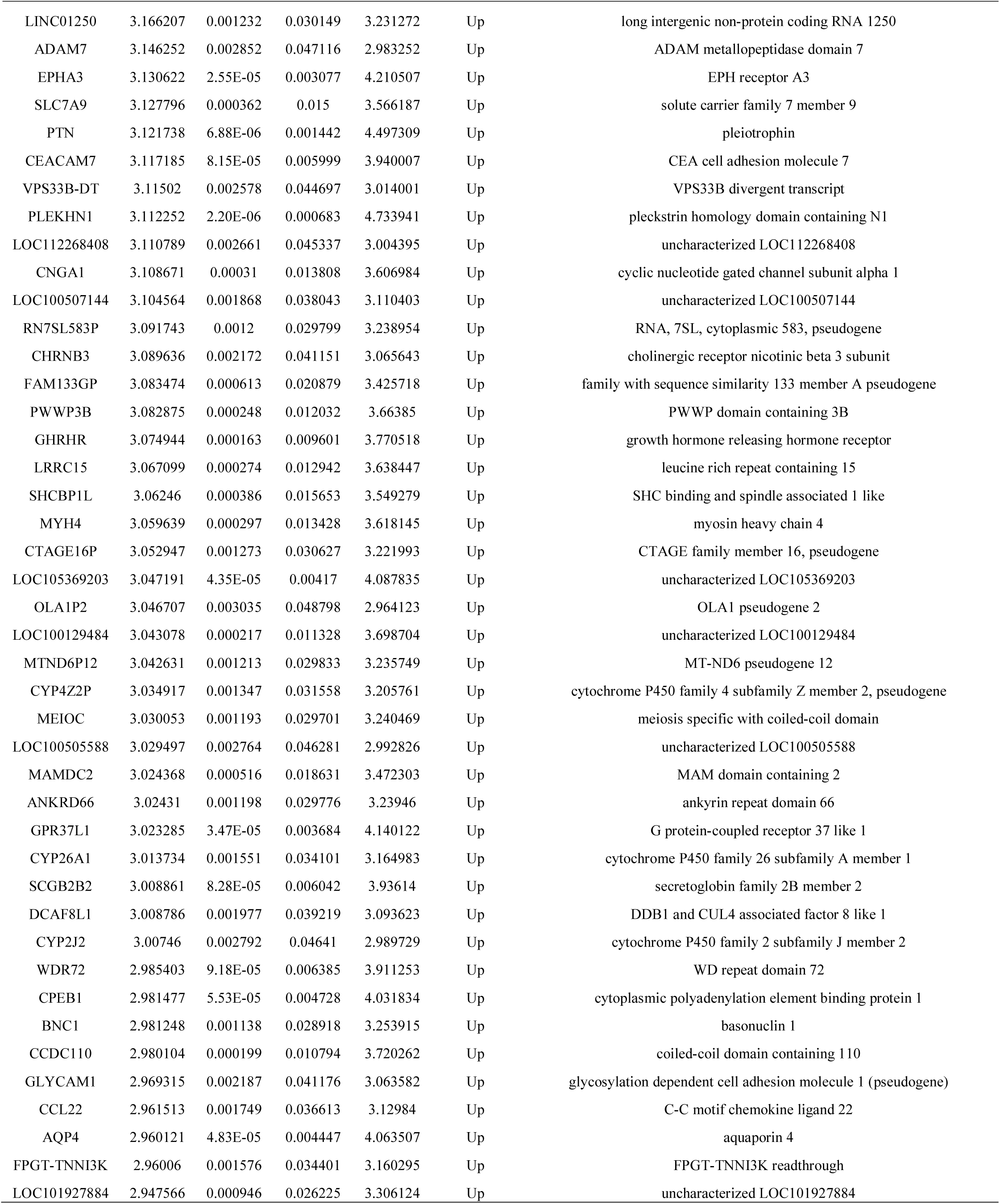
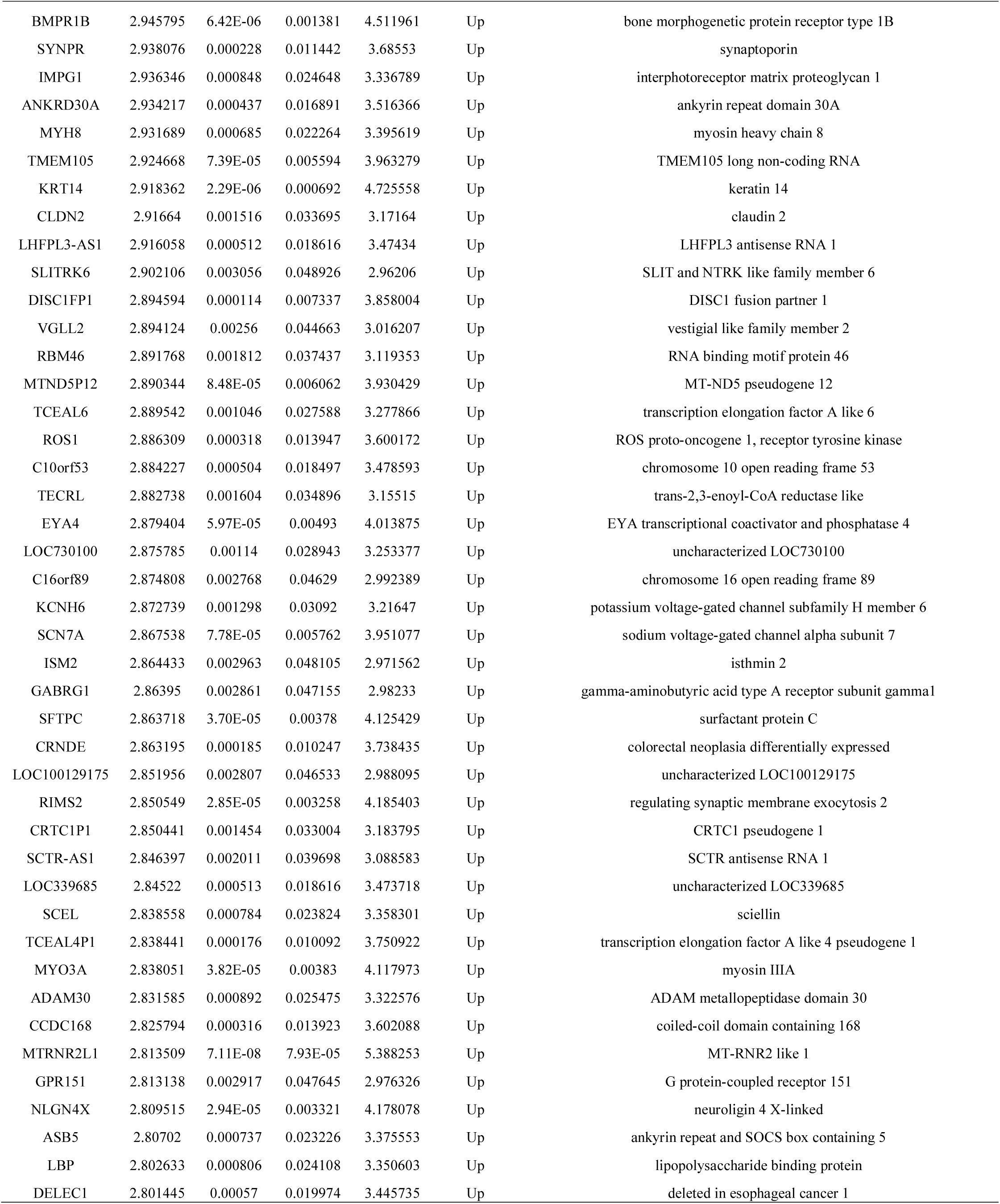
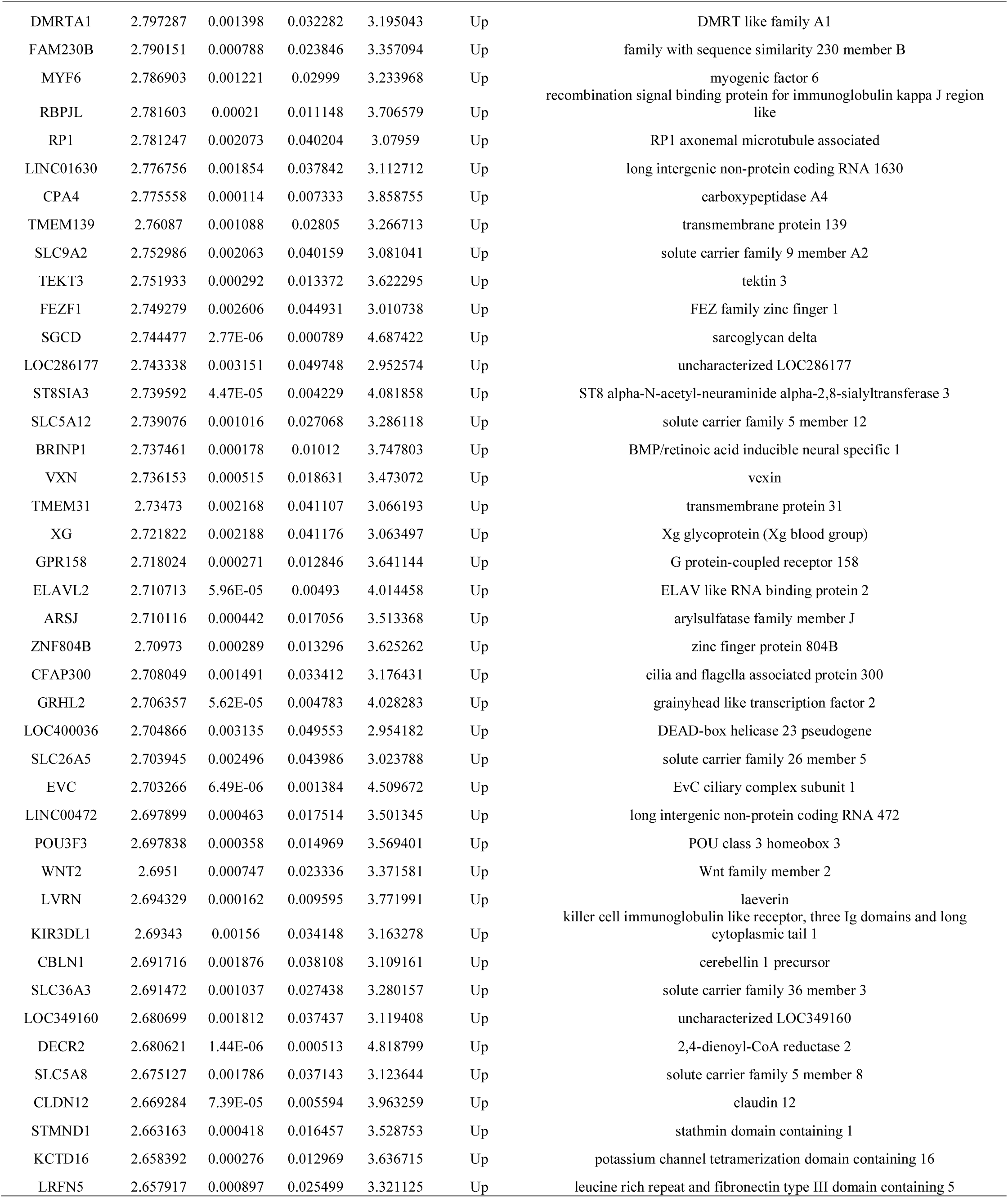
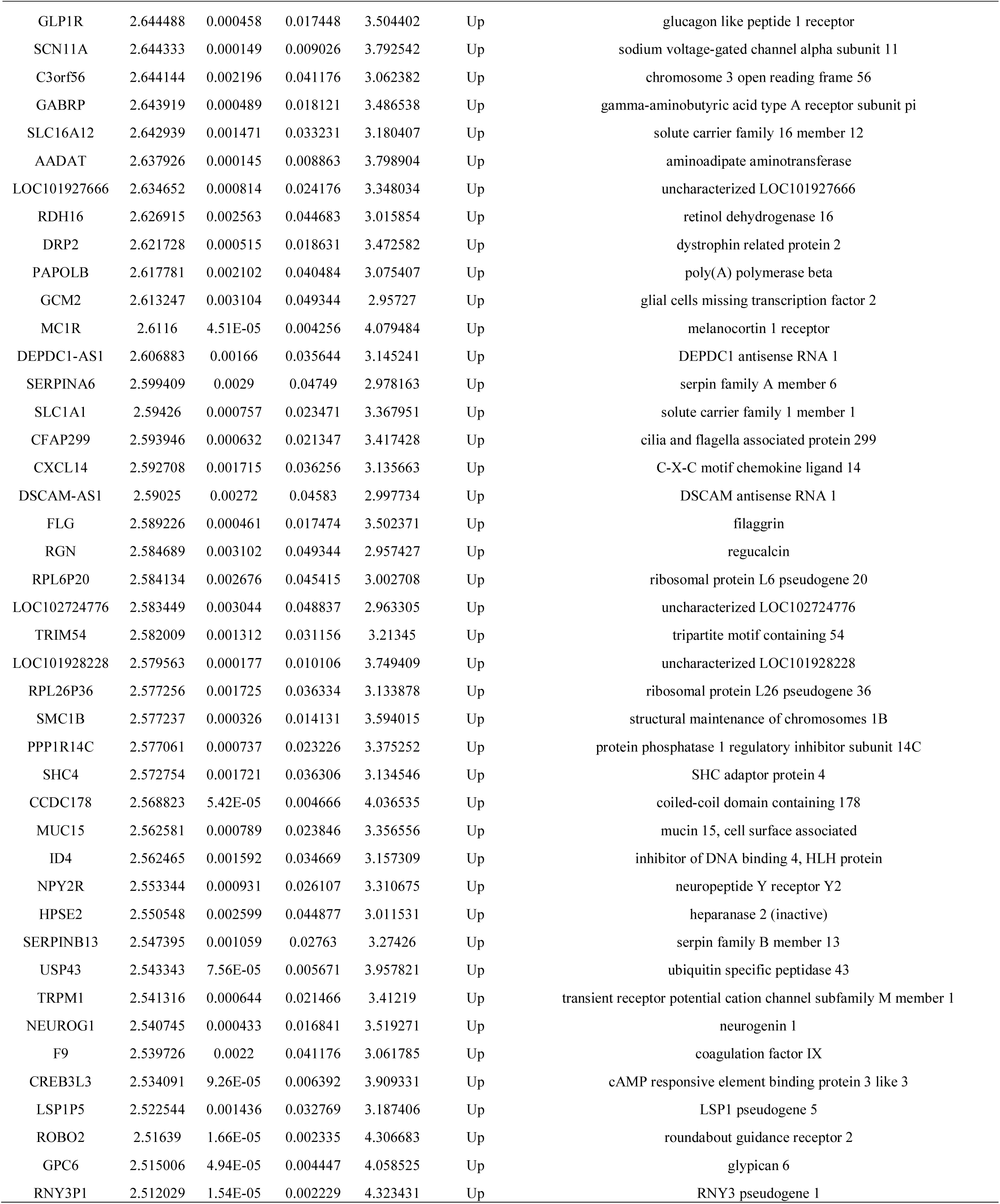
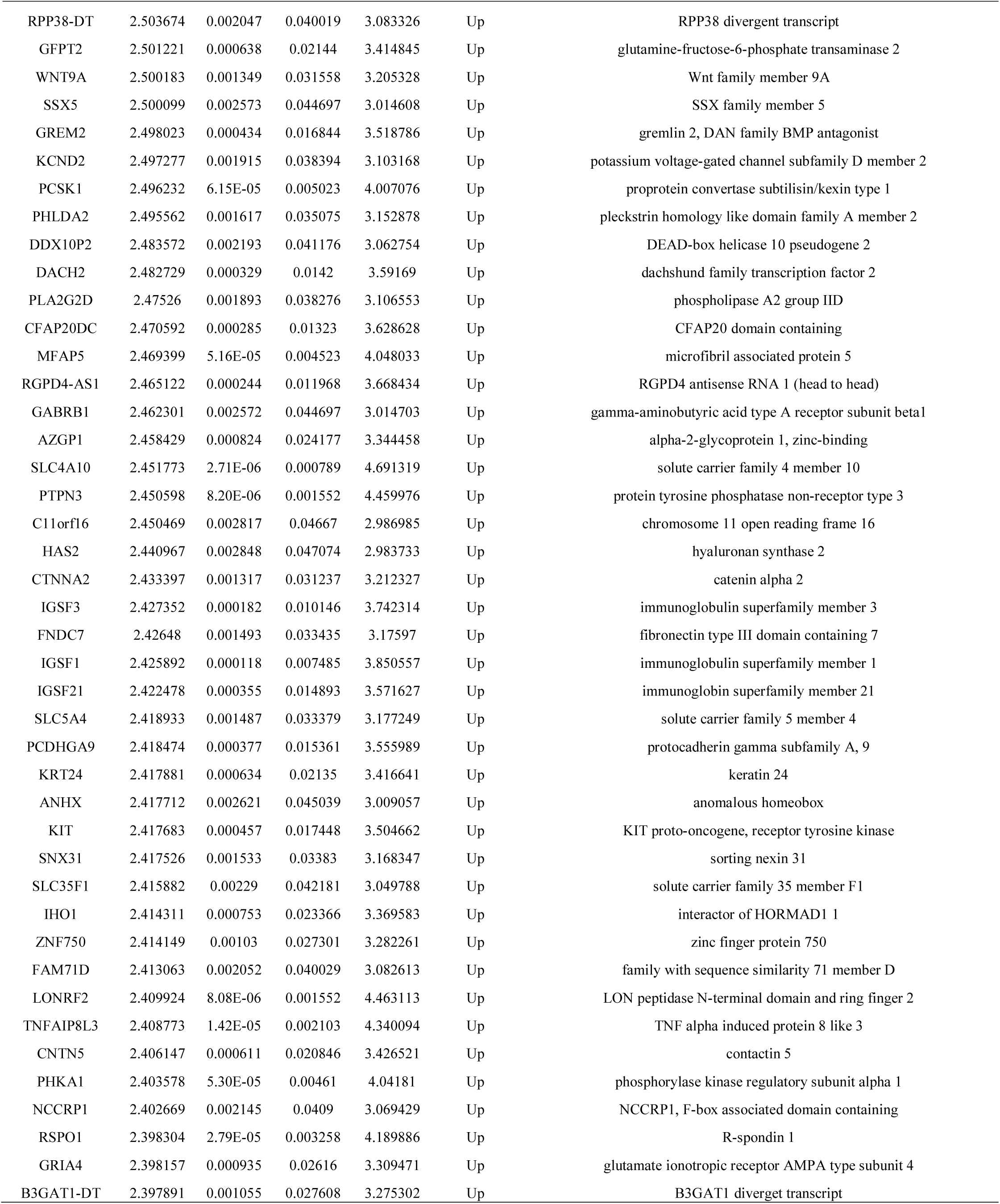
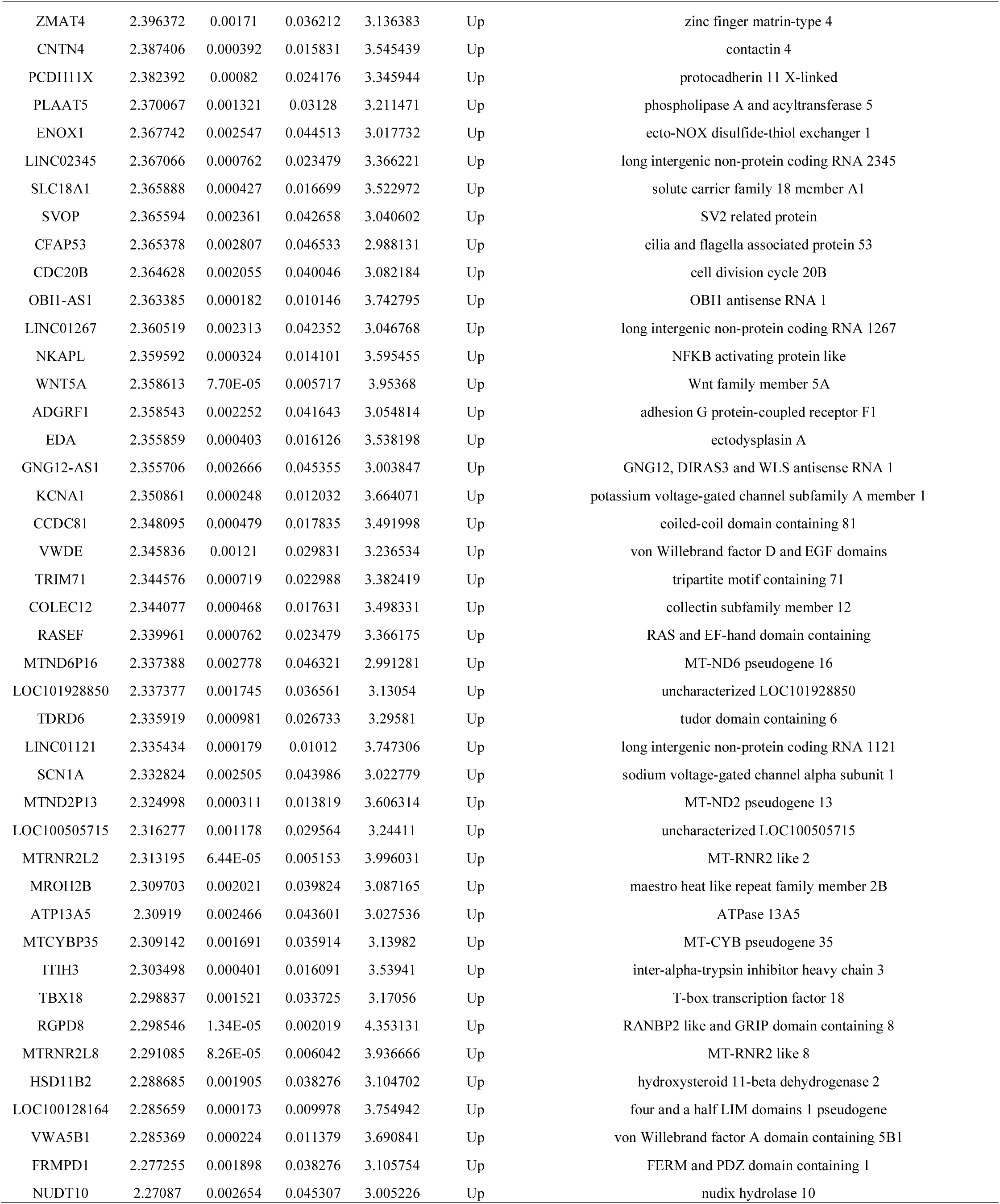
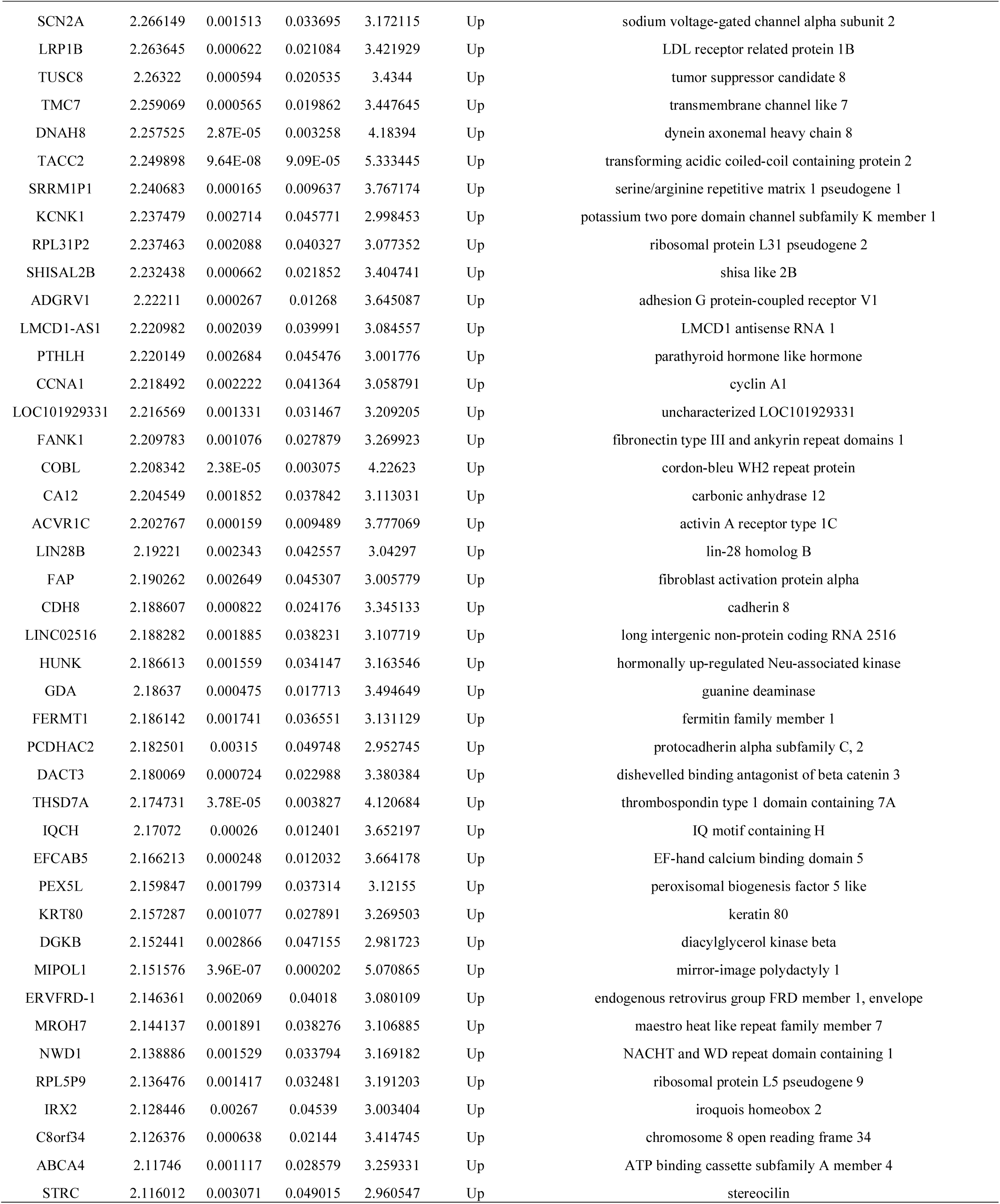
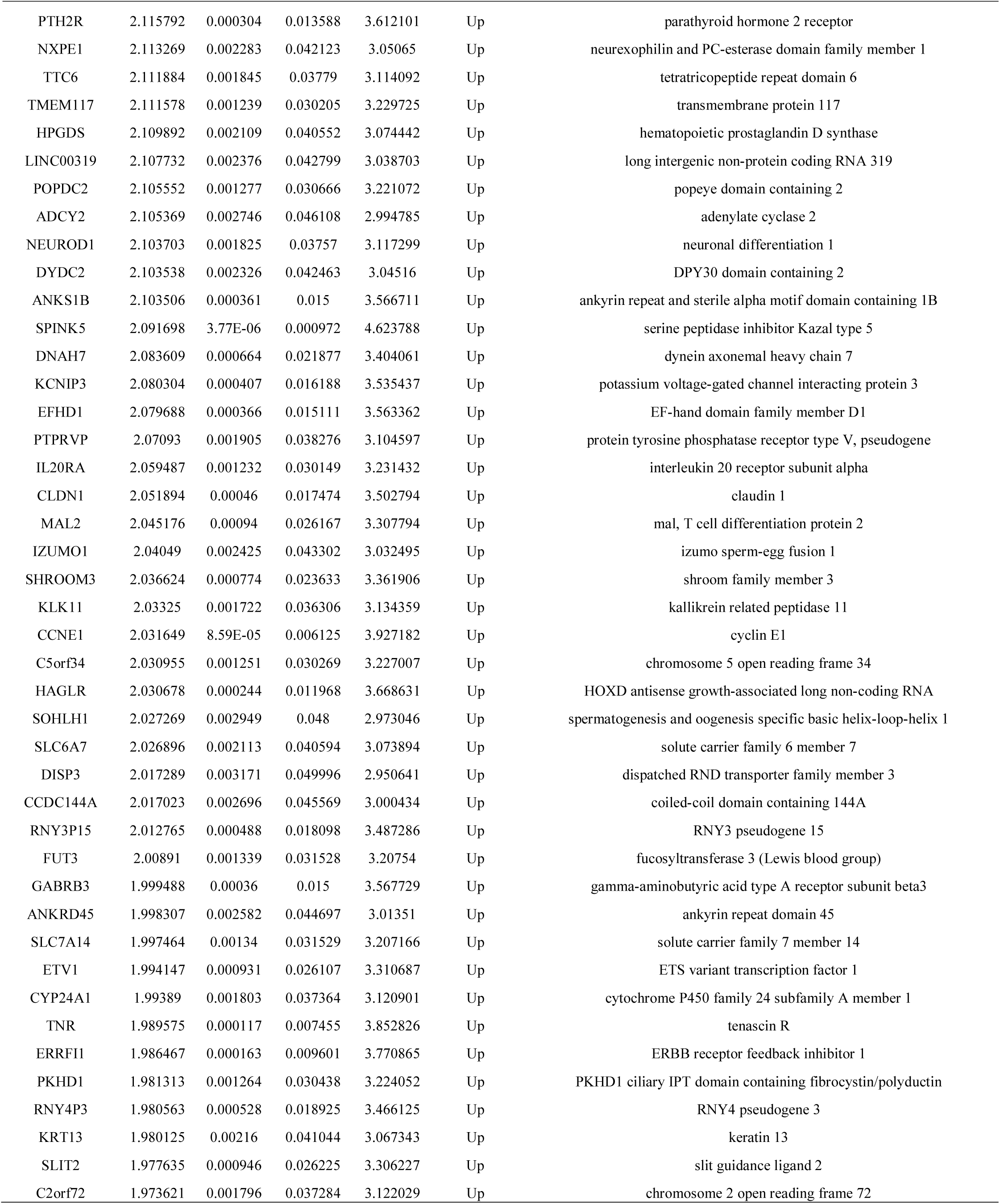
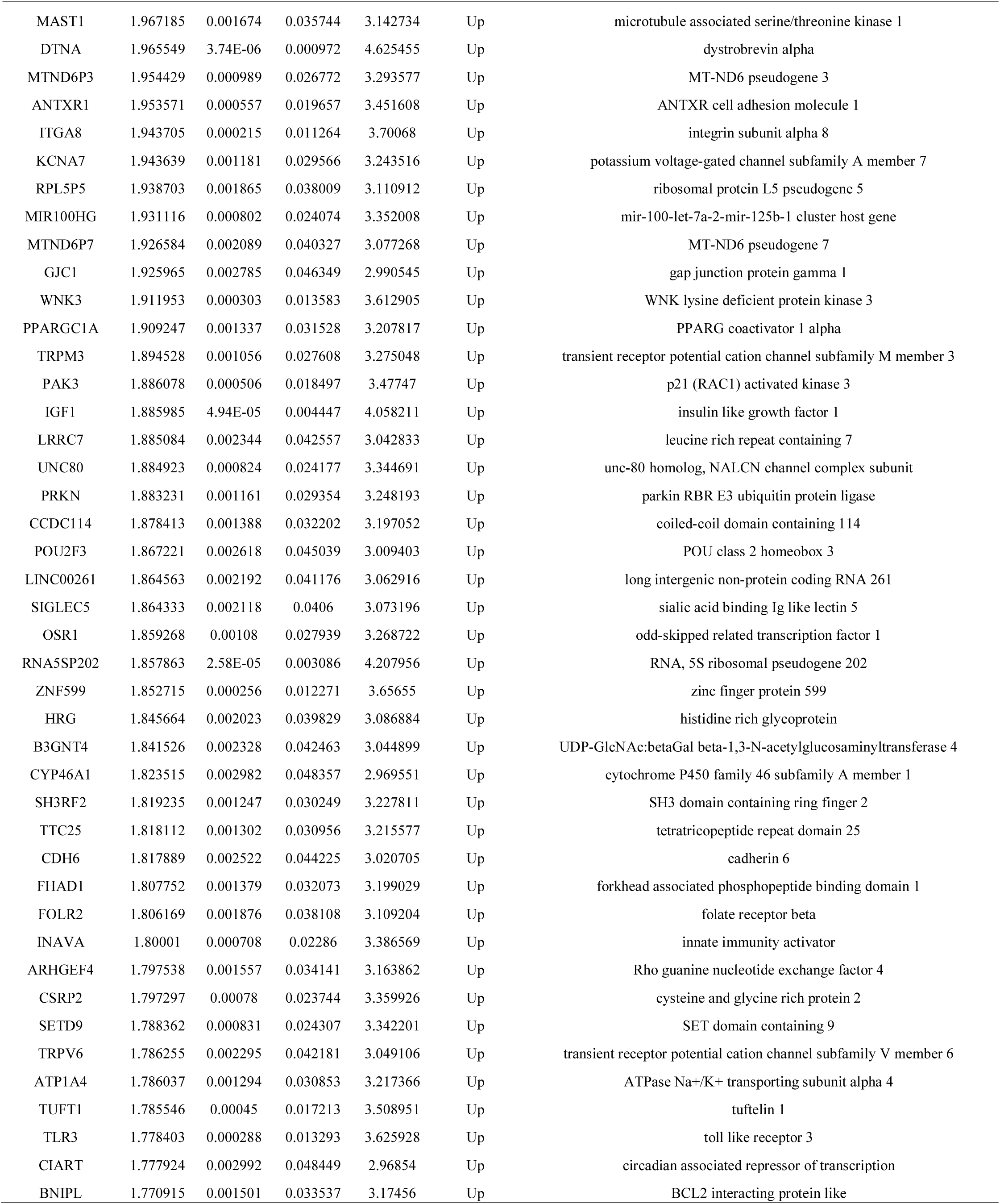
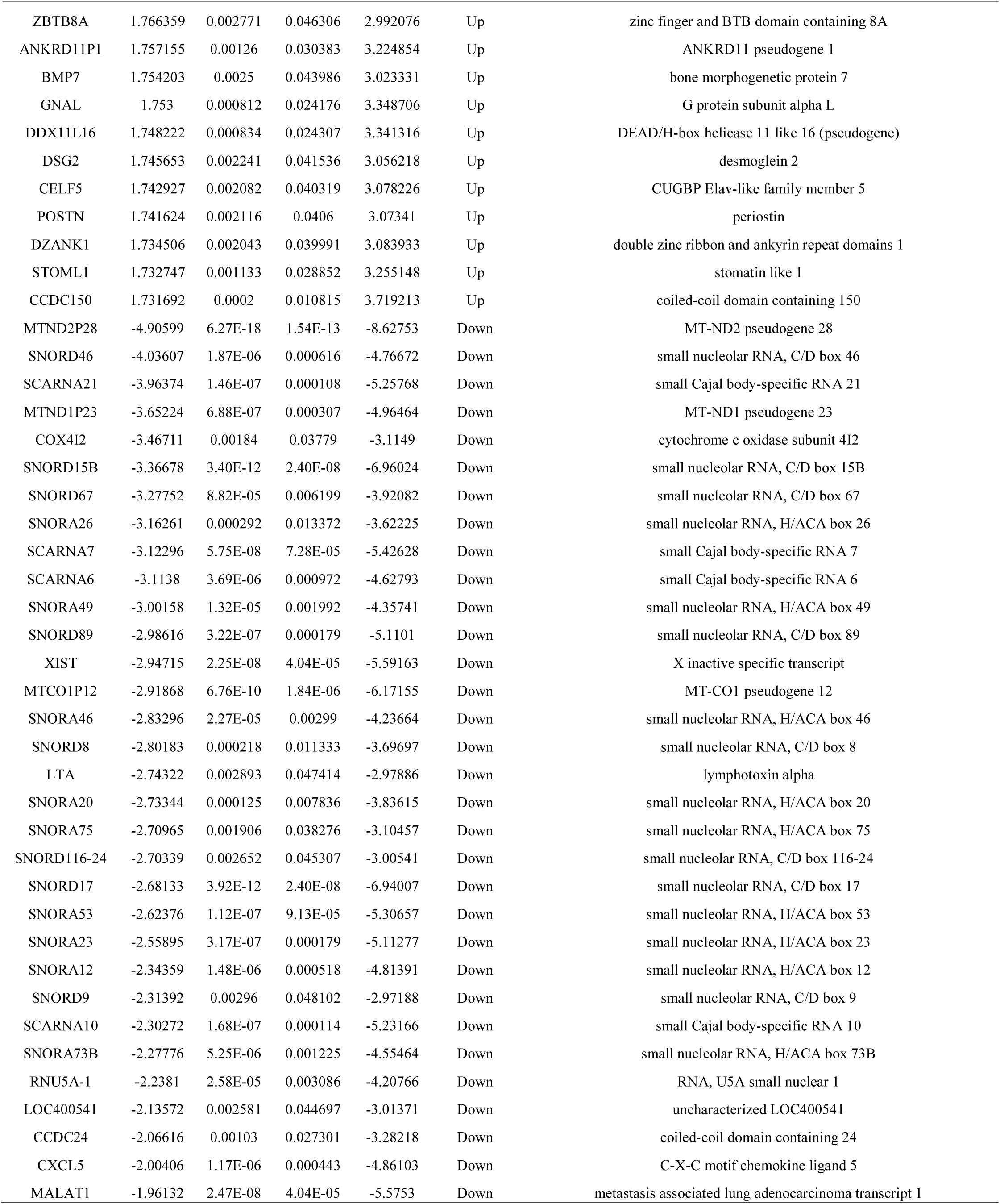
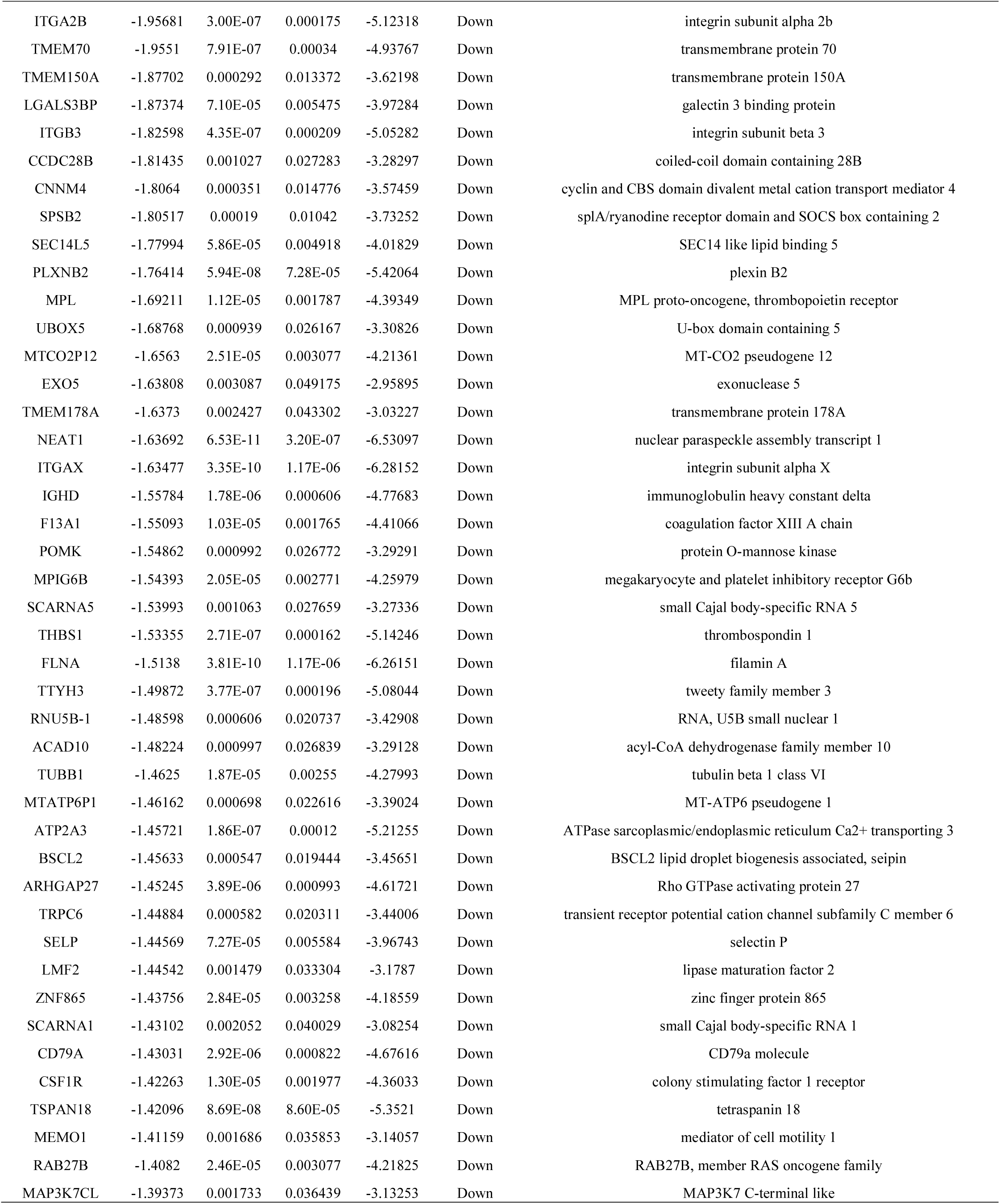
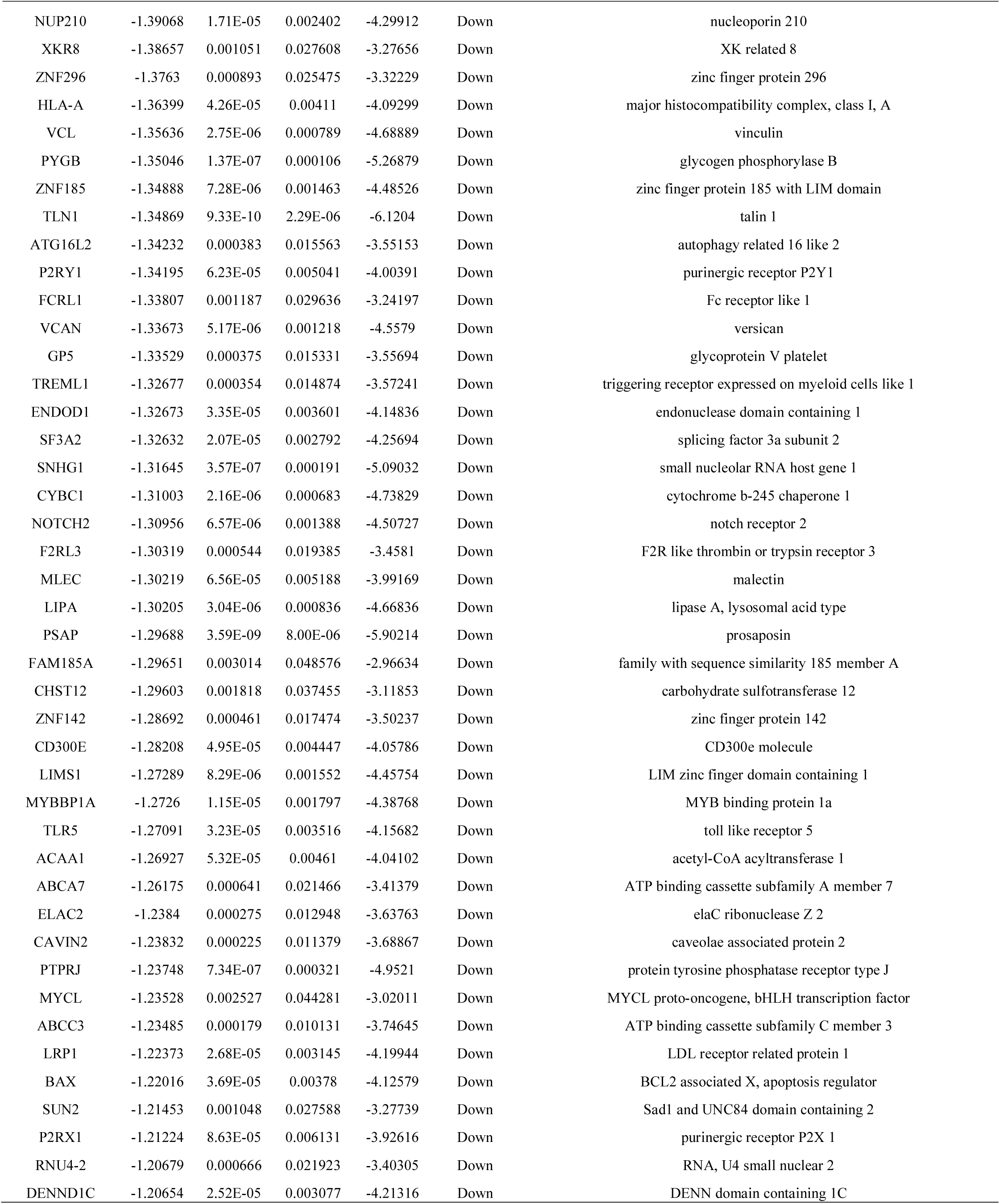
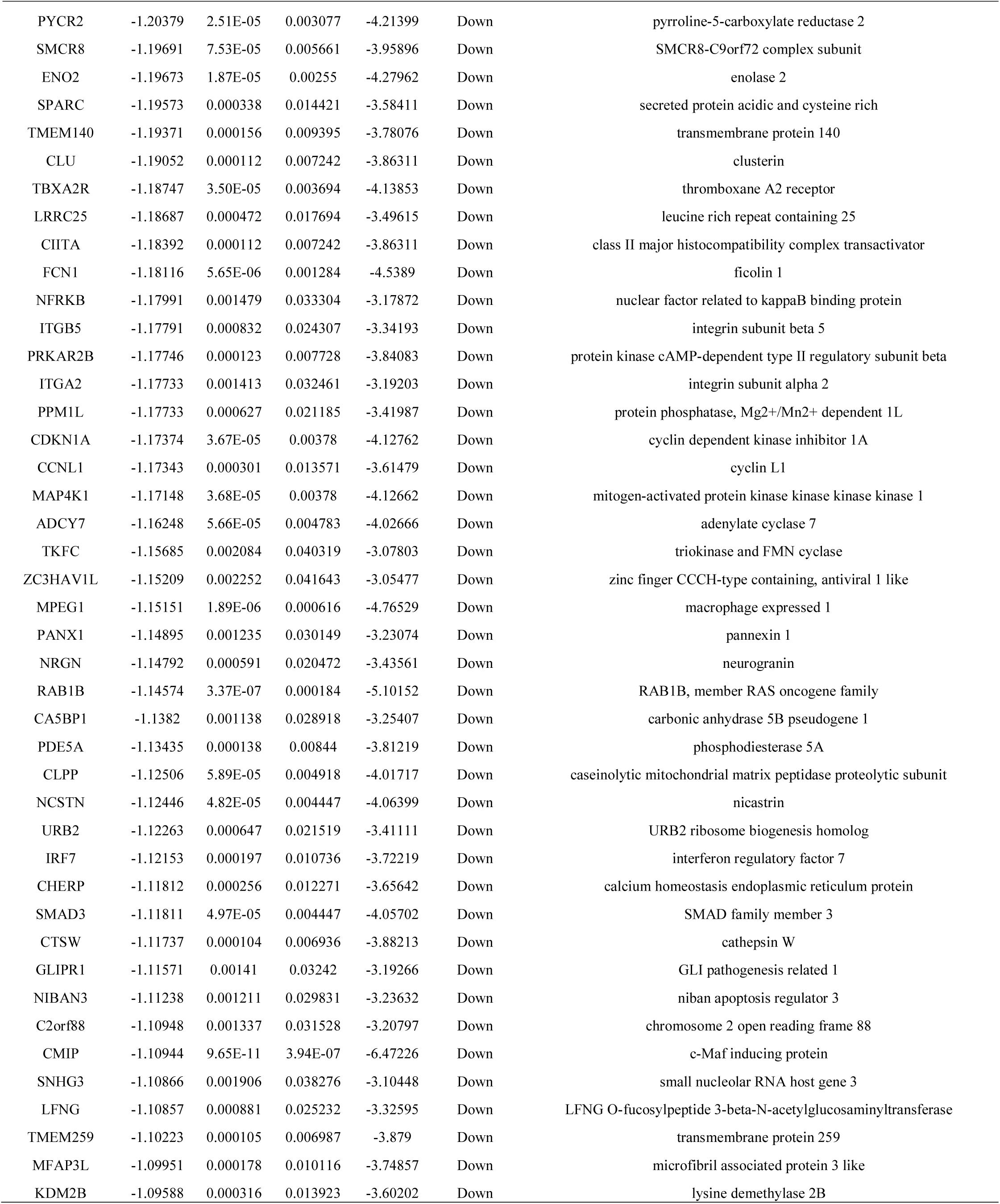
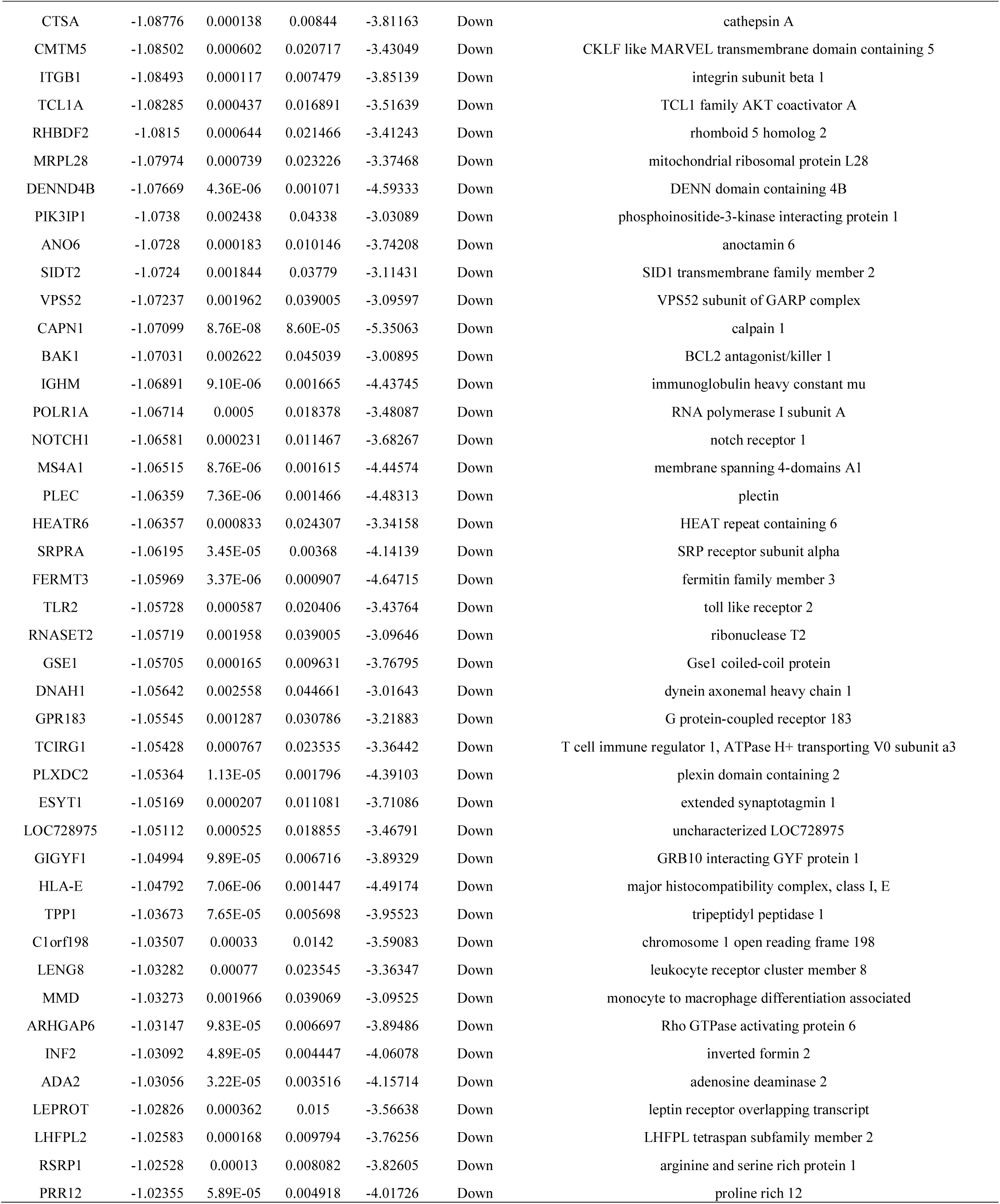
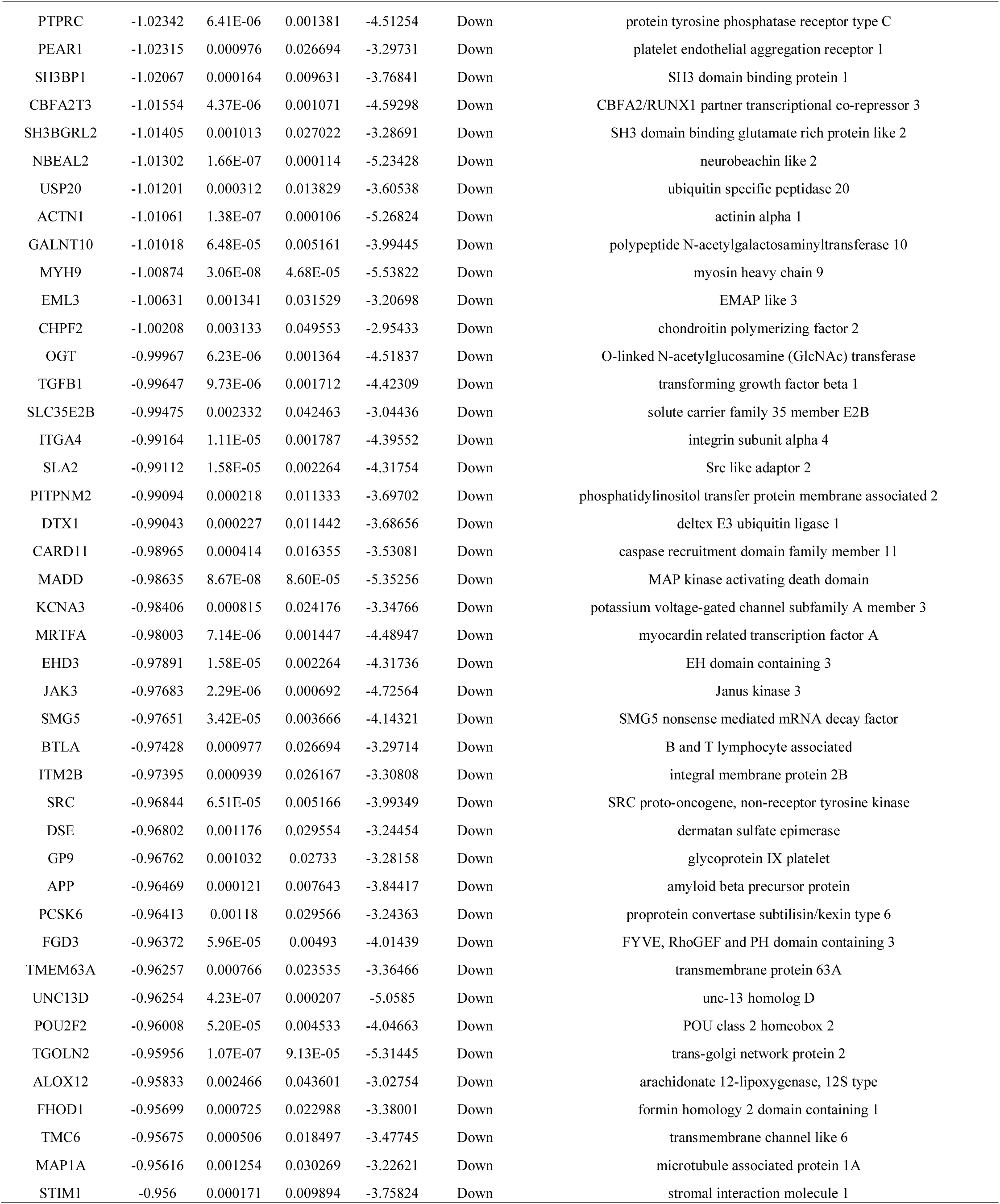
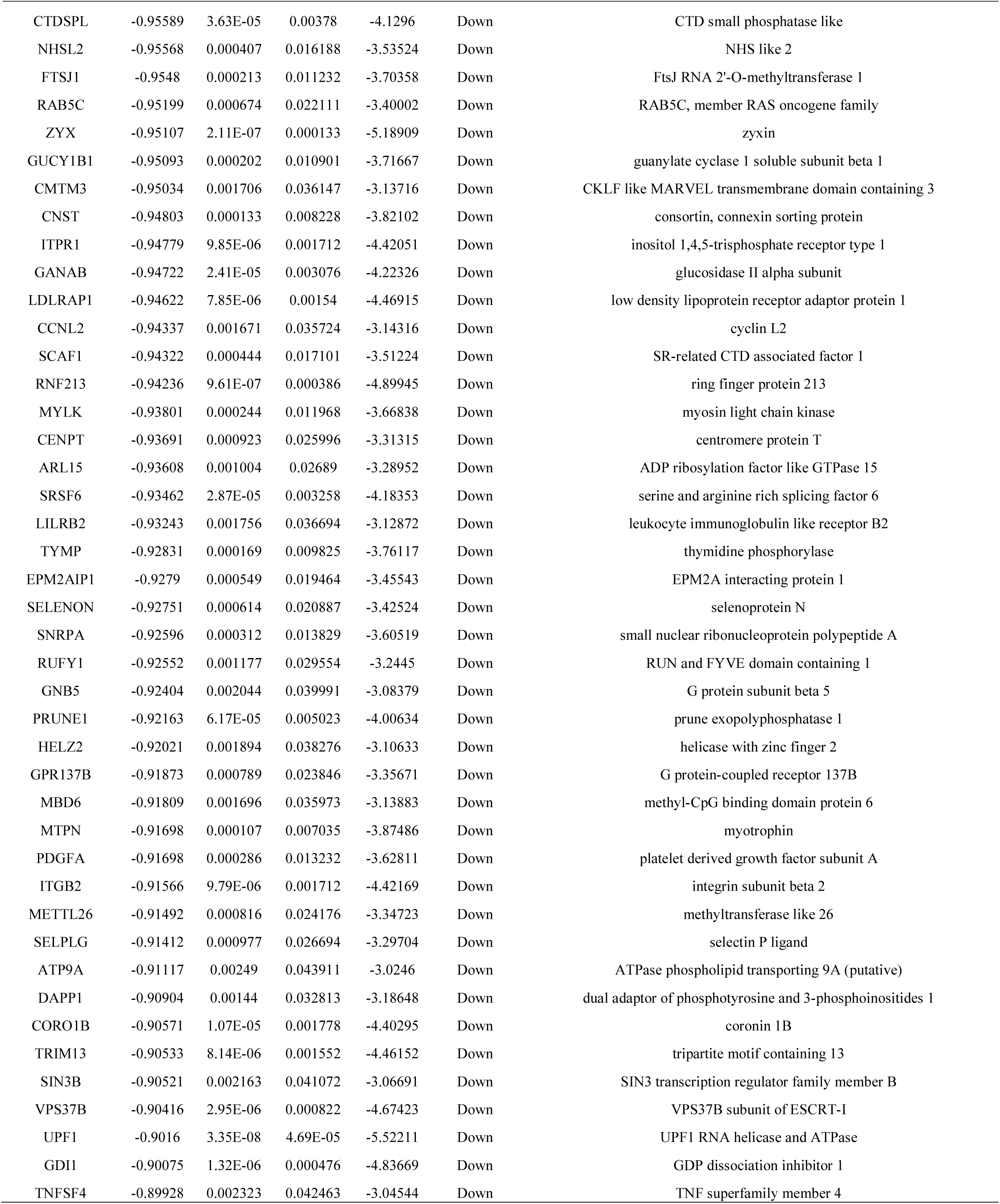
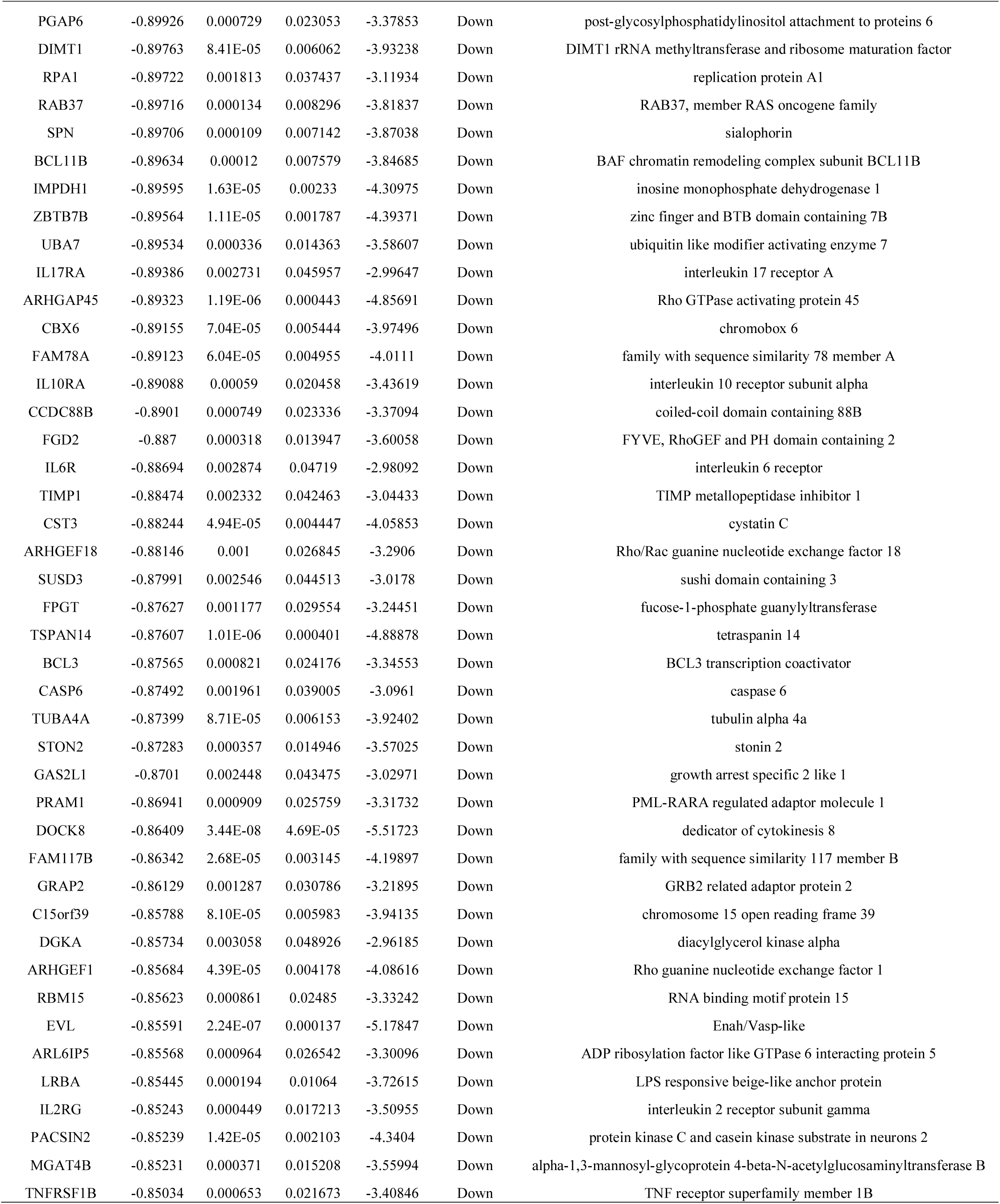
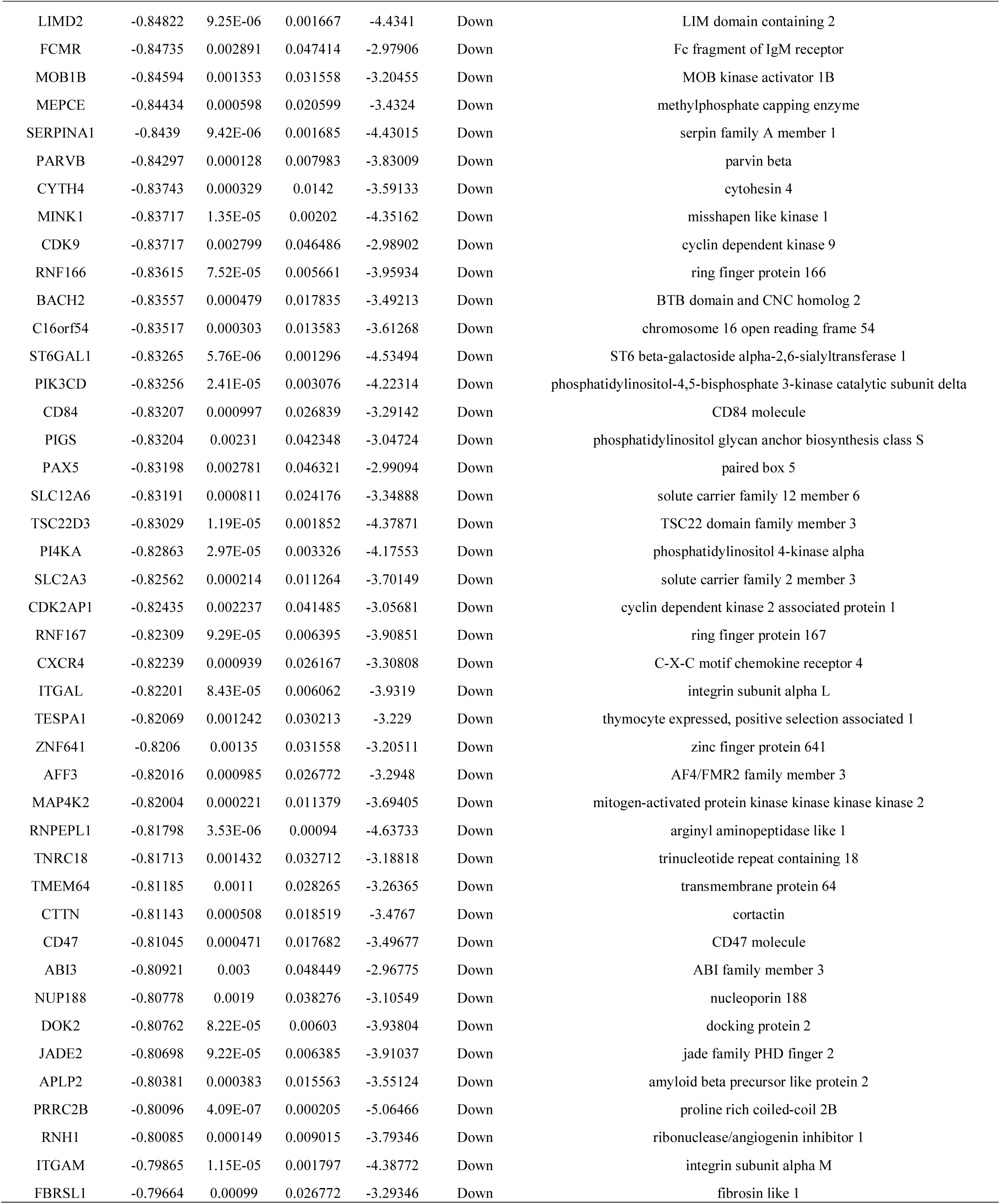
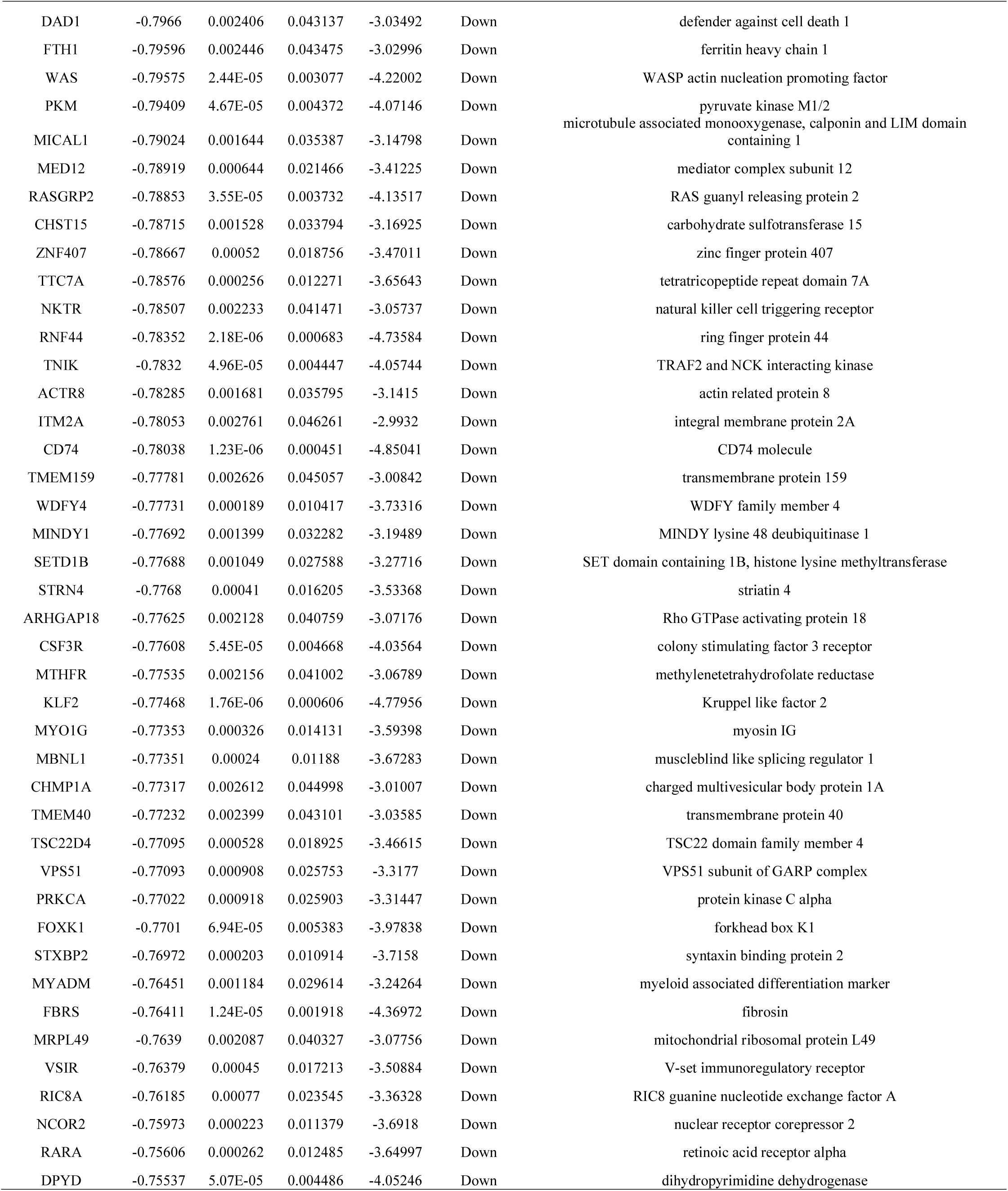
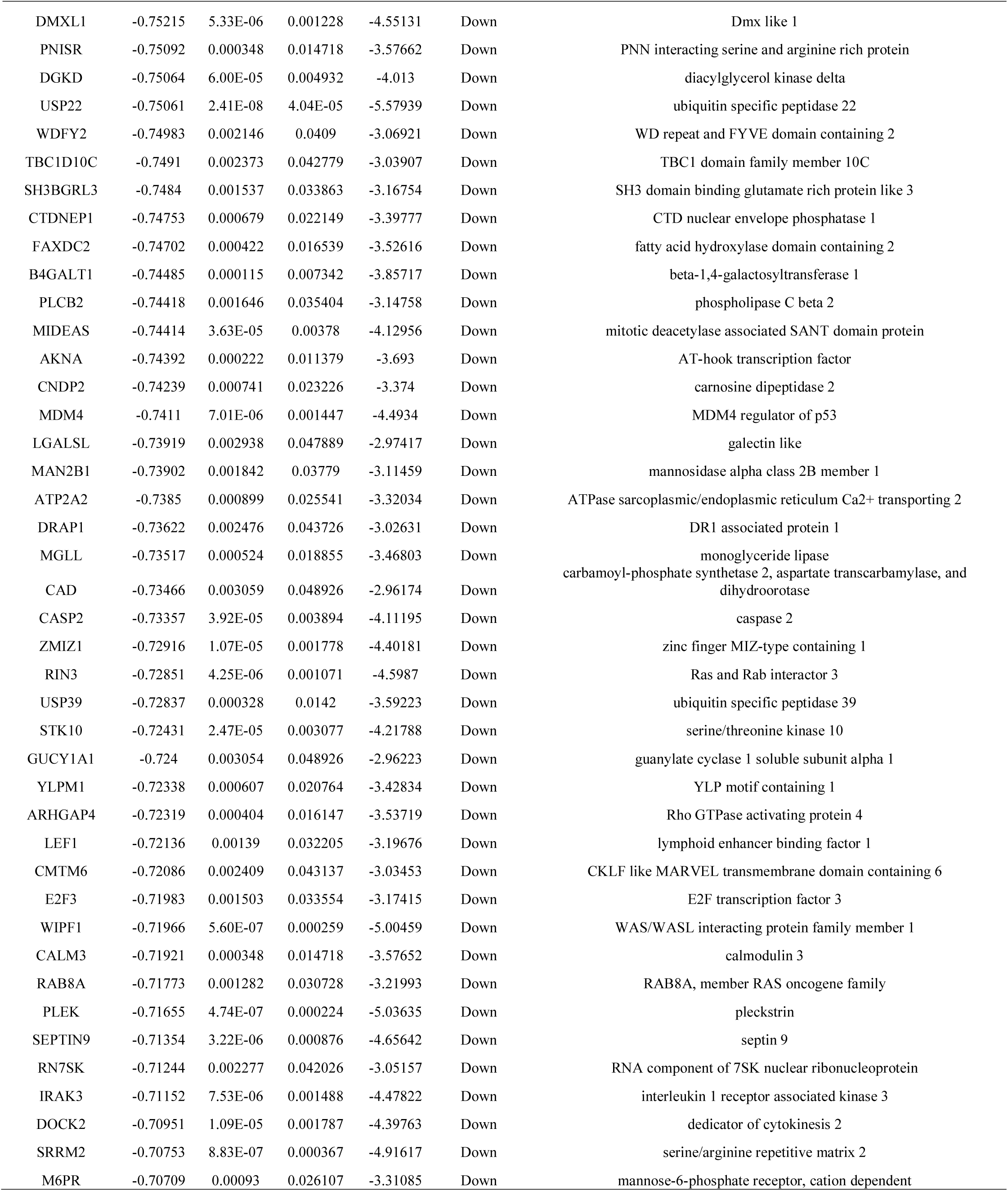
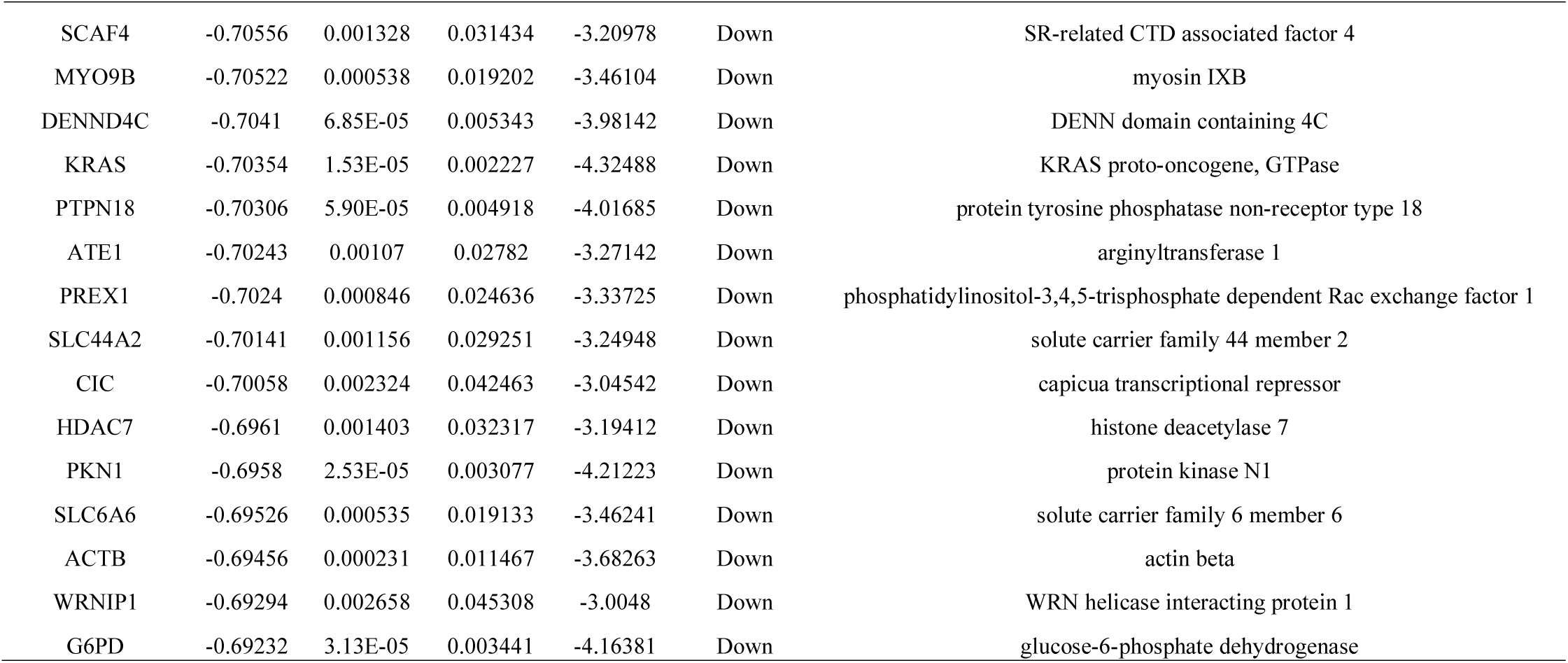
The statistical metrics for key differentially expressed genes (DEGs)

### Gene Ontology (GO) and REACTOME pathway enrichment analysis of DEGs

All 953 DEGs were analyzed by g:Profiler software and the results of GO enrichment analysis indicated that 1) for BP, up regulated genes were particularly enriched in multicellular organismal process and developmental process, and down regulated genes in cell activation and immune system process; 2) for CC, up regulated genes were enriched in cell periphery and intrinsic component of membrane, and down regulated genes in endomembrane system and cytoplasm; 3) for GO MF, up regulated genes were significantly enriched in the transporter activity and channel activity, and down regulated geness in enzyme binding and protein binding (Table 2). REACTOME pathway enrichment analysis results were shown in Table 3 which demonstrated that up regulated genes were particularly enriched in formation of the cornified envelope and cell-cell junction organization, while down regulated genes were particularly enriched in hemostasis and neutrophil degranulation (P_<_0.05).

**Table 2.**
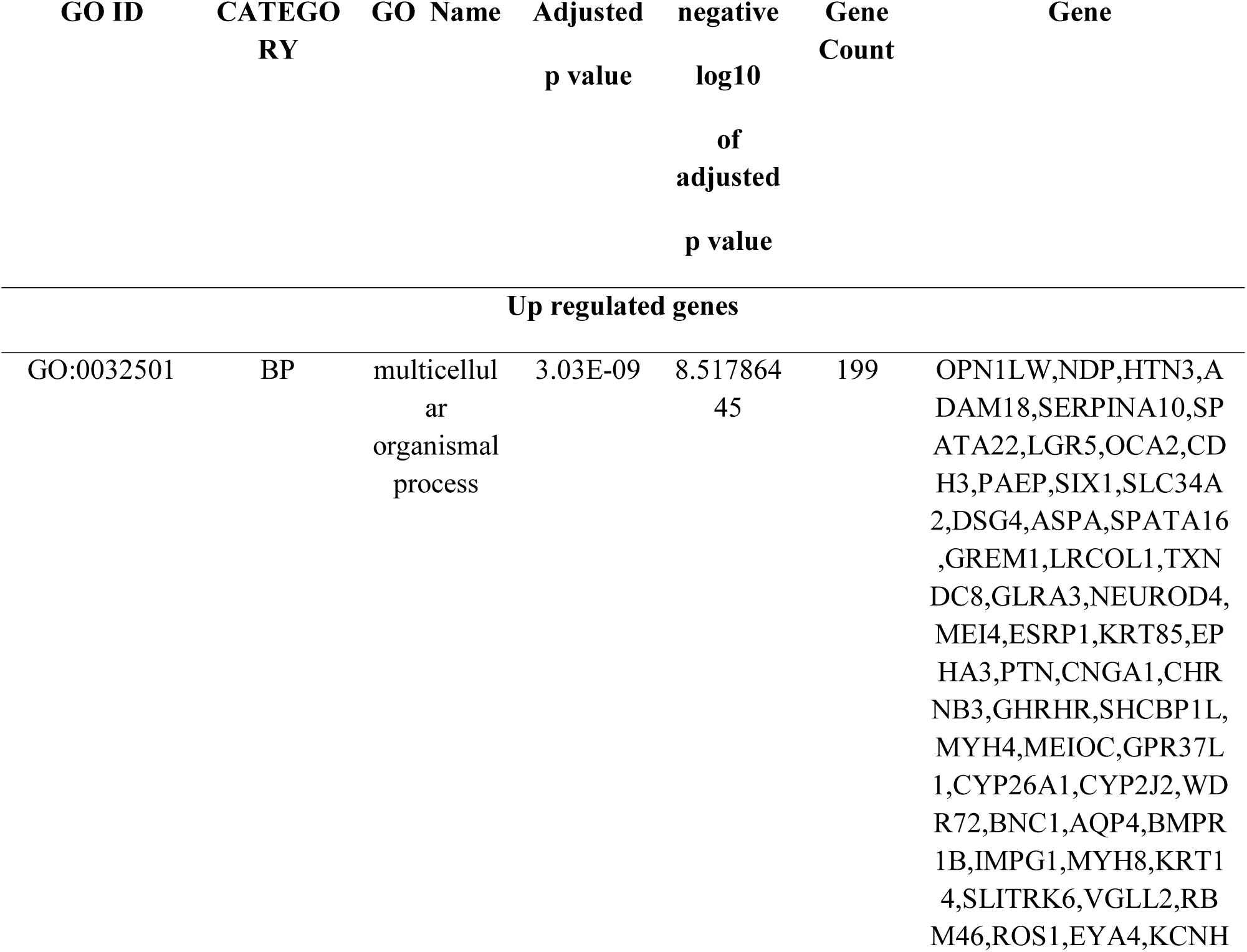
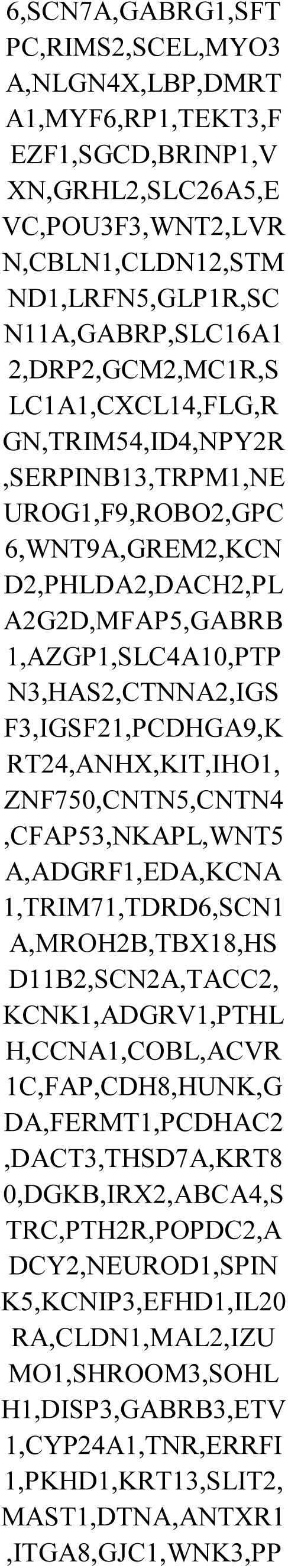
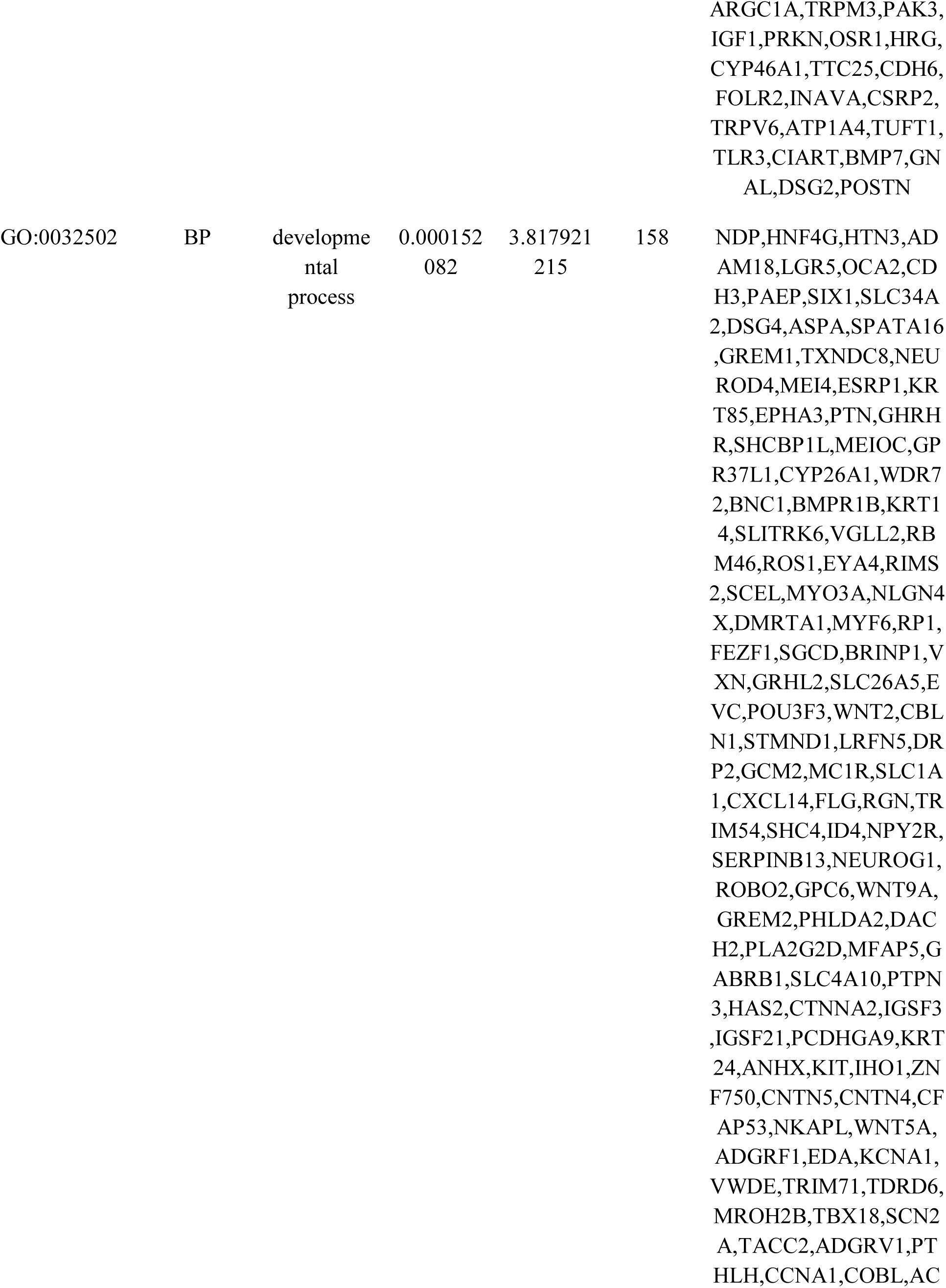
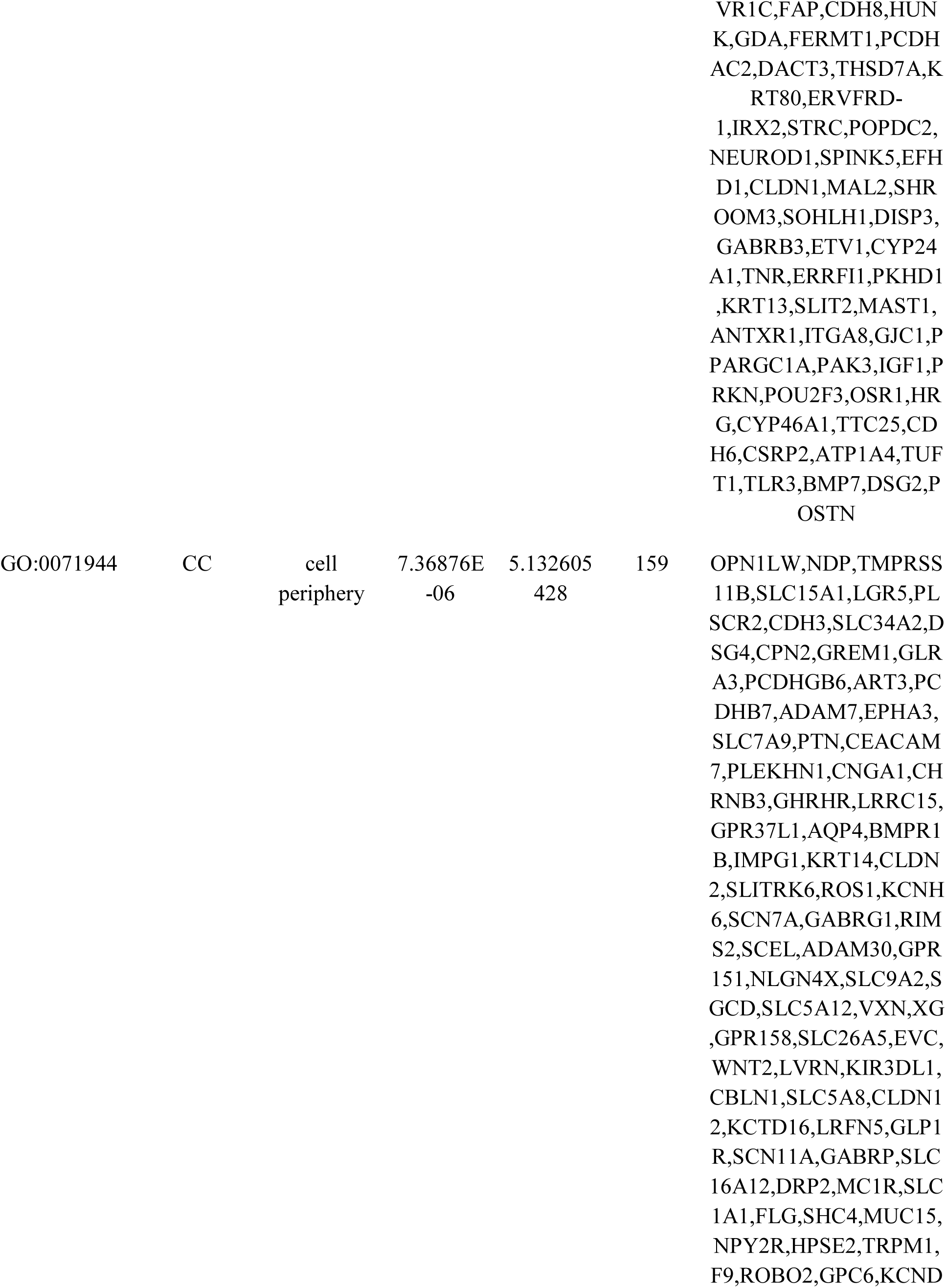
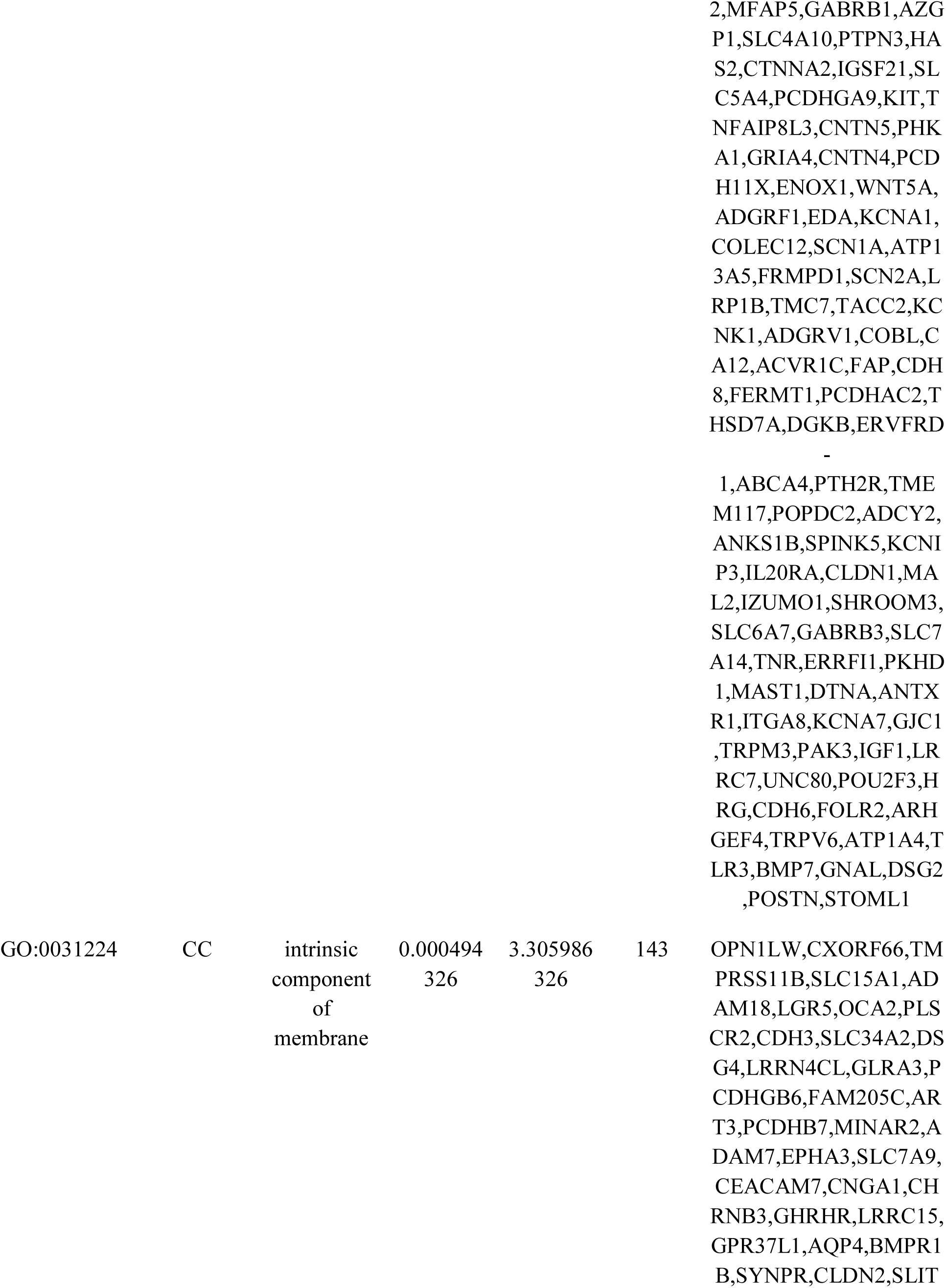
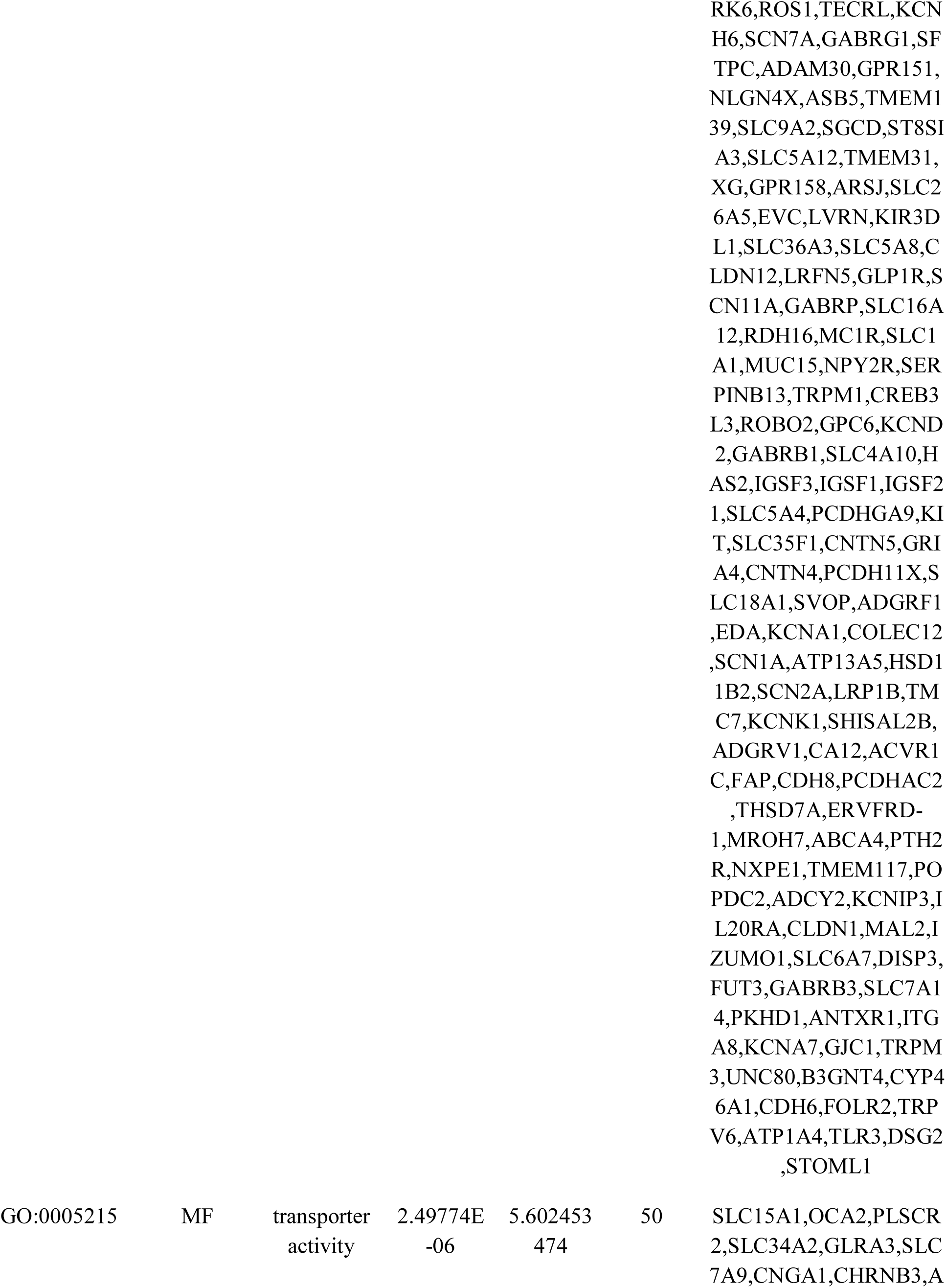
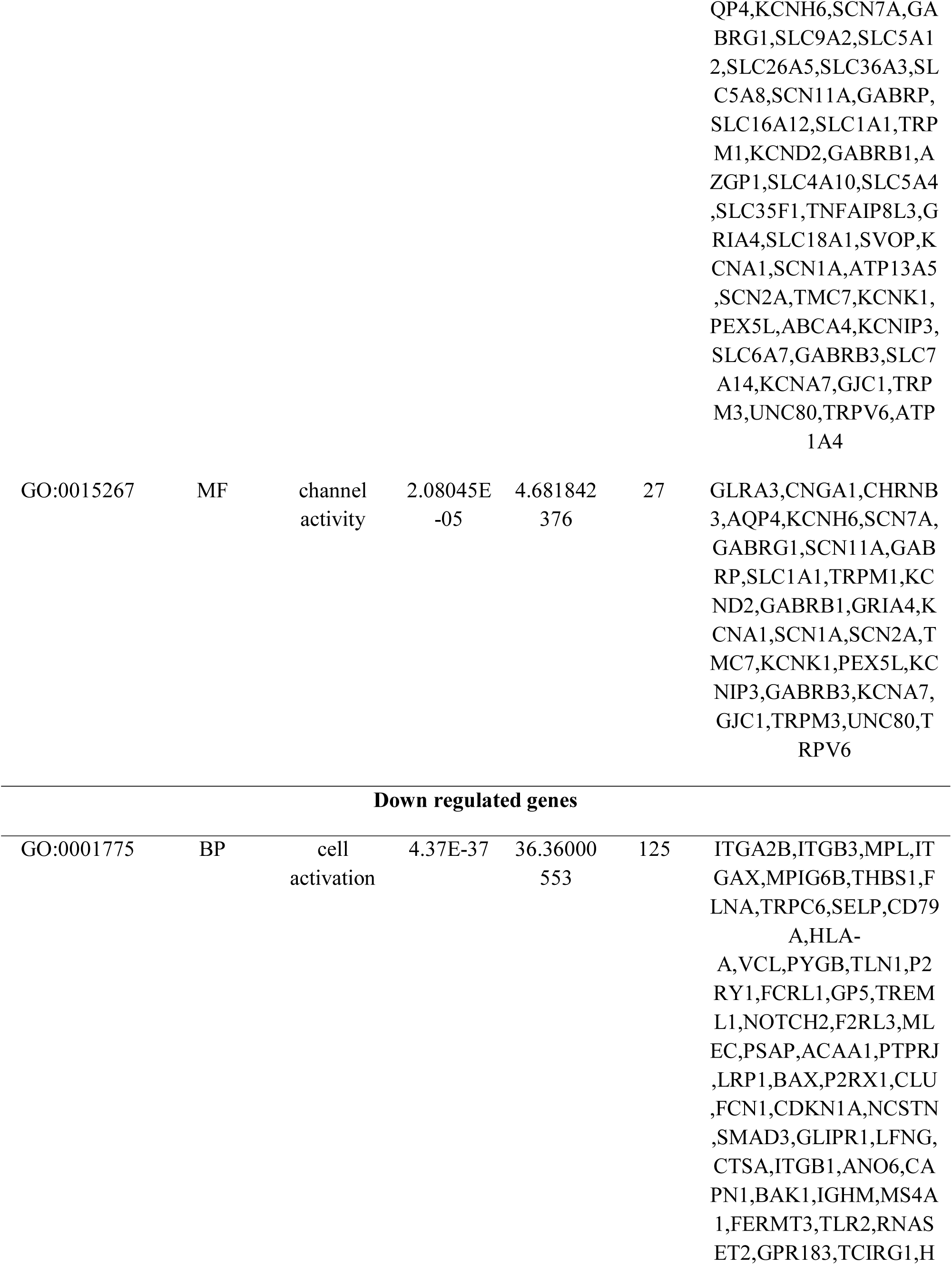
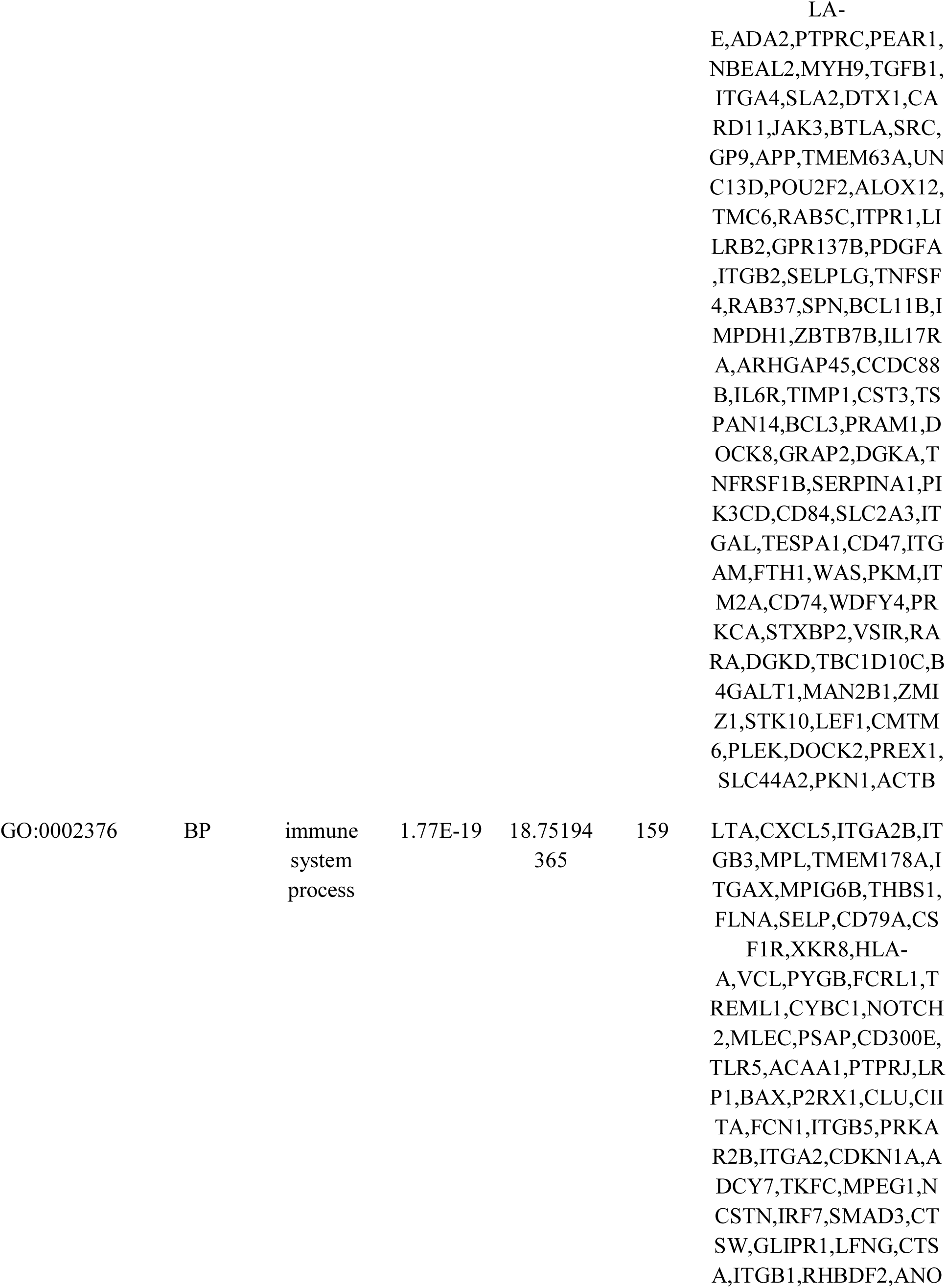
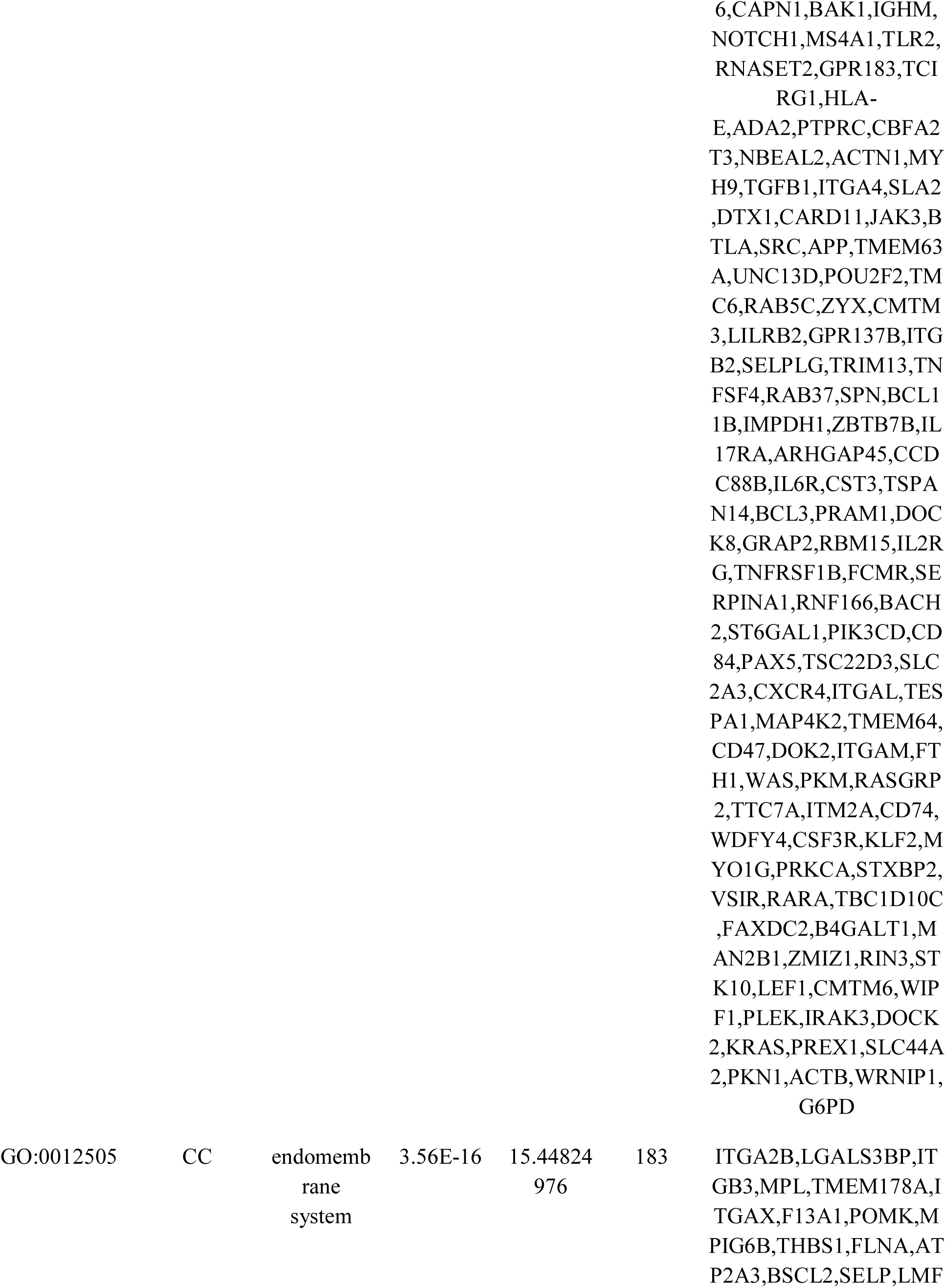
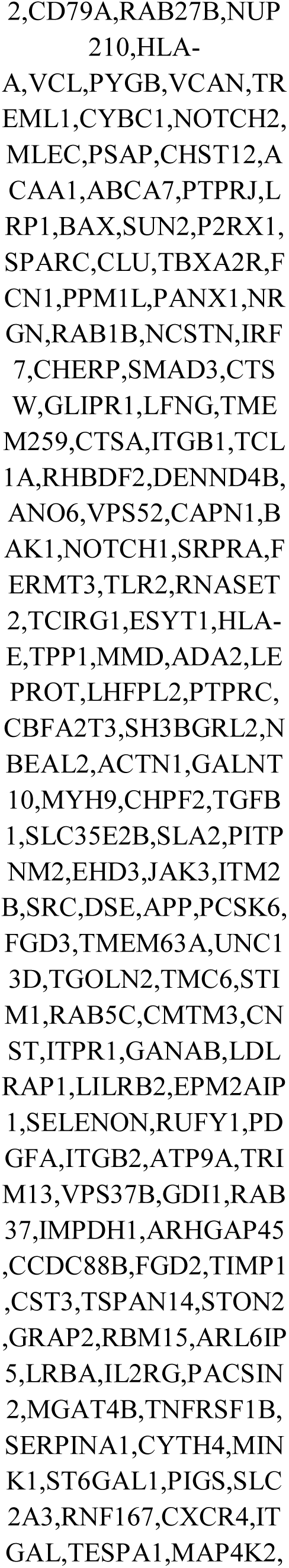
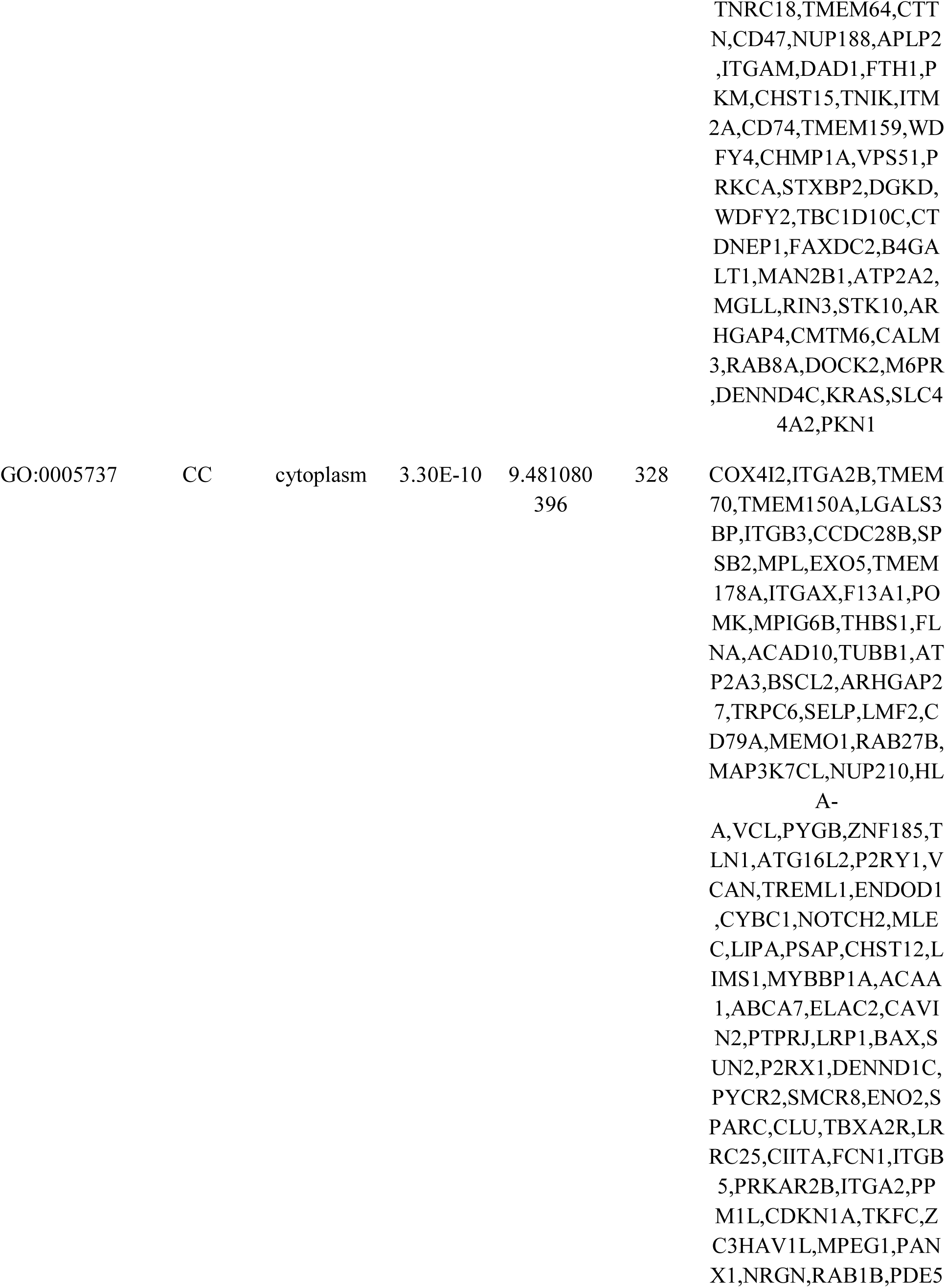
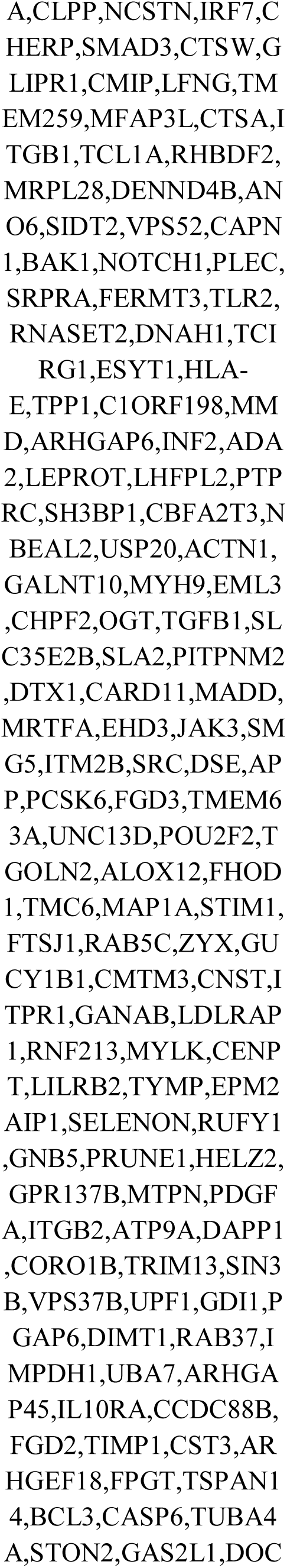
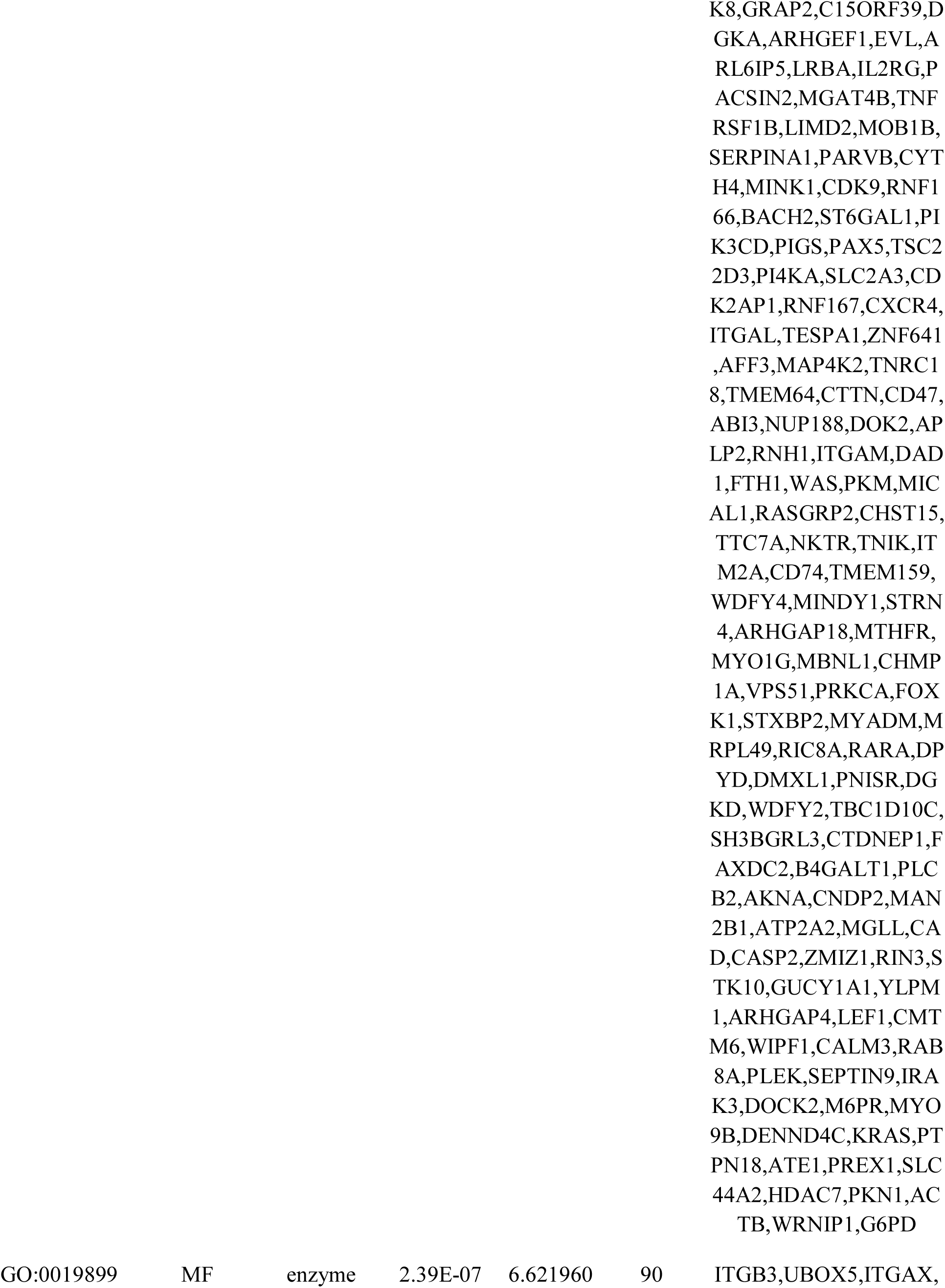
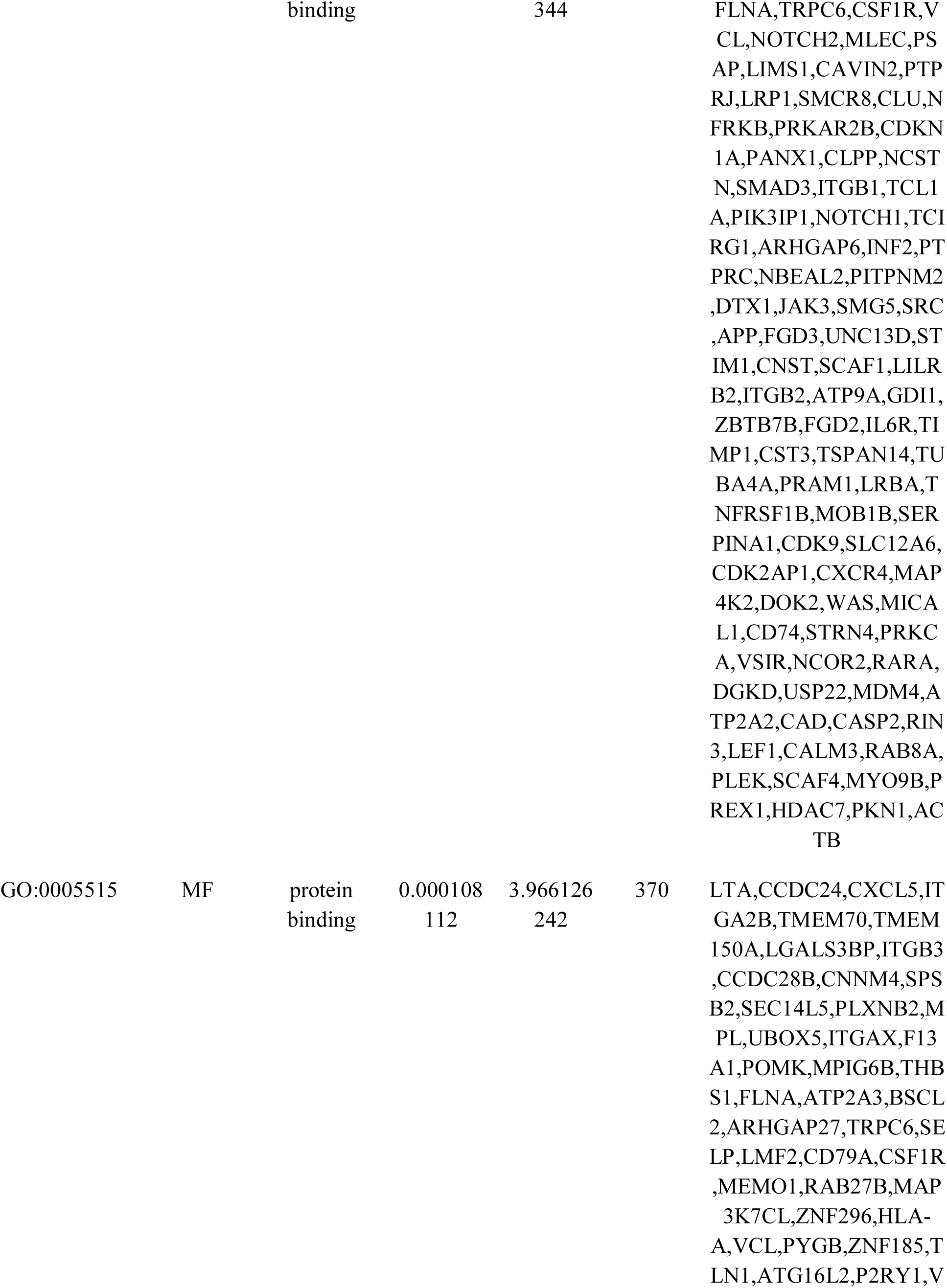
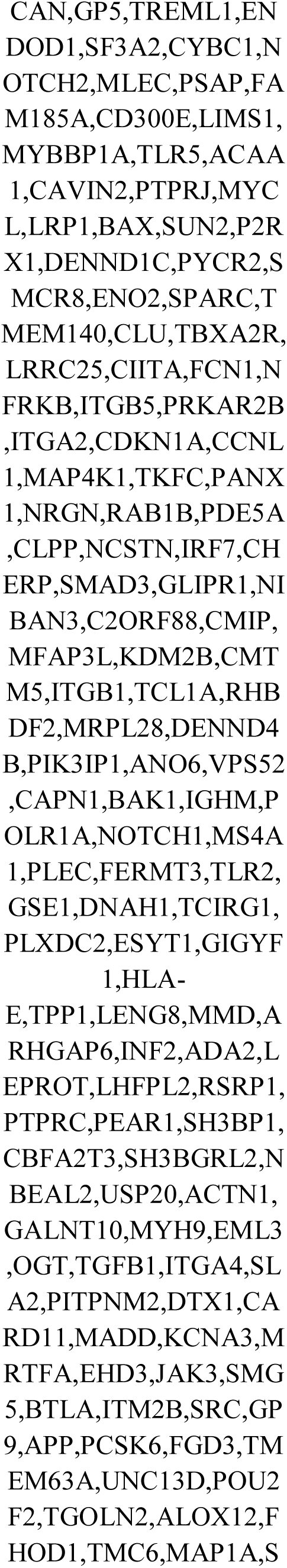
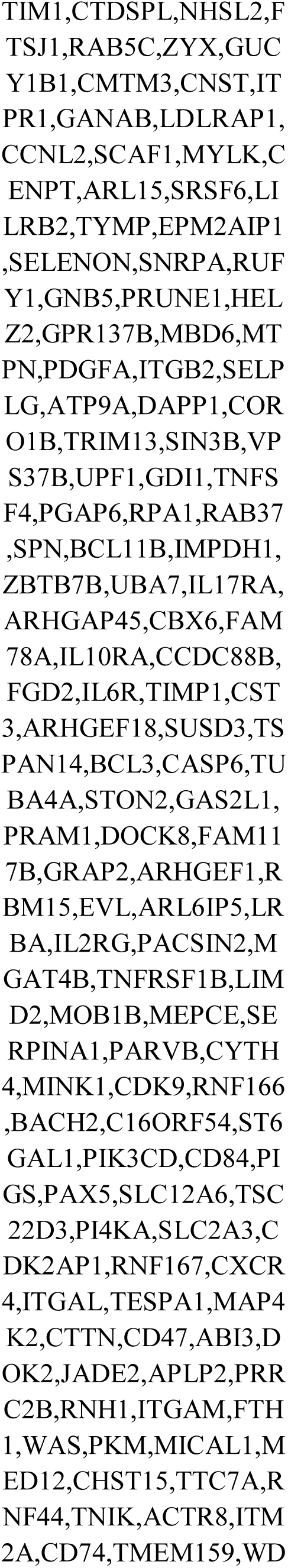
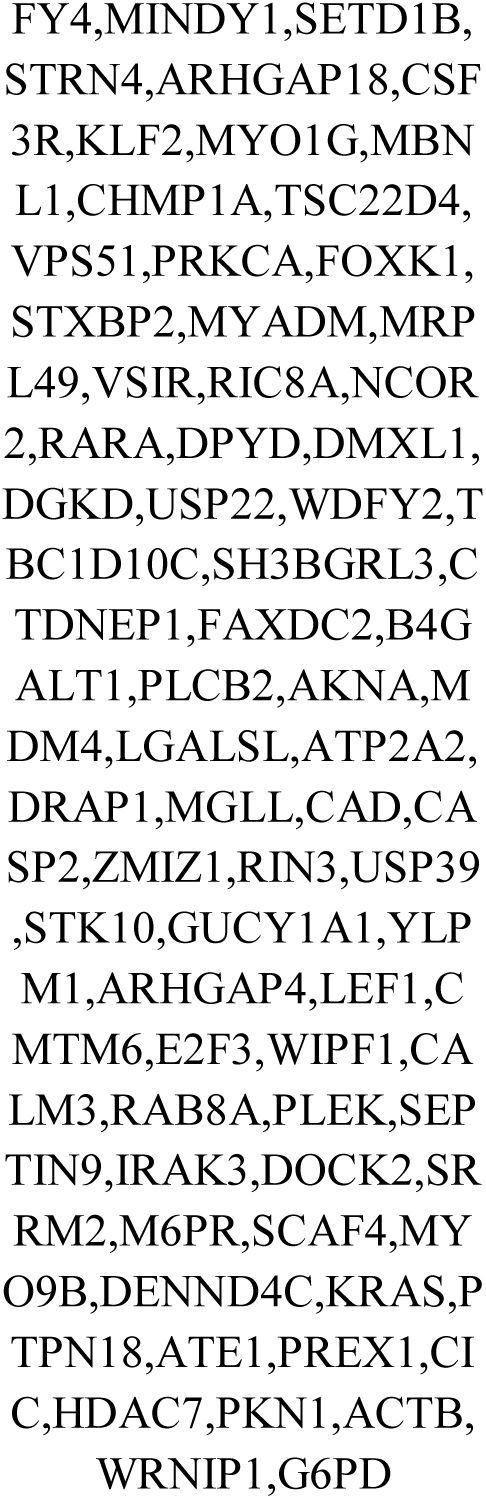
The enriched GO terms of the up and down regulated differentially expressed genes

**Table 3.**
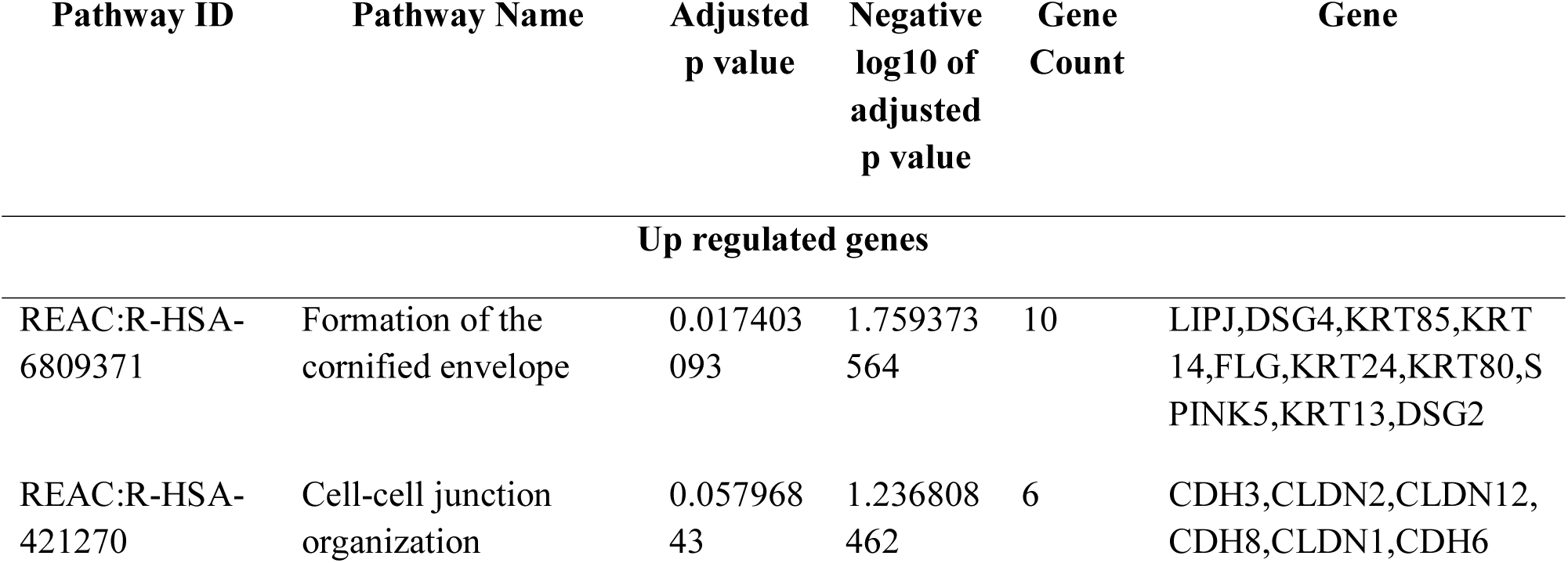
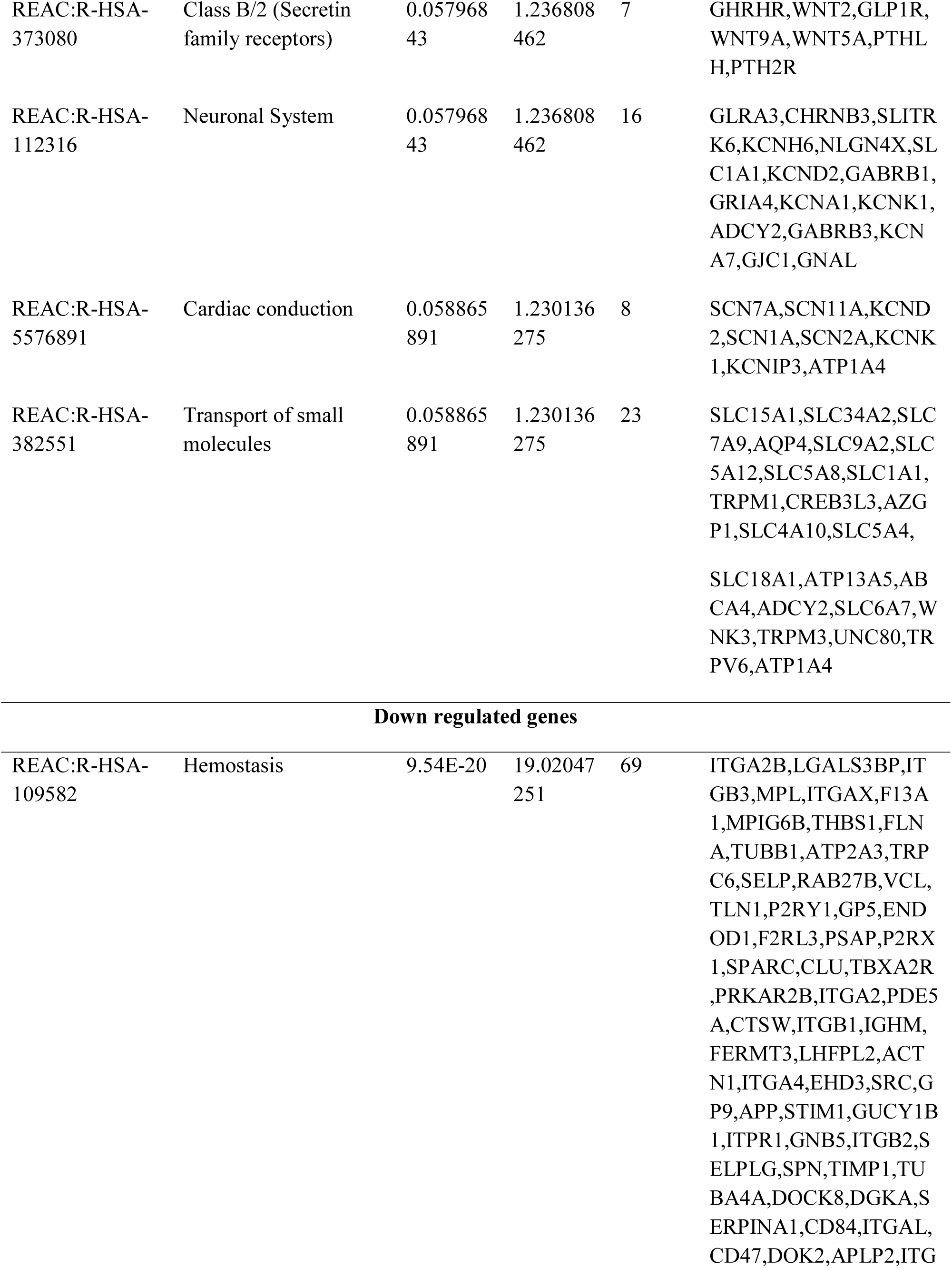
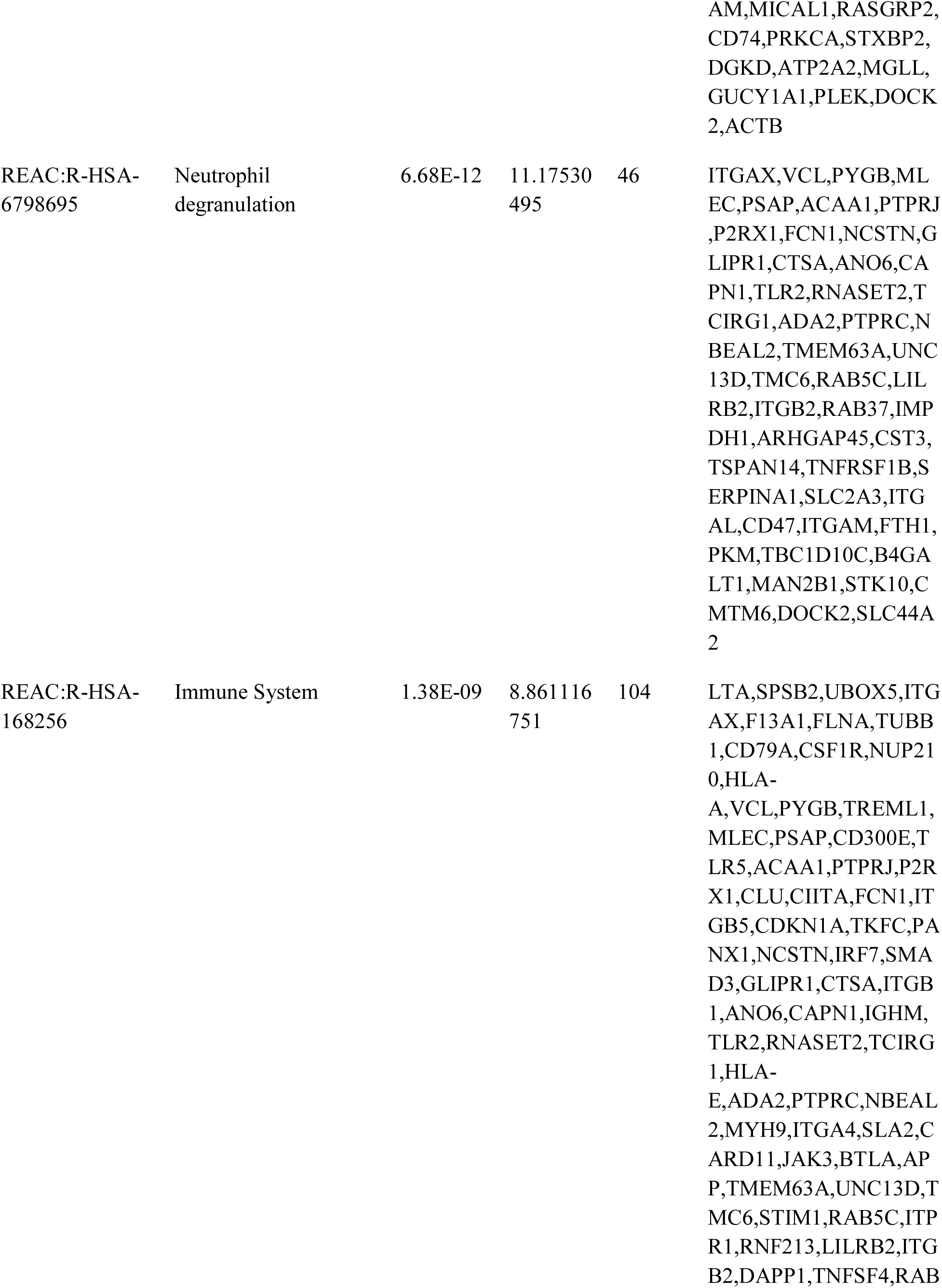
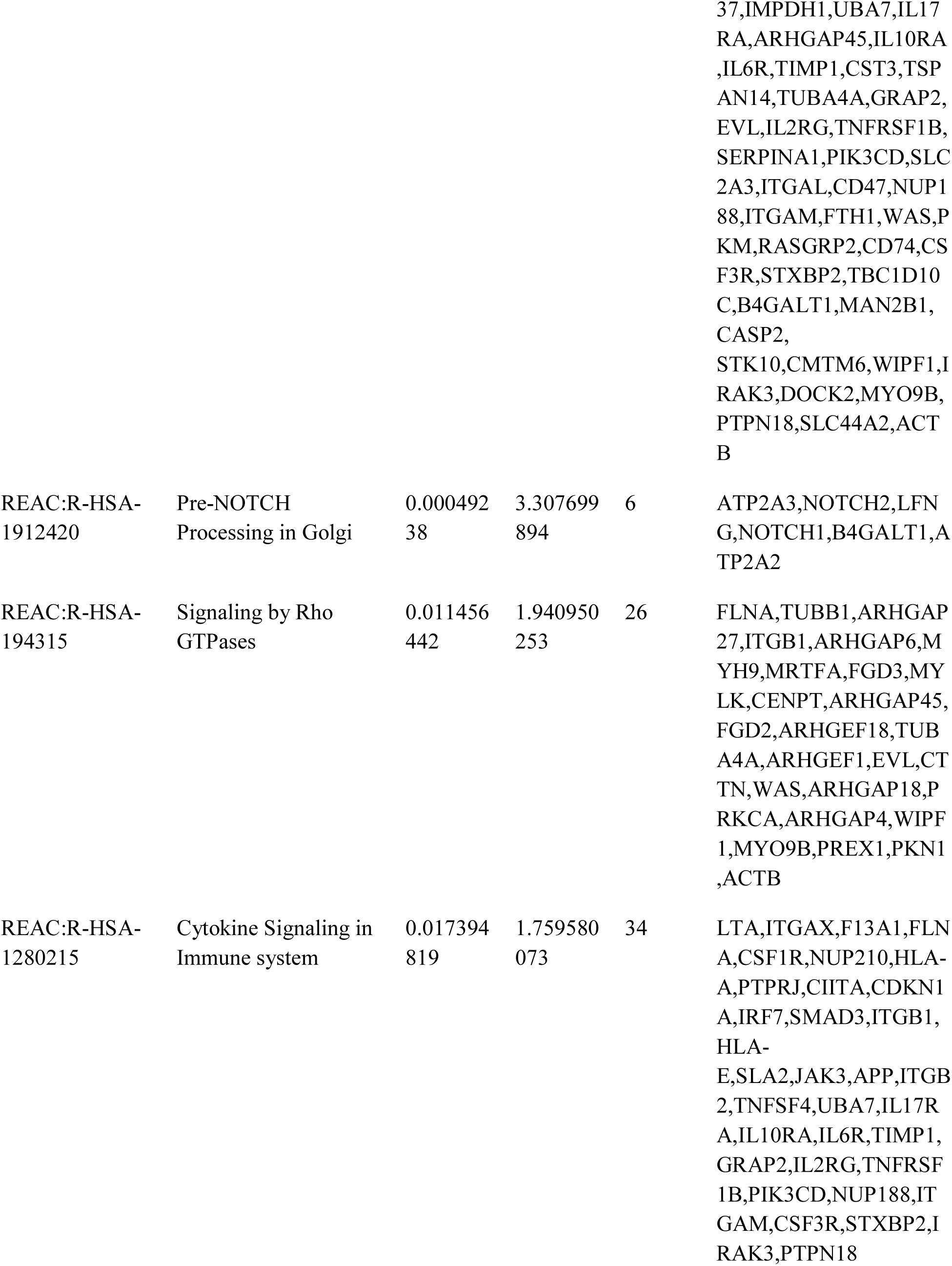
The enriched pathway terms of the up and down regulated differentially expressed genes

| Pathway ID | Pathway Name | Adjusted<br>p value | Negative<br>log10 of<br>adjusted<br>p value | Gene<br>Count | Gene |
| --- | --- | --- | --- | --- | --- |
| Up regulated genes |  |  |  |  |  |
| REAC:R-HSA-6809371 | Formation of the cornified envelope | 0.017403<br>093 | 1.759373<br>564 | 10 | LIPJ,DSG4,KRT85,KRT14,FLG,KRT24,KRT80,S<br>PINK5,KRT13,DSG2 |
| REAC:R-HSA-421270 | Cell-cell junction organization | 0.057968<br>43 | 1.236808<br>462 | 6 | CDH3,CLDN2,CLDN12,<br>CDH8,CLDN1,CDH6 |
| REAC:R-HSA-373080 | Class B/2 (Secretin family receptors) | 0.057968<br>43 | 1.236808<br>462 | 7 | GHRHR,WNT2,GLP1R,WNT9A,WNT5A,PTHLH,PTH2R |
| REAC:R-HSA-112316 | Neuronal System | 0.057968<br>43 | 1.236808<br>462 | 16 | GLRA3,CHRNA3,SLITRK6,KCNH6,NLGN4X,SLC1A1,KCND2,GABRB1,GRIA4,KCNA1,KCNK1,ADCY2,GABRB3,KCNA7,GJC1,GNAL |
| REAC:R-HSA-5576891 | Cardiac conduction | 0.058865<br>891 | 1.230136<br>275 | 8 | SCN7A,SCN11A,KCND2,SCN1A,SCN2A,KCNK1,KCNIP3,ATP1A4 |
| REAC:R-HSA-382551 | Transport of small molecules | 0.058865<br>891 | 1.230136<br>275 | 23 | SLC15A1,SLC34A2,SLC7A9,AQP4,SLC9A2,SLC5A12,SLC5A8,SLC1A1,TRPM1,CREB3L3,AZGP1,SLC4A10,SLC5A4,SLC18A1,ATP13A5,ABCA4,ADCY2,SLC6A7,WNK3,TRPM3,UNC80,TRPV6,ATP1A4 |
| **Down regulated genes** |  |  |  |  |  |
| REAC:R-HSA-109582 | Hemostasis | 9.54E-20 | 19.02047<br>251 | 69 | ITGA2B,LGALS3BP,ITGB3,MPL,ITGAX,F13A1,MPIG6B,THBS1,FLNA,TUBB1,ATP2A3,TRPC6,SELP,RAB27B,VCL,TLN1,P2RY1,GP5,ENDO1,F2RL3,PSAP,P2RX1,SPARC,CLU,TBXA2R,PRKAR2B,ITGA2,PDE5A,CTSW,ITGB1,IGHM,FERMT3,LHFPL2,ACTN1,ITGA4,EHD3,SRC,GP9,APP,STIM1,GUCY1B1,ITPR1,GNB5,ITGB2,SELPLG,SPN,TIMP1,TUBA4A,DOCK8,DGKA,SERPINA1,CD84,ITGAL,CD47,DOK2,APLP2,ITG |
|  |  |  |  |  | AM,MICAL1,RASGRP2,CD74,PRKCA,STXBP2,DGKD,ATP2A2,MGLL,GUCY1A1,PLEK,DOCK2,ACTB |
| REAC:R-HSA-6798695 | Neutrophil degranulation | 6.68E-12 | 11.17530495 | 46 | ITGAX,VCL,PYGB,MLEC,PSAP,ACAA1,PTPRJ,P2RX1,FCN1,NCSTN,GLIPR1,CTSA,ANO6,CAPN1,TLR2,RNASET2,TCIRG1,ADA2,PTPRC,NBEAL2,TMEM63A,UNC13D,TMC6,RAB5C,LILRB2,ITGB2,RAB37,IMPDH1,ARHGAP45,CST3,TSPAN14,TNFRSF1B,SERPINA1,SLC2A3,ITGAL,CD47,ITGAM,FTH1,PKM,TBC1D10C,B4GALT1,MAN2B1,STK10,CDMTM6,DOCK2,SLC44A2 |
| REAC:R-HSA-168256 | Immune System | 1.38E-09 | 8.861116751 | 104 | LTA,SPSB2,UBOX5,ITGAX,F13A1,FLNA,TUBB1,CD79A,CSF1R,NUP210,HLA-A,VCL,PYGB,TREML1,MLEC,PSAP,CD300E,TLR5,ACAA1,PTPRJ,P2RX1,CLU,CIITA,FCN1,ITGB5,CDKN1A,TKFC,PANX1,NCSTN,IRF7,SMAAD3,GLIPR1,CTSA,ITGB1,ANO6,CAPN1,IGHM,TLR2,RNASET2,TCIRG1,HLA-E,ADA2,PTPRC,NBEAL2,MYH9,ITGA4,SLA2,CARD11,JAK3,BTLA,APP,TMEM63A,UNC13D,TMC6,STIM1,RAB5C,ITPR1,RNF213,LILRB2,ITGB2,DAPPI,TNFSF4,RAB |
|  |  |  |  |  | 37,IMPDH1,UBA7,IL17<br>RA,ARHGAP45,IL10RA<br>,IL6R,TIMP1,CST3,TSP<br>AN14,TUBA4A,GRAP2,<br>EVL,IL2RG,TNFRSF1B,<br>SERPINA1,PIK3CD,SLC<br>2A3,ITGAL,CD47,NUP1<br>88,ITGAM,FTH1,WAS,P<br>KM,RASGRP2,CD74,CS<br>F3R,STXBP2,TBC1D10<br>C,B4GALT1,MAN2B1,<br>CASP2,<br>STK10,CMTM6,WIPF1,I<br>RAK3,DOCK2,MYO9B,<br>PTPN18,SLC44A2,ACT<br>B |
| REAC:R-HSA-1912420 | Pre-NOTCH<br>Processing in Golgi | 0.000492<br>38 | 3.307699<br>894 | 6 | ATP2A3,NOTCH2,LFN<br>G,NOTCH1,B4GALT1,A<br>TP2A2 |
| REAC:R-HSA-194315 | Signaling by Rho<br>GTPases | 0.011456<br>442 | 1.940950<br>253 | 26 | FLNA,TUBB1,ARHGAP<br>27,ITGB1,ARHGAP6,M<br>YH9,MRTFA,FGD3,MY<br>LK,CENPT,ARHGAP45,<br>FGD2,ARHGEF18,TUB<br>A4A,ARHGEF1,EVL,CT<br>TN,WAS,ARHGAP18,P<br>RKCA,ARHGAP4,WIPF<br>1,MYO9B,PREX1,PKN1<br>,ACTB |
| REAC:R-HSA-1280215 | Cytokine Signaling in<br>Immune system | 0.017394<br>819 | 1.759580<br>073 | 34 | LTA,ITGAX,F13A1,FLN<br>A,CSF1R,NUP210,HLA-A,PTPRJ,CIITA,CDKN1<br>A,IRF7,SMAD3,ITGB1,<br>HLA-E,SLA2,JAK3,APP,ITGB<br>2,TNFSF4,UBA7,IL17R<br>A,IL10RA,IL6R,TIMP1,<br>GRAP2,IL2RG,TNFRSF<br>1B,PIK3CD,NUP188,IT<br>GAM,CSF3R,STXBP2,I<br>RAK3,PTPN18 |

### Construction of the PPI network and module analysis

The PPI network of DEGs was constructed with 8472 nodes and 20624 edges (Fig.3). The node degree, betweenness centrality, stress centrality and closeness centrality methods were performed to calculate the top hub genes and are listed in Table 4. The results revealed 5 up and 5 down regulated genes identified as hub genes, including TRIM54, ELAVL2, PTN, KIT, BMPR1B, APP, SRC, ITGA4, RPA1 and ACTB. PEWCC1 was used to identify the significant modules in the PPI network and the two significant modules were selected. Module 1 consisted of 95 nodes and 103 edges (Fig.4A) and module 2 consisted of 79 nodes and 131 edges (Fig.4B). Following GO and pathway enrichment analysis, the module 1 was revealed to be associated with multicellular organismal process, developmental process and intrinsic component of membrane, and module 2 was revealed to be associated with hemostasis, immune system, enzyme binding, cytokine signaling in immune system and protein binding.

**Fig. 3.**
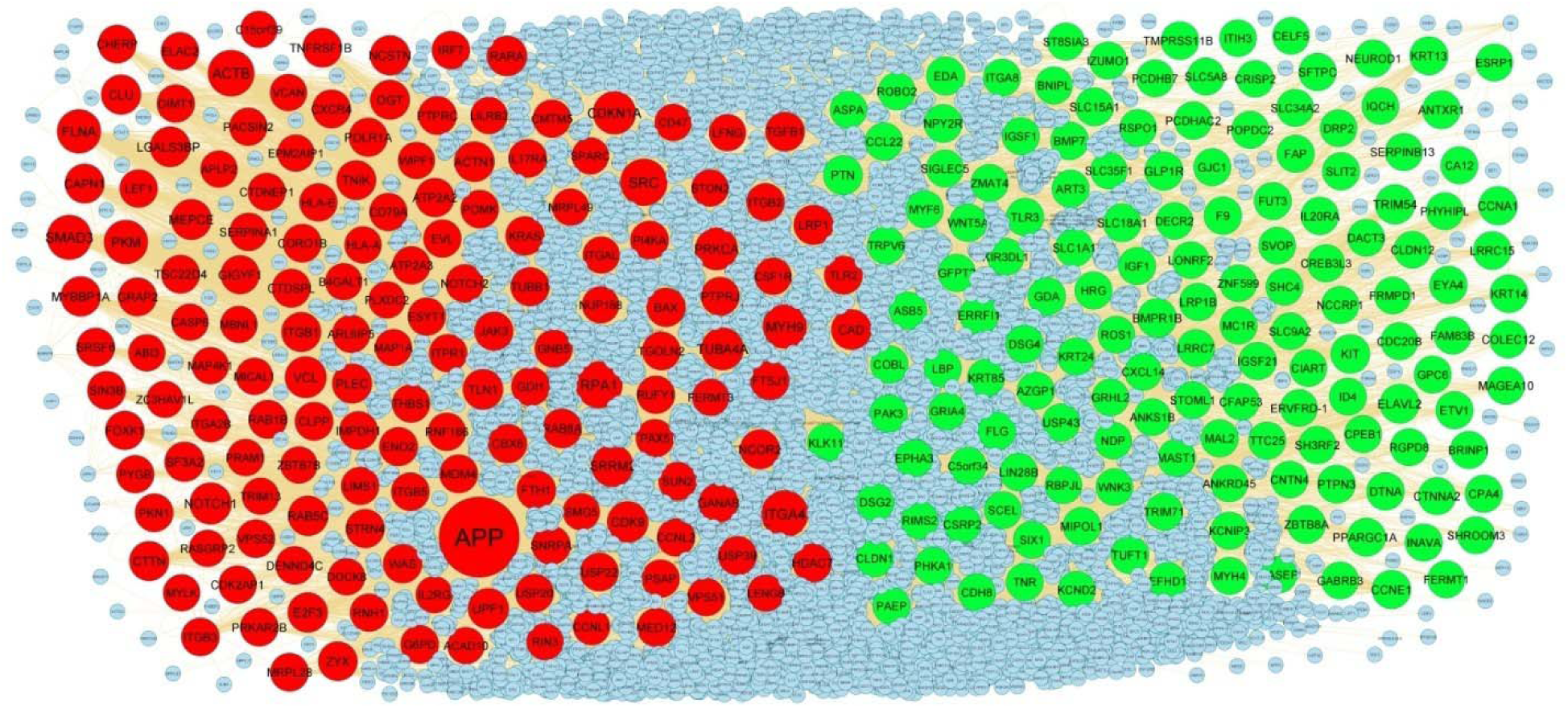
PPI network of DEGs. Up regulated genes are marked in green; down regulated genes are marked in red

**Fig. 4.**
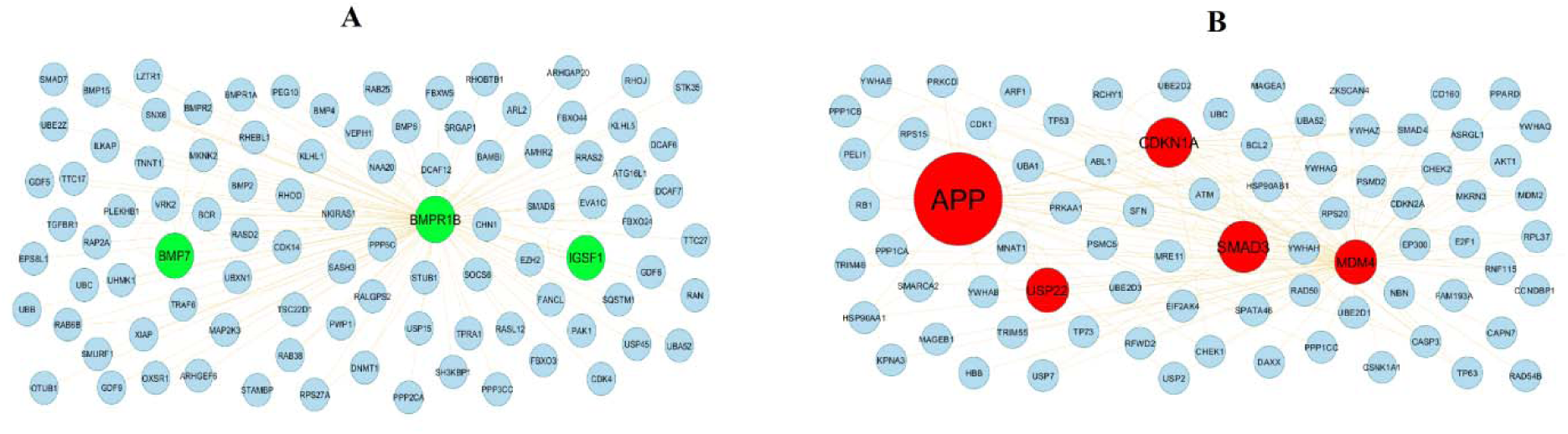
Modules of isolated form PPI of DEGs. (A) The most significant module was obtained from PPI network with 95 nodes and 103 edges for up regulated genes (B) The most significant module was obtained from PPI network with 79 nodes and 131 edges for down regulated genes. Up regulated genes are marked in green; down regulated genes are marked in red

**Table 4.** Topology table for up and down regulated genes.

| **Regulation** | **Node** | **Degree** | **Betweenness** | **Stress** | **Closeness** |
| --- | --- | --- | --- | --- | --- |
| Up | TRIM54 | 133 | 0.012147 | 21844832 | 0.322581 |
| Up | ELAVL2 | 118 | 0.009706 | 13548160 | 0.323122 |
| Up | PTN | 109 | 0.008667 | 10295782 | 0.325332 |
| Up | KIT | 105 | 0.005763 | 7684058 | 0.332132 |
| Up | BMPR1B | 102 | 0.012139 | 7911102 | 0.352362 |
| Up | ERRFI1 | 99 | 0.008331 | 11254818 | 0.338466 |
| Up | CCNE1 | 86 | 0.003025 | 5367340 | 0.330825 |
| Up | CCNA1 | 83 | 0.004101 | 5615582 | 0.34443 |
| Up | GLP1R | 78 | 0.010843 | 23727352 | 0.299903 |
| Up | EDA | 74 | 0.008686 | 5992546 | 0.315667 |
| Up | PPARGC1A | 72 | 0.002826 | 4898866 | 0.322485 |
| Up | KRT13 | 70 | 0.004347 | 10153884 | 0.31577 |
| Up | SLC15A1 | 64 | 0.00578 | 5083164 | 0.301448 |
| Up | ANTXR1 | 59 | 0.006338 | 4754724 | 0.364739 |
| Up | ZBTB8A | 59 | 0.00408 | 8024144 | 0.307376 |
| Up | KRT14 | 57 | 0.001607 | 3870640 | 0.322978 |
| Up | NPY2R | 57 | 0.004358 | 4577120 | 0.302607 |
| Up | SLC18A1 | 56 | 0.005946 | 4356010 | 0.306766 |
| Up | LRRC15 | 54 | 0.005085 | 12368332 | 0.298816 |
| Up | PTPN3 | 53 | 0.00338 | 4379752 | 0.322377 |
| Up | TMPRSS11B | 53 | 0.005232 | 13231264 | 0.28515 |
| Up | TUFT1 | 50 | 0.002391 | 2006556 | 0.308054 |
| Up | BMP7 | 48 | 0.004125 | 3406724 | 0.310316 |
| Up | MIPOL1 | 47 | 0.003199 | 7831728 | 0.296433 |
| Up | DSG2 | 43 | 0.00103 | 1538692 | 0.326669 |
| Up | MAGEA10 | 42 | 0.002659 | 2782598 | 0.311971 |
| Up | TLR3 | 41 | 0.002523 | 2126014 | 0.311365 |
| Up | MAST1 | 41 | 0.002679 | 1449942 | 0.345502 |
| Up | F9 | 40 | 0.003205 | 3436702 | 0.301448 |
| Up | DTNA | 39 | 0.003643 | 2835460 | 0.311825 |
| Up | DSG4 | 39 | 0.002547 | 6333108 | 0.286045 |
| Up | LIN28B | 38 | 0.002233 | 2392960 | 0.306429 |
| Up | EPHA3 | 38 | 0.003012 | 1844952 | 0.317553 |
| Up | GDA | 38 | 0.002698 | 4154080 | 0.301995 |
| Up | IL20RA | 37 | 0.003001 | 2163774 | 0.306885 |
| Up | EYA4 | 36 | 0.002123 | 2202158 | 0.300142 |
| Up | CCL22 | 35 | 0.003053 | 4599286 | 0.300548 |
| Up | SLC1A1 | 35 | 0.004187 | 3161236 | 0.346317 |
| Up | PCDHAC2 | 35 | 0.004863 | 3849886 | 0.292706 |
| Up | IGSF21 | 34 | 0.002215 | 2796766 | 0.296119 |
| Up | SH3RF2 | 34 | 0.001943 | 1330190 | 0.342509 |
| Up | USP43 | 34 | 0.00183 | 2158278 | 0.305694 |
| Up | COBL | 33 | 7.34E-04 | 789636 | 0.331775 |
| Up | FLG | 33 | 0.001565 | 2103066 | 0.312027 |
| Up | SERPINB13 | 33 | 0.002022 | 2211972 | 0.303009 |
| Up | COLEC12 | 32 | 0.002562 | 1515874 | 0.335126 |
| Up | GFPT2 | 30 | 0.002137 | 2091622 | 0.31094 |
| Up | SFTPC | 30 | 0.002498 | 2902984 | 0.304276 |
| Up | LRP1B | 29 | 0.003228 | 1563906 | 0.344553 |
| Up | AZGP1 | 29 | 0.001511 | 2326868 | 0.30975 |
| Up | ETV1 | 28 | 6.64E-04 | 1570448 | 0.306701 |
| Up | CPA4 | 27 | 0.001782 | 1645444 | 0.303476 |
| Up | KLK11 | 27 | 0.002312 | 3332010 | 0.2924 |
| Up | CTNNA2 | 27 | 0.001317 | 2711166 | 0.312714 |
| Up | WNT5A | 27 | 0.002257 | 1865676 | 0.296525 |
| Up | KRT85 | 27 | 0.001319 | 2630340 | 0.305975 |
| Up | FAP | 27 | 0.001924 | 3392714 | 0.291986 |
| Up | ROBO2 | 26 | 0.001435 | 2365116 | 0.291651 |
| Up | KCNIP3 | 26 | 0.001472 | 3374664 | 0.28202 |
| Up | BRINP1 | 25 | 0.0012 | 2988948 | 0.301301 |
| Up | NCCRP1 | 25 | 8.25E-04 | 1228816 | 0.315173 |
| Up | ART3 | 24 | 0.001611 | 3187858 | 0.283016 |
| Up | INAVA | 24 | 0.00154 | 1886482 | 0.291278 |
| Up | CREB3L3 | 24 | 0.001636 | 2012464 | 0.302332 |
| Up | SLIT2 | 23 | 0.001176 | 1918318 | 0.301259 |
| Up | POPDC2 | 23 | 0.002173 | 1488558 | 0.271797 |
| Up | IZUMO1 | 23 | 0.002732 | 4118726 | 0.259647 |
| Up | MAL2 | 23 | 0.001473 | 2898432 | 0.296261 |
| Up | RGPD8 | 22 | 0.001517 | 1034252 | 0.305586 |
| Up | PAEP | 22 | 0.001035 | 874582 | 0.343828 |
| Up | ASB5 | 22 | 4.24E-04 | 368080 | 0.303583 |
| Up | FAM83B | 22 | 8.25E-04 | 1549462 | 0.305166 |
| Up | SLC5A8 | 22 | 0.001364 | 1608962 | 0.276598 |
| Up | SIX1 | 21 | 0.00131 | 1136652 | 0.295735 |
| Up | KCND2 | 21 | 0.002331 | 1658754 | 0.303956 |
| Up | MYH4 | 21 | 9.70E-04 | 980960 | 0.299623 |
| Up | NEUROD1 | 21 | 0.001441 | 2678376 | 0.287985 |
| Up | PAK3 | 20 | 6.35E-05 | 179392 | 0.297748 |
| Up | TACC2 | 20 | 0.001037 | 1183890 | 0.309374 |
| Up | TTC25 | 20 | 0.001315 | 2071982 | 0.276139 |
| Up | LBP | 19 | 0.001144 | 837746 | 0.309917 |
| Up | HRG | 19 | 8.05E-04 | 1333666 | 0.292607 |
| Up | CDC20B | 19 | 0.001052 | 1653682 | 0.279211 |
| Up | TNR | 19 | 0.001002 | 1524772 | 0.286092 |
| Up | SHROOM3 | 18 | 6.22E-04 | 712586 | 0.316393 |
| Up | PHKA1 | 18 | 8.26E-04 | 664362 | 0.301763 |
| Up | RIMS2 | 18 | 8.17E-04 | 1001820 | 0.305575 |
| Up | CSRP2 | 18 | 2.40E-04 | 457510 | 0.304983 |
| Up | KRT24 | 18 | 2.78E-05 | 166448 | 0.30059 |
| Up | LRRC7 | 18 | 6.50E-04 | 1043008 | 0.299085 |
| Up | ITGA8 | 17 | 0.001062 | 974108 | 0.291524 |
| Up | SIGLEC5 | 17 | 0.001041 | 718852 | 0.290351 |
| Up | CLDN1 | 17 | 7.91E-04 | 1036670 | 0.284009 |
| Up | IGF1 | 17 | 0.001485 | 2375614 | 0.275937 |
| Up | MYF6 | 17 | 3.75E-04 | 472100 | 0.284503 |
| Up | CDH8 | 17 | 7.20E-04 | 923698 | 0.293351 |
| Up | TRIM71 | 17 | 9.54E-04 | 1482892 | 0.250325 |
| Up | ESRP1 | 16 | 9.05E-04 | 638852 | 0.286983 |
| Up | GPC6 | 16 | 0.001651 | 1480508 | 0.325674 |
| Up | CPEB1 | 16 | 0.001555 | 915000 | 0.327906 |
| Up | GRIA4 | 16 | 3.00E-04 | 243164 | 0.303519 |
| Up | ID4 | 16 | 9.00E-04 | 532420 | 0.307977 |
| Up | TRPV6 | 16 | 6.17E-04 | 609086 | 0.281005 |
| Up | GJC1 | 16 | 0.001409 | 1565962 | 0.27087 |
| Up | FERMT1 | 16 | 4.60E-04 | 808642 | 0.300267 |
| Up | EFHD1 | 16 | 9.97E-04 | 2285970 | 0.281553 |
| Up | ZMAT4 | 15 | 0.00194 | 2108242 | 0.331141 |
| Up | FUT3 | 15 | 0.001643 | 1729770 | 0.26295 |
| Up | KIR3DL1 | 15 | 6.23E-04 | 1269024 | 0.278188 |
| Up | MC1R | 15 | 0.001048 | 1221034 | 0.282739 |
| Up | BNIP1 | 15 | 9.69E-04 | 1242002 | 0.277315 |
| Up | PHYHIP1 | 15 | 4.86E-04 | 372398 | 0.288465 |
| Up | PCDHB7 | 15 | 0.00158 | 1015798 | 0.272361 |
| Up | IGSF1 | 14 | 7.87E-04 | 482056 | 0.268635 |
| Up | ST8SIA3 | 14 | 8.32E-04 | 665984 | 0.291543 |
| Up | CA12 | 14 | 6.44E-05 | 102792 | 0.268619 |
| Up | SCEL | 14 | 7.34E-04 | 388298 | 0.33479 |
| Up | SLC34A2 | 14 | 0.001583 | 1563014 | 0.247542 |
| Up | GABRB3 | 14 | 5.08E-04 | 274876 | 0.300924 |
| Up | ANKS1B | 4 | 4.96E-05 | 207240 | 0.331775 |
| Up | CIART | 2 | 5.70E-05 | 275826 | 0.319652 |
| Up | IQCH | 2 | 0 | 0 | 0.31964 |
| Up | CXCL14 | 2 | 3.86E-05 | 97804 | 0.322749 |
| Up | LONRF2 | 1 | 0 | 0 | 0.2311 |
| Up | ITIH3 | 1 | 0 | 0 | 0.243265 |
| Up | DACT3 | 1 | 0 | 0 | 0.240369 |
| Up | DECR2 | 1 | 0 | 0 | 0.317752 |
| Up | CFAP53 | 1 | 0 | 0 | 0.317752 |
| Up | NDP | 1 | 0 | 0 | 0.317752 |
| Up | ANKRD45 | 1 | 0 | 0 | 0.317752 |
| Up | CNTN4 | 1 | 0 | 0 | 0.317752 |
| Up | GRHL2 | 1 | 0 | 0 | 0.317752 |
| Up | STOML1 | 1 | 0 | 0 | 0.317752 |
| Up | RBPJL | 1 | 0 | 0 | 0.317752 |
| Up | ERVFRD-1 | 1 | 0 | 0 | 0.317752 |
| Up | SHC4 | 1 | 0 | 0 | 0.249331 |
| Up | SLC9A2 | 1 | 0 | 0 | 0.28788 |
| Up | RASEF | 1 | 0 | 0 | 0.247252 |
| Up | SVOP | 1 | 0 | 0 | 0.230718 |
| Up | SLC35F1 | 1 | 0 | 0 | 0.232315 |
| Up | CELF5 | 1 | 0 | 0 | 0.244219 |
| Up | ZNF599 | 1 | 0 | 0 | 0.232066 |
| Up | C5orf34 | 1 | 0 | 0 | 0.240823 |
| Up | ROS1 | 1 | 0 | 0 | 0.279842 |
| Up | CRISP2 | 1 | 0 | 0 | 0.269028 |
| Up | ASPA | 1 | 0 | 0 | 0.239909 |
| Up | CLDN12 | 1 | 0 | 0 | 0.246359 |
| Up | RSPO1 | 1 | 0 | 0 | 0.256266 |
| Up | WNK3 | 1 | 0 | 0 | 0.247598 |
| Up | FRMPD1 | 1 | 0 | 0 | 0.260037 |
| Up | DRP2 | 1 | 0 | 0 | 0.23771 |
| Down | APP | 2189 | 0.002666 | 3365310 | 0.317845 |
| Down | SRC | 504 | 0.400224 | 5.59E+08 | 0.465717 |
| Down | ITGA4 | 496 | 0.008595 | 8040232 | 0.303306 |
| Down | RPA1 | 493 | 0 | 0 | 0.2311 |
| Down | ACTB | 476 | 0.01213 | 21489770 | 0.335607 |
| Down | SMAD3 | 398 | 0.010658 | 16678612 | 0.357999 |
| Down | MYH9 | 364 | 0.006158 | 6598872 | 0.337227 |
| Down | FLNA | 353 | 0.007077 | 11964634 | 0.337885 |
| Down | PKM | 353 | 4.21E-05 | 68424 | 0.292597 |
| Down | CDKN1A | 302 | 0.010037 | 16637086 | 0.367027 |
| Down | TUBA4A | 281 | 0.004054 | 3547868 | 0.316706 |
| Down | PRKCA | 279 | 4.19E-05 | 33780 | 0.274616 |
| Down | SRRM2 | 259 | 0.022533 | 38294108 | 0.381347 |
| Down | NOTCH1 | 245 | 0.013646 | 27956328 | 0.345433 |
| Down | VCL | 233 | 0.005656 | 4112886 | 0.355311 |
| Down | CDK9 | 211 | 0.031272 | 43724982 | 0.387802 |
| Down | UPF1 | 199 | 0.048906 | 1.46E+08 | 0.371865 |
| Down | MEPCE | 186 | 0.005625 | 5736504 | 0.361481 |
| Down | LGALS3BP | 176 | 0.050731 | 56826626 | 0.397302 |
| Down | ACTN1 | 172 | 0.005719 | 8627714 | 0.316335 |
| Down | CAD | 171 | 0.010289 | 9147932 | 0.323472 |
| Down | TLN1 | 170 | 0.024835 | 32927066 | 0.377671 |
| Down | RARA | 168 | 0.016376 | 38790824 | 0.34524 |
| Down | RAB5C | 164 | 0.009051 | 26501322 | 0.32746 |
| Down | E2F3 | 157 | 0.067172 | 81164158 | 0.404238 |
| Down | MYBBP1A | 157 | 0.004253 | 4853872 | 0.319216 |
| Down | SF3A2 | 153 | 0.005345 | 5537860 | 0.321455 |
| Down | NCSTN | 148 | 0.013675 | 11620366 | 0.373614 |
| Down | PLEC | 148 | 0.007198 | 7735012 | 0.333667 |
| Down | CLU | 148 | 0.004284 | 4921862 | 0.315621 |
| Down | CTTN | 144 | 1.85E-05 | 34480 | 0.273479 |
| Down | LRP1 | 143 | 0.035606 | 66738612 | 0.368025 |
| Down | ITGB1 | 141 | 0.007016 | 11423404 | 0.330433 |
| Down | OGT | 139 | 0.017531 | 34670330 | 0.363911 |
| Down | NCOR2 | 136 | 0.05242 | 67476208 | 0.414314 |
| Down | TNIK | 133 | 0.003794 | 7500172 | 0.323255 |
| Down | CAPN1 | 132 | 0.011955 | 9201942 | 0.362783 |
| Down | TNFRSF1B | 126 | 0.005235 | 6048582 | 0.338347 |
| Down | RAB8A | 126 | 0.033442 | 45967308 | 0.398711 |
| Down | ZYX | 122 | 0.002615 | 4159134 | 0.316671 |
| Down | TGFB1 | 121 | 0.030548 | 55611552 | 0.380726 |
| Down | GANAB | 121 | 0.004766 | 9084764 | 0.316324 |
| Down | FTH1 | 112 | 0 | 0 | 0.240315 |
| Down | FTSJ1 | 110 | 0.008563 | 21677110 | 0.350069 |
| Down | WAS | 105 | 0.006824 | 14951804 | 0.316416 |
| Down | ATP2A2 | 104 | 0.005502 | 13512860 | 0.325454 |
| Down | BAX | 103 | 0.002749 | 7775802 | 0.312997 |
| Down | SNRPA | 103 | 0.004686 | 7615722 | 0.323146 |
| Down | FOXK1 | 103 | 6.56E-05 | 124624 | 0.297523 |
| Down | CBX6 | 102 | 2.60E-05 | 130320 | 0.300101 |
| Down | HDAC7 | 100 | 7.36E-06 | 41282 | 0.289409 |
| Down | KRAS | 100 | 0.00505 | 3946654 | 0.314372 |
| Down | TUBB1 | 99 | 0.010762 | 7334820 | 0.356496 |
| Down | USP39 | 99 | 0.004163 | 4124064 | 0.333346 |
| Down | STRN4 | 98 | 0.015639 | 13083920 | 0.371546 |
| Down | GDI1 | 97 | 0.00808 | 7245854 | 0.355151 |
| Down | POLR1A | 95 | 0.004493 | 3158464 | 0.315356 |
| Down | GIGYF1 | 94 | 0.00389 | 8518854 | 0.317868 |
| Down | MED12 | 94 | 1.58E-05 | 10046 | 0.275586 |
| Down | JAK3 | 93 | 0.011564 | 8350784 | 0.351704 |
| Down | ENO2 | 92 | 0 | 0 | 0.260201 |
| Down | RAB1B | 89 | 0.008892 | 17406094 | 0.336899 |
| Down | CTDSPL | 86 | 0.005896 | 10405078 | 0.326792 |
| Down | GRAP2 | 84 | 0.010691 | 20962628 | 0.34186 |
| Down | TSC22D4 | 84 | 0.008449 | 16411146 | 0.350849 |
| Down | PTPRJ | 83 | 0.003374 | 6154730 | 0.317449 |
| Down | CXCR4 | 82 | 0 | 0 | 0.319392 |
| Down | MDM4 | 81 | 0 | 0 | 0.321014 |
| Down | SRSF6 | 81 | 0.00369 | 2617526 | 0.348744 |
| Down | IMPDH1 | 79 | 0.008468 | 14872056 | 0.324636 |
| Down | PTPRC | 78 | 0 | 0 | 0.245083 |
| Down | CLPP | 78 | 0 | 0 | 0.317752 |
| Down | TLR2 | 77 | 4.79E-05 | 76992 | 0.322317 |
| Down | CORO1B | 77 | 0 | 0 | 0.317752 |
| Down | USP22 | 76 | 0 | 0 | 0.317752 |
| Down | SIN3B | 76 | 0.006673 | 5072566 | 0.357084 |
| Down | SUN2 | 75 | 0 | 0 | 0.319369 |
| Down | PI4KA | 75 | 5.09E-05 | 194840 | 0.320231 |
| Down | MRPL28 | 75 | 0 | 0 | 0.317752 |
| Down | IL17RA | 75 | 0 | 0 | 0.317752 |
| Down | PKN1 | 74 | 0 | 0 | 0.317752 |
| Down | USP20 | 74 | 0.003156 | 4712314 | 0.357777 |
| Down | CHERP | 73 | 1.58E-04 | 227708 | 0.320717 |
| Down | VPS52 | 73 | 5.49E-05 | 172518 | 0.322077 |
| Down | TGOLN2 | 71 | 0 | 0 | 0.317752 |
| Down | G6PD | 70 | 9.99E-05 | 528874 | 0.329578 |
| Down | POMK | 69 | 0 | 0 | 0.317752 |
| Down | DENND4C | 69 | 4.16E-05 | 440608 | 0.329353 |
| Down | PRKAR2B | 69 | 0 | 0 | 0.317752 |
| Down | ITGB3 | 66 | 0 | 0 | 0.317752 |
| Down | PAX5 | 65 | 9.66E-05 | 409222 | 0.327385 |
| Down | ITGB5 | 64 | 0 | 0 | 0.317752 |
| Down | NOTCH2 | 62 | 0.0076 | 4500746 | 0.353023 |
| Down | SERPINA1 | 61 | 0 | 0 | 0.317752 |
| Down | RNF166 | 61 | 0 | 0 | 0.317752 |
| Down | LEF1 | 60 | 0.002734 | 2878552 | 0.356893 |
| Down | RUFY1 | 59 | 0 | 0 | 0.317752 |
| Down | ITPR1 | 59 | 0.016504 | 15009770 | 0.367541 |
| Down | IRF7 | 58 | 4.20E-05 | 254816 | 0.325797 |
| Down | CDK2AP1 | 58 | 0.016675 | 15745286 | 0.373937 |
| Down | THBS1 | 57 | 0 | 0 | 0.317752 |
| Down | HLA-E | 57 | 0.007079 | 6193594 | 0.359694 |
| Down | RNH1 | 56 | 5.89E-05 | 307276 | 0.322725 |
| Down | ELAC2 | 56 | 0.006031 | 7284664 | 0.360863 |
| Down | PACSIN2 | 55 | 0 | 0 | 0.318476 |
| Down | ARL6IP5 | 55 | 0 | 0 | 0.317752 |
| Down | CCNL2 | 55 | 0 | 0 | 0.317752 |
| Down | ATP2A3 | 55 | 0.033293 | 57837312 | 0.388567 |
| Down | DIMT1 | 54 | 0 | 0 | 0.317752 |
| Down | PSAP | 54 | 0 | 0 | 0.317752 |
| Down | CD79A | 54 | 3.27E-04 | 585398 | 0.330711 |
| Down | VPS51 | 54 | 0 | 0 | 0.317752 |
| Down | CTDNEP1 | 53 | 0 | 0 | 0.321443 |
| Down | ITGB2 | 53 | 0.002039 | 3160000 | 0.334208 |
| Down | LIMS1 | 53 | 0.009901 | 17849898 | 0.334376 |
| Down | CMTM5 | 53 | 0 | 0 | 0.279878 |
| Down | GNB5 | 52 | 0.042962 | 1.29E+08 | 0.371403 |
| Down | ESYT1 | 52 | 0.010431 | 19740350 | 0.351404 |
| Down | EVL | 52 | 0.002936 | 3873720 | 0.321861 |
| Down | PYGB | 50 | 0 | 0 | 0.243498 |
| Down | HLA-A | 7 | 3.15E-06 | 4282 | 0.25335 |
| Down | CSF1R | 4 | 0 | 0 | 0.243498 |
| Down | MYLK | 4 | 0.02758 | 53431176 | 0.357349 |
| Down | MICAL1 | 4 | 0 | 0 | 0.263631 |
| Down | ZBTB7B | 3 | 0.011364 | 18936744 | 0.356555 |
| Down | RIN3 | 3 | 0 | 0 | 0.28158 |
| Down | SPARC | 3 | 5.99E-05 | 100996 | 0.279112 |
| Down | LFNG | 3 | 0 | 0 | 0.25378 |
| Down | PRAM1 | 3 | 0.007566 | 10658530 | 0.33669 |
| Down | FERMT3 | 3 | 0.004345 | 6463622 | 0.323183 |
| Down | APLP2 | 3 | 0.002097 | 5599528 | 0.312207 |
| Down | ACAD10 | 3 | 5.54E-06 | 5790 | 0.281452 |
| Down | ITGA2B | 3 | 0.003888 | 5735454 | 0.329328 |
| Down | VCAN | 3 | 0.005452 | 8532800 | 0.331116 |
| Down | LENG8 | 3 | 0.005412 | 6353682 | 0.338466 |
| Down | EPM2AIP1 | 3 | 0.004426 | 3470384 | 0.316764 |
| Down | MAP1A | 3 | 0.00816 | 16680766 | 0.33231 |
| Down | STON2 | 3 | 0.003943 | 9629562 | 0.331737 |
| Down | ABI3 | 3 | 6.24E-05 | 141136 | 0.260663 |
| Down | TRIM13 | 3 | 0.006012 | 17818956 | 0.328279 |
| Down | WIPF1 | 3 | 0.006612 | 10493584 | 0.327423 |
| Down | IL2RG | 3 | 0 | 0 | 0.249331 |
| Down | B4GALT1 | 3 | 0.004053 | 8509232 | 0.32549 |
| Down | DOCK8 | 2 | 0.008903 | 15185354 | 0.333205 |
| Down | NUP188 | 2 | 0.007135 | 7672290 | 0.329065 |
| Down | MAP4K1 | 2 | 0.003653 | 4056316 | 0.316949 |
| Down | PLXDC2 | 2 | 2.45E-06 | 6914 | 0.241879 |
| Down | RASGRP2 | 2 | 0.004447 | 3796740 | 0.322389 |
| Down | CCNL1 | 2 | 0 | 0 | 0.244253 |
| Down | CASP6 | 2 | 3.41E-05 | 75570 | 0.268802 |
| Down | C15orf39 | 2 | 0.008306 | 17301536 | 0.332017 |
| Down | MBNL1 | 2 | 1.36E-05 | 74714 | 0.306896 |
| Down | ZC3HAV1L | 2 | 0.004666 | 6660660 | 0.327695 |
| Down | SMG5 | 2 | 2.15E-05 | 101210 | 0.303529 |
| Down | MRPL49 | 2 | 1.78E-05 | 51682 | 0.289293 |
| Down | LILRB2 | 2 | 0 | 0 | 0.28788 |
| Down | ITGAL | 2 | 0 | 0 | 0.28788 |
| Down | CD47 | 2 | 0 | 0 | 0.28788 |

### MiRNA-hub gene regulatory network construction

To investigate interaction correlations between the hub genes and miRNAs, a miRNA-hub gene interaction network was constructed, which has 2598 nodes (hub genes:307; miRNAs: 2291) and 17035 edges. Results from the current investigation demonstrated that one target gene might be regulated by several miRNAs (Fig. 5) and are listed in Table 5. Up regulated hub genes including, ERRFI1 targeted by 70 miRNAs (ex; hsa-mir-198); CCNE1 targeted by 50 miRNAs (ex; hsa-mir-181b-5p); ELAVL2 targeted by 43 miRNAs (ex; hsa-mir-766-3p); ANTXR1 targeted by 36 miRNAs (ex; hsa-mir-624-5p); ZBTB8A targeted by 34 miRNAs (ex; hsa-mir-10b-5p). Similarly, down regulated hub genes including, SRRM2 targeted by 250 miRNAs (ex; hsa-mir-582-5p); CDKN1A targeted by 187 miRNAs (ex; hsa-mir-657); MYH9 targeted by 126 miRNAs (ex; hsa-mir-1231); APP targeted by 125 miRNAs (ex; hsa-mir-509-3-5p); ACTB targeted by 122 miRNAs (ex; hsa-mir-192-3p).

**Fig. 5.**
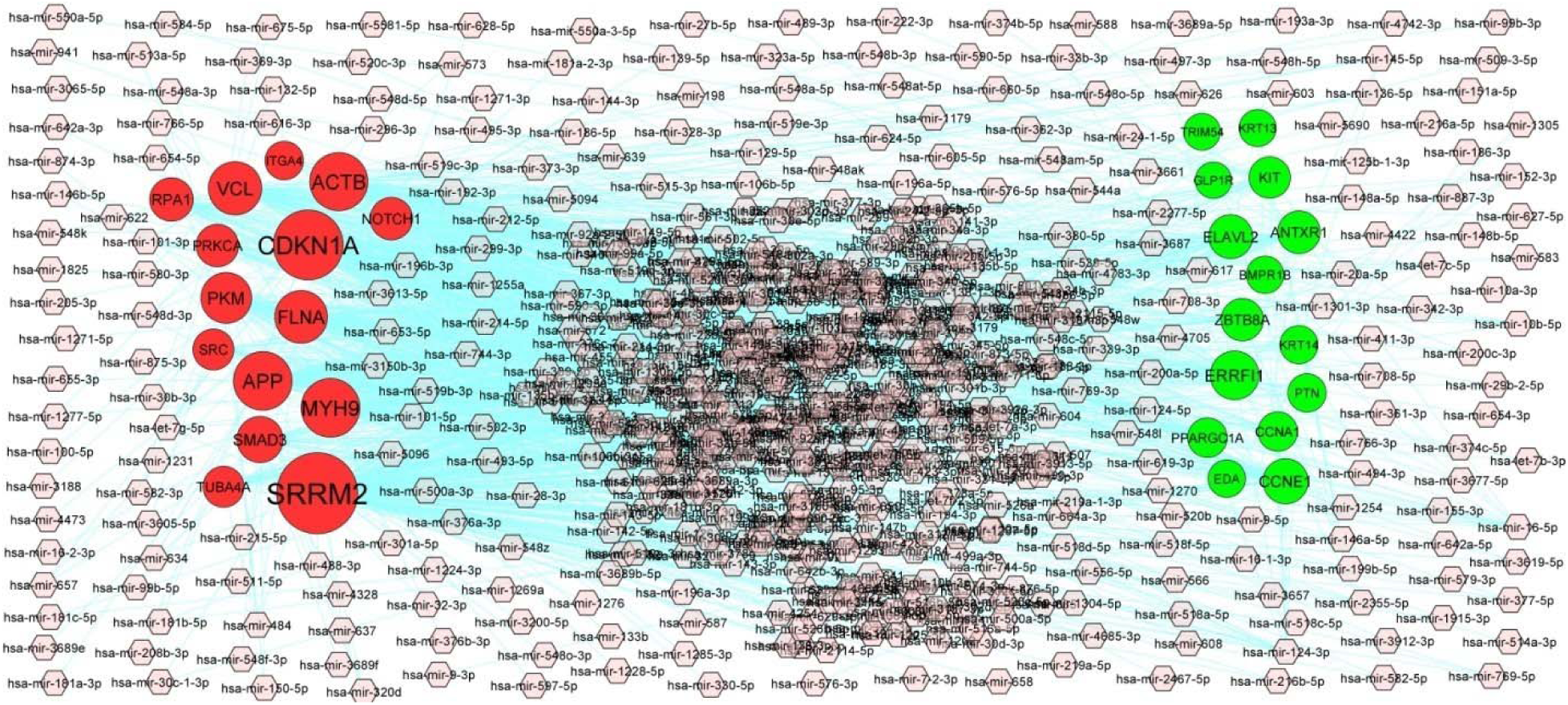
Target gene - miRNA regulatory network between target genes. The blue color diamond nodes represent the key miRNAs; up regulated genes are marked in green; down regulated genes are marked in red.

**Table 5.**
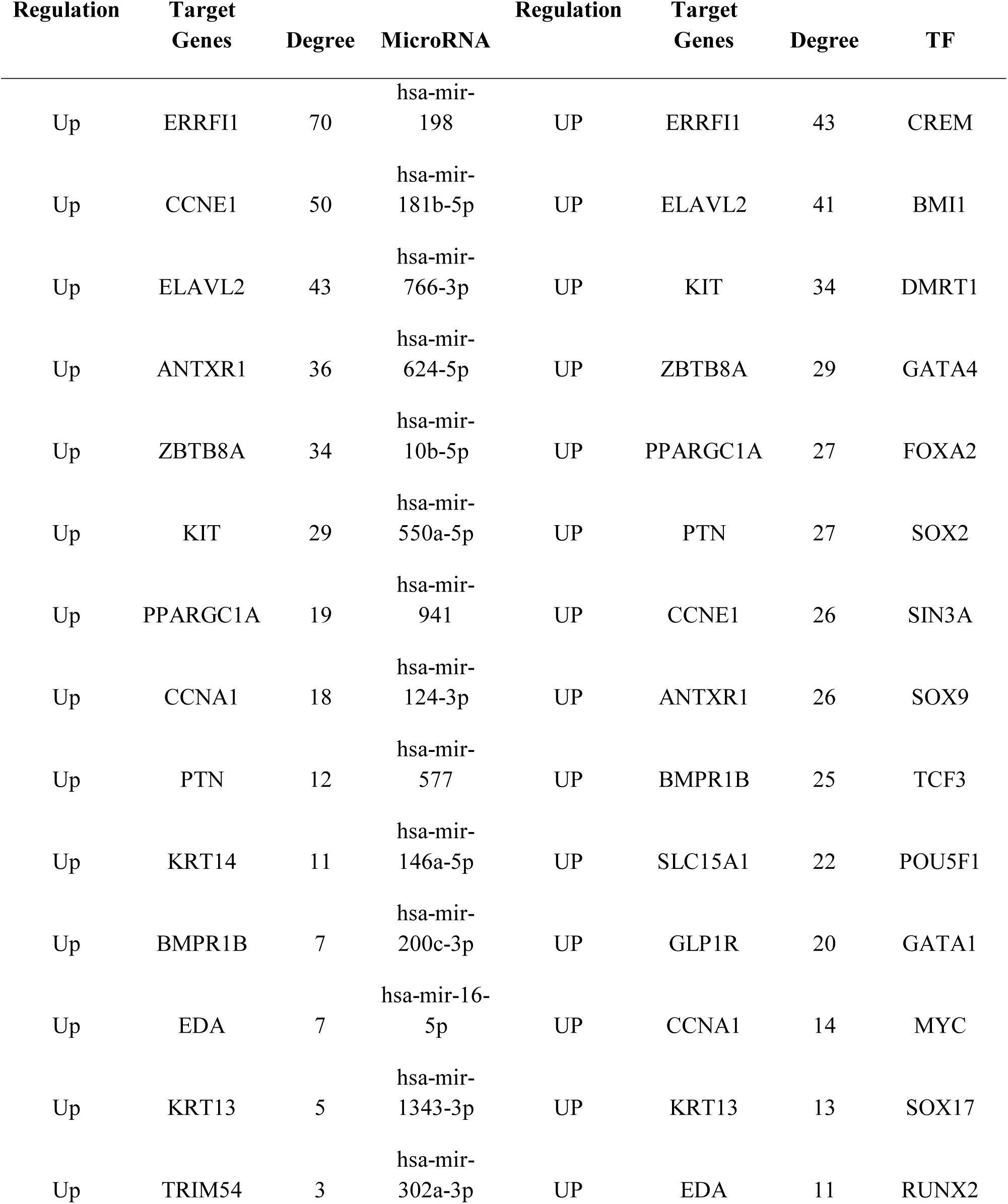
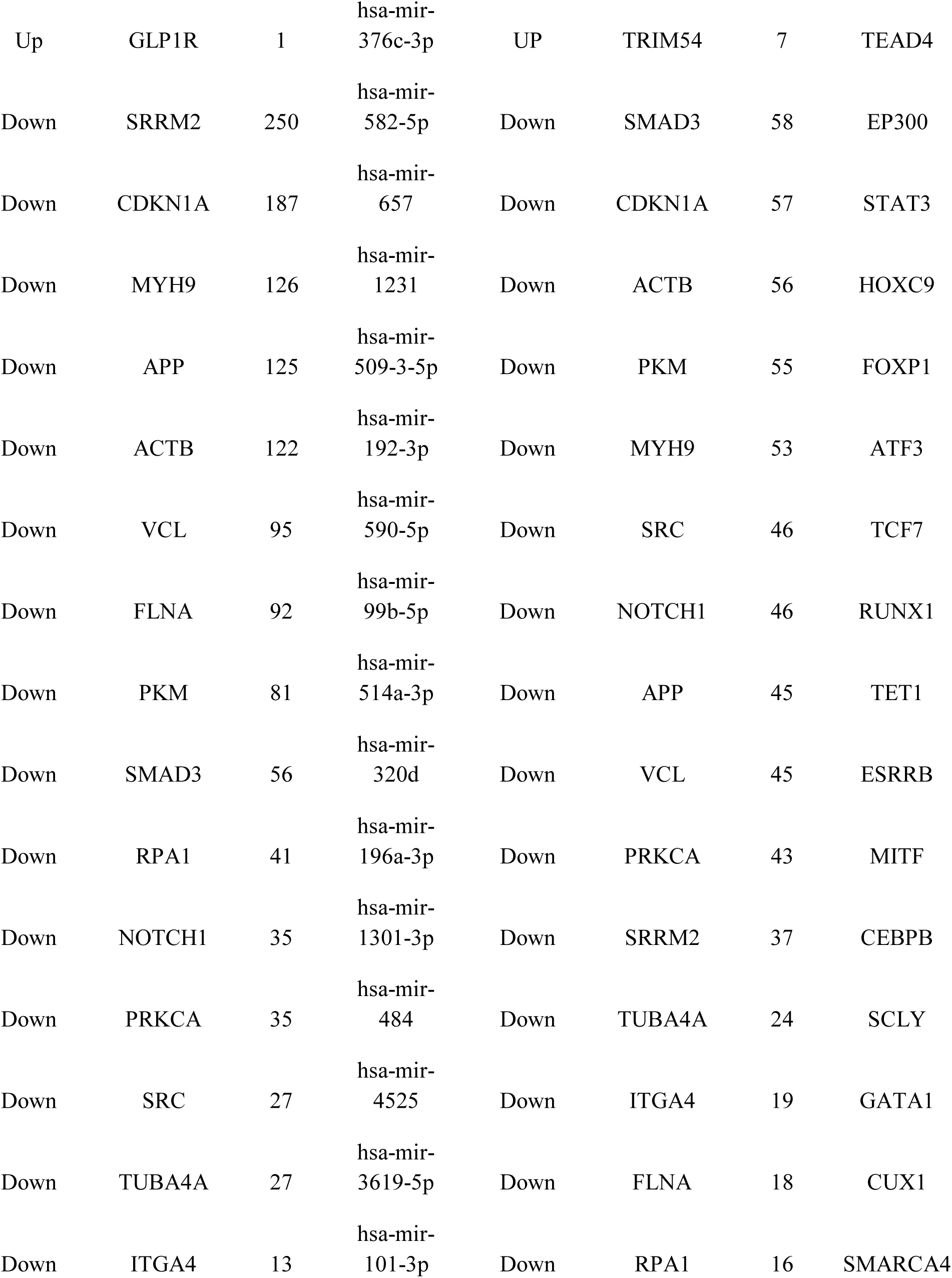
miRNA - target gene and TF - target gene interaction

| Regulation | Target Genes | Degree | MicroRNA | Regulation | Target Genes | Degree | TF |
| --- | --- | --- | --- | --- | --- | --- | --- |
| Up | ERRFI1 | 70 | hsa-mir-198 | UP | ERRFI1 | 43 | CREM |
| Up | CCNE1 | 50 | hsa-mir-181b-5p | UP | ELAVL2 | 41 | BMI1 |
| Up | ELAVL2 | 43 | hsa-mir-766-3p | UP | KIT | 34 | DMRT1 |
| Up | ANTXR1 | 36 | hsa-mir-624-5p | UP | ZBTB8A | 29 | GATA4 |
| Up | ZBTB8A | 34 | hsa-mir-10b-5p | UP | PPARGC1A | 27 | FOXA2 |
| Up | KIT | 29 | hsa-mir-550a-5p | UP | PTN | 27 | SOX2 |
| Up | PPARGC1A | 19 | hsa-mir-941 | UP | CCNE1 | 26 | SIN3A |
| Up | CCNA1 | 18 | hsa-mir-124-3p | UP | ANTXR1 | 26 | SOX9 |
| Up | PTN | 12 | hsa-mir-577 | UP | BMPR1B | 25 | TCF3 |
| Up | KRT14 | 11 | hsa-mir-146a-5p | UP | SLC15A1 | 22 | POU5F1 |
| Up | BMPR1B | 7 | hsa-mir-200c-3p | UP | GLP1R | 20 | GATA1 |
| Up | EDA | 7 | hsa-mir-16-5p | UP | CCNA1 | 14 | MYC |
| Up | KRT13 | 5 | hsa-mir-1343-3p | UP | KRT13 | 13 | SOX17 |
| Up | TRIM54 | 3 | hsa-mir-302a-3p | UP | EDA | 11 | RUNX2 |
| Up | GLP1R | 1 | hsa-mir-376c-3p | UP | TRIM54 | 7 | TEAD4 |
| Down | SRRM2 | 250 | hsa-mir-582-5p | Down | SMAD3 | 58 | EP300 |
| Down | CDKN1A | 187 | hsa-mir-657 | Down | CDKN1A | 57 | STAT3 |
| Down | MYH9 | 126 | hsa-mir-1231 | Down | ACTB | 56 | HOXC9 |
| Down | APP | 125 | hsa-mir-509-3-5p | Down | PKM | 55 | FOXP1 |
| Down | ACTB | 122 | hsa-mir-192-3p | Down | MYH9 | 53 | ATF3 |
| Down | VCL | 95 | hsa-mir-590-5p | Down | SRC | 46 | TCF7 |
| Down | FLNA | 92 | hsa-mir-99b-5p | Down | NOTCH1 | 46 | RUNX1 |
| Down | PKM | 81 | hsa-mir-514a-3p | Down | APP | 45 | TET1 |
| Down | SMAD3 | 56 | hsa-mir-320d | Down | VCL | 45 | ESRRB |
| Down | RPA1 | 41 | hsa-mir-196a-3p | Down | PRKCA | 43 | MITF |
| Down | NOTCH1 | 35 | hsa-mir-1301-3p | Down | SRRM2 | 37 | CEBPB |
| Down | PRKCA | 35 | hsa-mir-484 | Down | TUBA4A | 24 | SCLY |
| Down | SRC | 27 | hsa-mir-4525 | Down | ITGA4 | 19 | GATA1 |
| Down | TUBA4A | 27 | hsa-mir-3619-5p | Down | FLNA | 18 | CUX1 |
| Down | ITGA4 | 13 | hsa-mir-101-3p | Down | RPA1 | 16 | SMARCA4 |

### TF-hub gene regulatory network construction

To investigate interaction correlations between the hub genes and TFs, a TF-hub gene interaction network was constructed, which has 503 nodes (hub genes:309; miRNAs: 194) and 7686 edges. Results from the current investigation demonstrated that one target gene might be regulated by several TFs, (Fig. 6) and are listed in Table 5. Up regulated hub genes including, ERRFI1 targeted by 43 TFs (ex; CREM); ELAVL2 targeted by 41 TFs (ex; BMI1); KIT targeted by 34 TFs (ex; DMRT1); ZBTB8A targeted by 29 TFs (ex; GATA4); PPARGC1A targeted by 27 TFs (ex; FOXA2). Similarly, down regulated hub genes including, SMAD3 targeted by 58 TFs (ex; EP300); CDKN1A targeted by 57 TFs (ex; STAT3); ACTB targeted by 56 TFs (ex; HOXC9); PKM targeted by 55 TFs (ex; FOXP1); MYH9 targeted by 53 TFs (ex; ATF3).

**Fig. 6.**
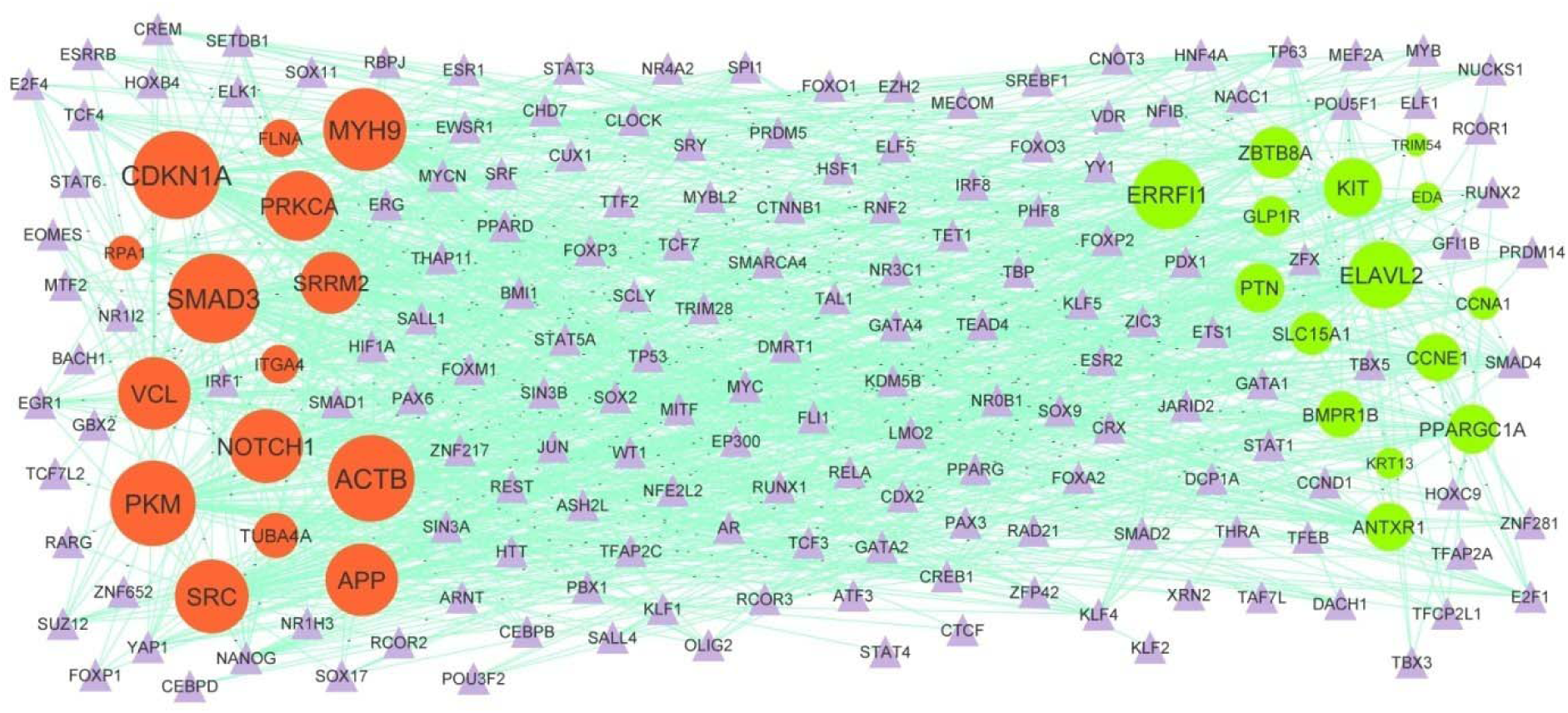
Target gene - TF regulatory network between target genes. The purple color triangle nodes represent the key TFs; up regulated genes are marked in green; down regulated genes are marked in red.

### Receiver operating characteristic curve (ROC) analysis

Based on the data of GSE154377, the patients were divided into the normal pregnancy group and GDM group. The ROC curves were analyzed by pROC package in R statistical software to evaluate the diagnostic accuracy of hub genes for GDM. As shown in Fig. 7, TRIM54, ELAVL2, PTN, KIT, BMPR1B, APP, SRC, ITGA4, RPA1 and ACTB achieved an AUC value of >0.90, demonstrating that these genes have high sensitivity and specificity for GDM diagnosis. Therefore, the ten hub genes identified, TRIM54, ELAVL2, PTN, KIT, BMPR1B, APP, SRC, ITGA4, RPA1 and ACTB might be considered as new biomarkers for GDM and require further investigation to validate these preliminary findings.

**Fig. 7.**
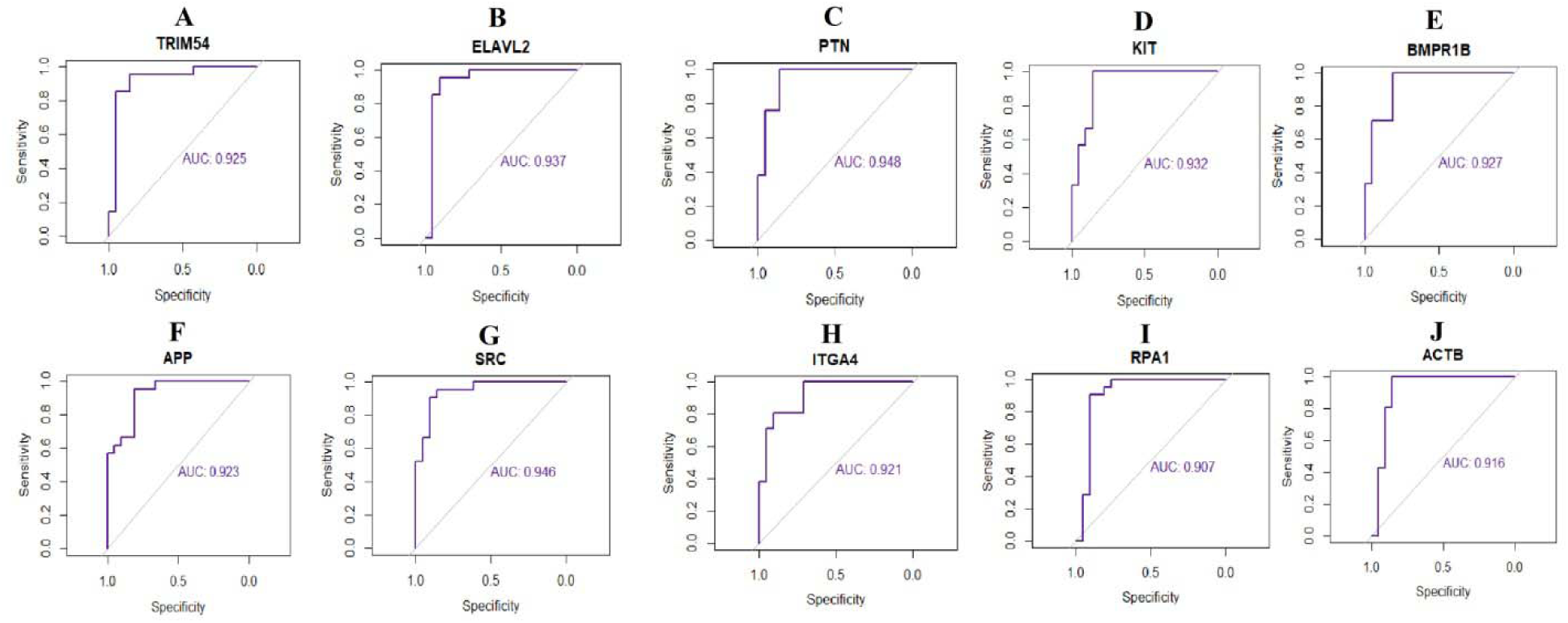
ROC curve analyses of hub genes. A) TRIM54 B) ELAVL2 C) PTN D) KIT E) BMPR1B F) APP G) SRC H) ITGA4 I) RPA1 J) ACTB

## Discussion

A number of pre-clinical and clinical investigations have been conducted to reveal the underlying mechanisms of GDM in the past decades; however, the incidence of GDM remains high. This is primarily due to the majority of the investigation focusing genetic risk factor [38]. In the current investigation, expression profiling by high throughput sequencing dataset was analyzed and 953 DEGs (478 up and 475 down regulated genes) were identified. Existing evidences have reported that HNF4G [39], SLC15A1 [40], CXCL5 [41] and MALAT1 [42] promotes obesity, but these genes might be novel target for GDM. NPHP3-AS1 [43], LTA (lymphotoxin alpha) [44], CXCL5 [45] and MALAT1 [46] are found to be expressed in cardiovascular diseases, but these genes might be novel targets for GDM. A recent study demonstrated that XIST (X inactive specific transcript) [47] and MALAT1 [48] serves a crucial role in GDM progression. LTA (lymphotoxin alpha) [49], CXCL5 [50] and MALAT1 [51] are associated with prognosis of diabetes mellitus, but these genes might be novel targets for GDM. LTA (lymphotoxin alpha) [52], CXCL5 [53], MALAT1 [54] and TMEM70 [55] have been demonstrated to function in hypertension, but these genes might be novel targets for GDM. CXCL5 has been shown to be activated abortion [56].

GO and REACTOME pathway enrichment analyses were used to investigate the interactions of DEGs. Pathways include hemostasis [57], neutrophil degranulation [58], immune system [59] and cytokine signaling in immune system [60] are responsible for progression of GDM. LGR5 [61], GREM1 [62], GLRA3 [63], NEUROD4 [64], CYP2J2 [65], KCNH6 [66], LBP (lipopolysaccharide binding protein) [67], CXCL14 [68], RGN (regucalcin) [69], NPY2R [70], SERPINB13 [71], WNT5A [72], EDA (ectodysplasin A) [73], HSD11B2 [74], ACVR1C [75], NEUROD1 [76], SLIT2 [77], PPARGC1A [78], IGF1 [79], OSR1 [80], CYP46A1 [81], TLR3 [82], BMP7 [83], SELP (selectin P) [84], HLA-A [85], NOTCH2 [86], LRP1 [87], CLU (clusterin) [88], FCN1 [89], CDKN1A [90], SMAD3 [91], HLA-E [92], PTPRC (protein tyrosine phosphatase receptor type C) [93], MYH9 [94], JAK3 [95], IL6R [96], TIMP1 [97], DOCK8 [98], TNFRSF1B [99], ITGAL (integrin subunit alpha L) [100], CD47 [101], RARA (retinoic acid receptor alpha) [102], DGKD (diacylglycerol kinase delta) [103], PLEK (pleckstrin) [104], PREX1 [105], BSCL2 [106], PANX1 [107], IRF7 [108], NOTCH1 [109], STIM1 [110], TRIM13 [111], LRBA (LPS responsive beige-like anchor protein) [112], CXCR4 [113], MDM4 [114], MYO9B [115] and PDE5A [116] were revealed to be expressed in diabetes mellitus, but these genes might be novel targets for GDM. SIX1 [117], GREM1 [118], GHRHR (growth hormone releasing hormone receptor) [119], GPR37L1 [120], CYP2J2 [121], AQP4 [122], ROS1 [123], LBP (lipopolysaccharide binding protein) [124], SGCD (sarcoglycan delta) [125], CXCL14 [126], RGN (regucalcin) [127], F9 [128], KCND2 [129], AZGP1 [130], HAS2 [131], CNTN4 [132], WNT5A [133], PTHLH [134], FAP (fibroblast activation protein alpha) [135], THSD7A [136], SHROOM3 [137], ETV1 [138], CYP24A1 [139], SLIT2 [140], GJC1 [141], PPARGC1A [142], TRPM3 [143], IGF1 [144], TRPV6 [145], TLR3 [146], BMP7 [147], DSG2 [148], POSTN (periostin) [149], ENOX1 [150], KCNA7 [151], ITGB3 [152], THBS1 [153], TRPC6 [154], SELP (selectin P) [155], VCL (vinculin) [156], P2RY1 [157], NOTCH2 [158], F2RL3 [159], PSAP (prosaposin) [160], LRP1 [161], CLU (clusterin) [162], FCN1 [163], CDKN1A [164], SMAD3 [165], CAPN1 [166], BAK1 [167], FERMT3 [168], HLA-E [169], ADA2 [170], PTPRC (protein tyrosine phosphatase receptor type C) [171], PEAR1 [172], TGFB1 [173], JAK3 [174], ALOX12 [175], ITGB2 [176], SELPLG (selectin P ligand) [177], TNFSF4 [178], BCL11B [179], IL17RA [180], CST3 [181], BCL3 [182], TNFRSF1B [183], SERPINA1 [184], CD47 [185], CD74 [186], STXBP2 [187], LEF1 [188], F13A1 [189], BSCL2 [190], VCAN (versican) [191], SUN2 [192], PPM1L [193], IRF7 [194], NOTCH1 [195], EHD3 [196], PCSK6 [197], STIM1 [198], CXCR4 [199], ATP2A2 [200], CALM3 [201], CSF1R [202], CLPP (caseinolytic mitochondrial matrix peptidase proteolytic subunit) [203], PIK3IP1 [204], CDK9 [205], USP22 [206], MDM4 [207], TUBB1 [208], PDE5A [209] and GNB5 [210] plays important roles in cardiovascular diseases progression, but these genes might be novel targets for GDM. Polonikov et al [211], Chida et al [212], Yamada et al [213], Zhang et al [214], Kawarazaki et al [215], Ueda et al [216], Bao et al [217], Leng et al [218], Taghvaei et al [219], Norling et al [220], Brown et al [221], Singh et al [222], Liu et al [223], Nie et al [224], Liu et al [225], Sharma et al [226], Liu et al [227], Jain et al [228], Novoyatleva et al [229], Yu et al [230], Sahoo et al [231], Gamboa et al [232], Zeng et al [233], Zabini et al [234], Olivi et al [235], Liu et al [236], Nabrdalik et al [237], Katayose et al [238], Topchieva et al [239], Onal et al [240], Speirs et al [241], Novelli et al [242], Le Hiress et al [243], Chang et al [244], Deng et al [245], Wang et al [246], Dhande et al [247], Yu et al [248] and Li et al [249] showed that CYP2J2, BMPR1B, ROS1, SCN7A, WNT5A, HSD11B2, CYP24A1, WNK3, PPARGC1A, IGF1, OSR1, TLR3, BMP7, POSTN (periostin), ITGB3, THBS1, FLNA (filamin A), TRPC6, SELP (selectin P), VCL (vinculin), NOTCH2, F2RL3, LRP1, CLU (clusterin), SMAD3, PEAR1, MYH9, TGFB1, PDGFA (platelet derived growth factor subunit A), IL6R, TIMP1, TNFRSF1B, CD47, CD74, VCAN (versican), IRF7, NOTCH1, STIM1, CXCR4 and PDE5A expression can be associated with hypertension progression, but these genes might be novel targets for GDM. ROS1 [250], WNT2 [251], PHLDA2 [252], WNT5A [253], HSD11B2 [254], CLDN1 [255], CYP24A1 [256], HRG (histidine rich glycoprotein) [257], TLR3 [258], THBS1 [259], TRPC6 [260], SELP (selectin P) [261], CLU (clusterin) [262], CDKN1A [263], ITGB1 [264], HLA-E [265], IL6R [266], TIMP1 [267], SERPINA1 [268], CD74 [269], PANX1 [270], STIM1 [271] and CXCR4 [272] were revealed to be correlated with disease outcome in patients with preeclampsia. Previously reported studies have shown that the expression of LBP (lipopolysaccharide binding protein) [273], MC1R [274], CXCL14 [275], RGN (regucalcin) [276], NPY2R [277], MFAP5 [278], WNT5A [279], EDA (ectodysplasin A) [280], THSD7A [281], NEUROD1 [282], SLIT2 [283], PPARGC1A [219], IGF1 [144], OSR1 [284], TLR3 [285], BMP7 [286], POSTN (periostin) [287], LRP1B [288], THBS1 [289], NOTCH2 [290]. LRP1 [291], CLU (clusterin) [292], SMAD3 [91], TGFB1 [293], APP (amyloid beta precursor protein) [294], ITGB2 [295], IL6R [296], TIMP1 [297], CD47 [298], CD74 [299], RARA (retinoic acid receptor alpha) [300], DOCK2 [301], F13A1 [302], IRF7 [303], STIM1 [304], CXCR4 [305], MGLL (monoglyceride lipase) [306], M6PR [307], USP22 [308] and CASP2 [309] were mainly involved in progression of obesity, but these genes might be novel targets for GDM. Recent studies have shown that GLP1R [310], NEUROD1 [311], PPARGC1A [312], IGF1 [313], LRP1B [314], NOTCH2 [315], BAK1 [316], TLR2 [317], IL6R [318], TIMP1 [319], PIK3CD [320], PRKCA (protein kinase C alpha) [321], CXCR4 [322], RAB8A [323] and M6PR [324] are associated with GDM. Abnormal expression PDE5A was linked with abortion [325].

Hub genes in the PPI network and modules are linked to the progression of GDM. TRIM54, ELAVL2, PTN (pleiotrophin), KIT (KIT proto-oncogene, receptor tyrosine kinase), SRC (SRC proto-oncogene, non-receptor tyrosine kinase), ITGA4, RPA1, ACTB (actin beta) and IGSF1 were might be the novel biomarkers for GDM.

At the same time, hub genes, miRNAs and TFs imlpicated with GDM, all of which were located in the core nodes in MiRNA-hub gene regulatory network and TF-hub gene regulatory network, meaning that these hub genes, miRNAs and TFs could be critical therapeutic targets to protect against GDM. hsa-mir-766-3p [326], hsa-mir-10b-5p [327], hsa-mir-657 [328], GATA4 [329], FOXA2 [330], STAT3 [331] and ATF3 [332] plays an important role in the progression of diabetes mellitus, but these miRNAs might be novel targets for GDM. hsa-mir-582-5p is closely related to obesity [333], but these miRNAs might be novel targets for GDM. Wang et al. [334] found that the altered expression of hsa-mir-657 in GDM. Improta Caria et al [335], Li et al. [336], Park et al. [337], Cheng et al. [338] and Davis, [339] demonstrated that hsa-mir-657, BMI1, GATA4, STAT3 and ATF3 played a pivotal role in the hypertension, but these miRNAs might be novel targets for GDM. Improta Caria et al [335], Patankar et al. [340], Bochkis et al. [341], Nishimura et al. [342], Su et al. [343] and Kim et al. [344] suggested that hsa-mir-657, GATA4, FOXA2, EP300, STAT3 and ATF3 could induce obesity, but these genes might be novel targets for GDM. Accumulating evidence has demonstrated that hsa-mir-657 [345], hsa-mir-1231 [346], CREM (cAMP responsive element modulator) [347], BMI1 [348], GATA4 [349], EP300 [350], STAT3 [351], FOXP1 [352] and ATF3 [353] appears to be constitutively activated in cardiovascular diseases, but these genes might be novel targets for GDM. Abnormal expression of FOXP1 was responsible for loss of pregnancy [354]. ERRFI1, CCNE1, ANTXR1, ZBTB8A, SRRM2, PKM (pyruvate kinase M1/2), hsa-mir-198, hsa-mir-181b-5p, hsa-mir-624-5p, hsa-mir-509-3-5p, hsa-mir-192-3p, DMRT1, HOXC9, were might be the novel biomarkers for GDM.

## Conclusions

The findings from this bioinformatics analysis investigations identified signaling pathwayes and the essential hub genes that might contribute to GDM. These findings improve our understanding of the molecular pathogenesis of GDM. Our investigation has key clinical importance for the early diagnosis and treatment, as well as the prevention, of GDM and provides effective targets for the treatment of GDM. However, further validation experiments are required to confirm the function of the identified hub genes linked with GDM.

## Acknowledgement

I thank Giorgia Del Vecchio, University of California Los Angeles, Los Angeles, USA, very much, the author who deposited their profiling by high throughput sequencing dataset GSE154377, into the public GEO database.

## Ethical approval

This article does not contain any studies with human participants or animals performed by any of the authors.

## Informed consent

No informed consent because this study does not contain human or animals participants.

## Availability of data and materials

The datasets supporting the conclusions of this article are available in the GEO (Gene Expression Omnibus) (https://www.ncbi.nlm.nih.gov/geo/) repository. [(GSE154377) https://www.ncbi.nlm.nih.gov/geo/query/acc.cgi?acc=GSE154377)]

## Consent for publication

Not applicable.

## Author Contributions

B. V. - Writing original draft, and review and editing

C. V. - Software and investigation

